# Antimicrobial resistance in dairy slurry tanks: a critical point for measurement and control

**DOI:** 10.1101/2022.02.22.481441

**Authors:** Michelle Baker, Alexander D Williams, Steven P.T. Hooton, Richard Helliwell, Elizabeth King, Thomas Dodsworth, Rosa María Baena-Nogueras, Andrew Warry, Catherine A. Ortori, Henry Todman, Charlotte J. Gray-Hammerton, Alexander C. W. Pritchard, Ethan Iles, Ryan Cook, Richard D. Emes, Michael A Jones, Theodore Kypraios, Helen West, David A Barrett, Stephen J Ramsden, Rachel L Gomes, Chris Hudson, Andrew D Millard, Sujatha Raman, Carol Morris, Christine E R Dodd, Jan-Ulrich Kreft, Jon L Hobman, Dov J Stekel

## Abstract

Waste from dairy production is one of the world’s largest sources of contamination from antimicrobial resistant bacteria (ARB) and genes (ARGs). However, studies to date do not provide necessary evidence to inform antimicrobial resistance (AMR) countermeasures. We undertook a detailed, interdisciplinary, longitudinal analysis of dairy slurry waste. The slurry contained a population of ARB and ARGs, with resistances to current, historical and never-used on-farm antibiotics; resistances were associated with Gram-negative and Gram-positive bacteria and mobile elements (IS*Ecp1*, Tn*916*, Tn*21*-family transposons). Modelling and experimental work suggested that these populations are in dynamic equilibrium, with microbial death balanced by fresh input. Consequently, storing slurry without further waste input for at least 60 days was predicted to reduce ARB spread onto land, with >99% reduction in cephalosporin resistant *Escherichia coli*. The model also indicated that for farms with low antibiotic use, further reductions are unlikely to reduce AMR further. We conclude that the slurry tank is a critical point for prevalence and control of AMR, and that measures to limit the spread of AMR from dairy waste should combine responsible antibiotic use, including low total quantity, avoidance of human critical antibiotics, and choosing antibiotics with shorter half-lives, coupled with appropriate slurry storage.

## Introduction

Antibiotics provided to food-producing animals account for 73% of global antibiotic sales (1), prompting concerns about the selection of antibiotic resistance bacteria (ARB) and genes (ARGs), and their migration from livestock and their environment to humans. ARB and ARGs associated with livestock can enter humans through consumption of animal products, e.g. contaminated meat (2, 3) and dairy (4, 5), or more indirectly, e.g. through land-application of animal waste, which may subsequently infiltrate crops (6, 7) and connected water resources (8, 9).

Cattle production comprises 50% of global Livestock Standard Units (10), so has considerable environmental impacts that need to be mitigated (11). There are approximately 265 million dairy cows globally, producing high volumes of waste manure, estimated at 3 billion tonnes per year (www.faostat.org). In the UK, the site of this study, dairy farms are estimated to account for 80% (67 million tonnes) of total annual livestock manure production (12), with more cattle waste material applied to soil in England and Wales than swine and poultry combined (13).

Antibiotics are routinely administered to dairy cattle for treatment, and, in some cases, prevention of common illnesses, including mastitis and respiratory disease (14–16). Lameness, the most costly disease to UK dairy farms (17), is often prevented with application of antimicrobial metals (copper, zinc) or other chemicals (formalin, glutaraldehyde) in the form of footbaths (18), known to co-select for antibiotic resistance (19, 20). Dairy waste can therefore contain selective and co-selective pressures in the form of mixtures of antibiotics and assorted antimicrobials, as well as ARB, including Extended Spectrum Cephalosporin-Resistant (ESC-R) *E. coli* (21, 22), and genetic resistance determinants (23, 24). Thus, dairy waste may represent one of the world’s most substantial routes for AMR to enter the environment, including onto fields and grasslands used for food production and into water ways.

To limit the risks of AMR, many countries have introduced responsible use policies, including reducing overall agricultural use of antibiotics (25), or of human critical antibiotics, including 3^rd^/4^th^ generation cephalosporins (26). However, antibiotics and other antimicrobials remain necessary for safeguarding animal health and welfare. Thus, other countermeasures are also needed to reduce the transmission or prevalence of ARB and ARGs from dairy waste into the environment. For example, current UK guidelines suggest that storage of solid manure and slurry without fresh input for three months can ameliorate AMR risk (27), but no evidence is provided. Slurry storage is essential in the UK and other countries where dairy cows are housed indoors for large parts of the year, and where slurry cannot be spread onto land that is frozen or deemed nitrate vulnerable. Two European studies have assessed storage effects on dairy manure, finding that certain ARGs increased during storage (28, 29); however, this ‘stored’ effluent regularly received fresh input. Contrastingly, a survey of several US dairy farms evaluating a different set of ARGs did not detect clear storage effects on ARG abundancesHurst, Oliver (30).

Other dairy waste studies took a ’snapshot in time’ (31–34), which does not allow for assessment of temporal stability of the resistome and the influence of storage. Factors such as temperature also influence the prevalence of enteric pathogens, indicator organisms and resistance phenotypes during manure storage (35–39). Meanwhile, studies assessing how cattle faecal resistomes respond to contrasting antibiotic management practices generally place emphasis on individual cattle (40–42), with different microbiomes, rather than the collective faecal output of the herd. Liquid-solid separation of manure may also influence the persistence of AMR (43).Therefore, there is a need for detailed longitudinal studies of AMR in dairy slurry and potential mitigations.

This study assessed three key questions about AMR in slurry and its relationship to antibiotic use and slurry storage: (1) does the slurry tank select for or against AMR; (2) how does the resistance content of the slurry tank relate to altered patterns of farm antibiotic use; and (3) can slurry storage help reduce AMR in slurry before application to land? Our interdisciplinary approach combined phenotypic, genomic, and metagenomic microbiological analyses with chemical analyses, antibiotic use records and predictive mathematical models, to provide a temporal evaluation of slurry tank content over six months. This was supplemented by concurrent mini-slurry tank experiments which facilitated the controlled study of isolated slurry. We designed the mathematical model to enable us to study the impact of farm practices that would be impractical or unethical to perform through purely empirical approaches. These included major changes to farm slurry handling, antibiotic reduction to a level that would threaten animal welfare, or the reintroduction of use of human critical 3^rd^ or 4^th^ generation cephalosporins. Thus, this study enables the identification of approaches to reduce the spread of AMR into the environment from an important source of such contamination.

## Methods

### Sample site

We surveyed a mid-sized, high performance commercial dairy farm in England, housing ∼200 milking Holstein Friesian cattle at the time of study. Practice at this farm is typical of management methods at high-performance dairy farms, although all farms vary. Milking cattle are housed indoors on concrete, and all excreta are regularly removed from cattle yards by automatic scrapers into a drainage system terminating at the 3000m^3^ slurry tank. The drainage system also receives used cleaning materials and wash water, used footbath containing zinc and copper, waste milk from cows treated with antibiotics, and rainwater runoff. An automated screw press (Bauer S655 slurry separator with sieve size 0.75 mm; Bauer GmbH, Voitsberg, Austria) performs liquid-solid separation prior to the slurry tank. Liquids enter the slurry tank semi-continuously, while solids are removed to a muck heap. Calves, dry cows, and heifers are housed separately from the milking cows. Faeces and urine from calves drain into the common drainage system, whilst dirty straw from calf housing is taken directly to the muck heap. Excess slurry can be pumped to an 8000m^3^ lagoon for long term storage. Slurry from either the slurry tank or lagoon is used to fertilise grassland and arable fields.

### Microbiological sampling, strain isolation, antimicrobial susceptibility testing and whole genome sequencing

Liquid samples were collected from the slurry tank on 17 dates between May and November 2017 (Table S1). *Escherichia coli* strains were isolated using Tryptone Bile X-Glucuronide (TBX) or MacConkey agar or TBX/MacConkey supplemented with 16 µg ml^-1^ ampicillin (AMP), or 2 µg ml^-1^ cefotaxime (CTX); or on CHROMagar ESBL™ agar. Putative *E. coli* isolates were subcultured onto TBX agar or TBX agar supplemented with 2 µg ml^-1^ CTX. *E. coli* strains were confirmed using oxidase (22) and catalase tests. Antimicrobial susceptibility testing (AST) using a range of antibiotic discs (Table S2) was carried out on 811 *E. coli* isolates in accordance with CLSI (44) guidelines. ESC-R *E. coli* strains were identified by phenotypic resistance profile as putatively *ampC* or CTX type, and confirmed by PCR (22). Presence of Tn*21*-like mercury resistance transposons within the *E. coli* isolates was initially screened for by growing isolates on LB agar containing 25 µg ml^-1^ HgCl_2_. Their presence was confirmed by PCR (45). Genome assembly of selected ESC-R and mercury resistant *E. coli* strains using PacBio, was carried out by the Centre for Genomic Research (CGR), University of Liverpool, with methods for library preparation and sequencing as previously described (46) or by Illumina short read WGS by MicrobesNG (University of Birmingham, UK). Genome sequence analysis and annotation was conducted using Prokka (47), CSARweb (48), Snapgene viewer (Insightful Science; snapgene.com), Res Finder (49) and Plasmid Finder (50). Genome sequences are deposited with NCBI under BioProject PRJNA736866.

### Metagenomics Sample collection and DNA extraction

#### Main tank Sample Collection

Samples were collected from the slurry tank monthly between June and October 2017, using a clean stainless steel bucket, and aliquoted into 2 large glass bottles with external PE protection. Three replicate extractions were performed on 250 μl of each sample using a PowerFecal Kit (Qiagen), according to manufacturer’s instructions (15 extractions in total). DNA was quantified using a Qubit fluorometer (Invitrogen) while quality was assessed via Nanodrop 1000 (ThermoFisher). Extracted DNA was stored at 4°C pending sequencing.

#### Mini-Tank Experiments

Miniaturised experimental slurry tanks were set up to assess the impact of storing slurry (control tanks) and to measure antibiotic stability. Twelve mini-tanks were situated on the farm from 23/4/2018 to 15/6/2018 at ambient temperature (mean 24 hour temperature in liquid ranged between 7° to 17°), protected from rain and direct sunlight, and containing 10L grab samples of slurry from the surface of the main slurry tank. Six different conditions were tested in duplicate (all amounts per litre): control; + SSD (0.2mL of slurry solids homogenised by stomacher, including 67 CFU of CTX-resistant *E. coli*); + SSD + 3μg cefquinome weekly addition; + SSD + 40μg cefalexin weekly addition; + SSD + 16.8g of footbath mix (Cu + Zn); + SSD + footbath + cefquinome). Mini-tanks were sampled four times (0, 2, 4 and 7 weeks after initial filling). Experimental conditions were mainly used for model calibration (Supplementary Text 1). *E. coli* were isolated and cultured as described above except MacConkey agar was not used. DNA was extracted and processed for sequencing as above. Antibiotic concentrations were measured as described previously Baena-Nogueras, Ortori (51) with further methods described in Supplementary Text 3.

#### Metagenomic Sequencing, Assembly and Analysis

Metagenomic sequencing of DNA extracted from the main slurry tank was performed by Liverpool Genomics using the Illumina HiSeq platform, and from the mini-slurry tanks by Edinburgh Genomics using the Illumina NovaSeq platform (150 bp paired end libraries in both cases). For the main tank, reads were trimmed with Cutadapt v1.2.1 (52) and Sickle v1.2.0.0 (53), while mini-tank reads were trimmed with Fastp v0.19.07 (54). Assembly was performed on trimmed reads using Megahit v1.1.3 (55). Main tank technical replicates were pooled by date and assembled using the settings: k-step 10; k-range 27-87. Mini-tank metagenomes were assembled individually (k-step ∼20, k-range: 21-99). Metagenome sequences are deposited with the ENA under Study Accession PRJEB38990.

Read-based searches for ARGs were performed with DeepARG v2 (56). ARGs were also identified on contigs (>1.5 kb length) in order to investigate the wider genetic context of the core resistome using ABRicate v1.0.1 (57), using MegaRes 2.0 for ARGs and metal resistance genes (MRGs) (58) (including experimentally verified MRGs; genes requiring SNP validation were excluded) and ACLAME 0.4 for MGEs (59). All data were analysed with stringencies: >60% gene coverage, >80% identity(60). Lastly, the BacMet2 database (61) was screened against translated peptides (based on Prodigal (62) output) from meta-assemblies of the main and mini-tanks (stringencies: >60% sequence identity and match length >50% of peptide length).

Taxonomic assignment of reads was performed using Kaiju v1.6.2 (63), with default settings. The reference database used was a microbial subset of the NCBI database (64), including additional fungal and other microbial eukaryotic peptide sequences. Contigs of interest were assigned putative identities using NCBI-nucleotide BLAST (65) (MegaBlast(66), highly similar sequences).

For both ARG and taxonomic assignments, statistical comparisons were carried out using the DirtyGenes likelihood ratio test (67), using randomized resampling (n=1000) from the null distribution to establish p-values.

### Water Quality Analysis

Water quality analysis was performed on the same samples as for microbiological culturing. For each sample, 2.5L was initially sampled. Probes were used to assess the pH (Hach PHC201), dissolved oxygen (Hach LDO101) and NaCl (Hach). The probe tip was rinsed in Milli-Q water (Merck), dabbed dry and submerged into the bottle containing slurry and left to equilibrate. The sample was then homogenized by shaking vigorously before decanting into a 250mL bottle for analysis using a Hach DR3900 Laboratory Spectrophotometer with cuvette test kits: sulphate (LCK153); ammonium (LCK303); chloride (LCK311); copper (LCK329); LATON total nitrogen (LCK338); nitrate (LCK340); nitrite (LCK342); phosphate (LCK348); zinc (LCK360); COD (LCK514); and TOC (LCK381). Standard procedures are available from https://uk.hach.com.

### Mathematical Model

A mechanistic, multi-strain model of AMR in the slurry tank was constructed to simulate a range of relevant farm management scenarios that would have been impractical or unethical to carry out empirically. In brief, it is a coupled ordinary differential equation model of bacterial populations including logistic growth, death (baseline and antimicrobial induced), horizontal transfer and fitness cost of resistance, inflow and outflow (68, 69). The model considered mobile resistance to penicillin, tetracycline, cephalexin, cefquinome, copper, and zinc, and was simulated for a full year in order to capture the recorded input of cephalexin and other antibiotics. The choice of resistances reflects our interests in ESC-R *E. coli* strains, and the risk of environmental contamination by mobile genes following slurry spreading. Full model description is provided in Supplementary Text 1, equations in Supplementary Text 2 and parameter values in Table S4. This model was deposited in BioModels (70) as MODEL1909100001. The secondary storage model is derived from this model by duplicating equations for each storage vessel (70) and also deposited as MODEL1909120002. A reduced model was used for parameter inference from mini-tank data. Model simulations were carried out in Matlab using the ode45 solver.

## Results

### XXX

#### Resistance to antibiotics with historic, current and no documented farm use

The majority of antibiotics administered to milking cows during the sampling period were aminocoumarins, aminoglycosides and beta-lactams delivered in combination, and beta-lactams and tetracyclines administered individually (Table S3). The last recorded use of sulphonamides (sulfadoxine) was in June 2016; of first generation cephalosporins (cephalexin) was in April 2017 (shortly before the start of the sampling period); of third generation cephalosporins (ceftiofur) was in January 2016; and of fourth generation cephalosporins (cefquinome) was in August 2015. Residual antibiotics or ARB associated with historical use could potentially be present in sludge at the bottom of the tank that cannot be piped for spreading. Smaller quantities of antibiotics are also given to youngstock; their waste does not enter the slurry system.

The dominant resistance phenotypes of cultured *E. coli* isolates from the slurry tank (Figure 1a) were ampicillin (34.6%), cefpodoxime (39.3%), cefotaxime (29.6%) and streptomycin (26.5%); other common phenotypes included tetracycline (13.6%), chloramphenicol (10.7%) and nalidixic acid (9.6%). Multidrug resistant *E. coli* strains (≥3 different antibiotic classes, Magiorakos, Srinivasan (71)) represented 37% of the cultured isolates (Figure 1b), detected in strains isolated on both antibiotic-supplemented and non-supplemented media. Of these isolates, 12 cefotaxime resistant *E. coli* strains were sequenced to characterize the resistance genes and mobile elements carrying them. Three carried IS*Ecp1* CTX-M-15, additionally carrying *qnrS* and *tetM* within the *ISEcp1* element. The other sequenced ESC-R strains were chromosomal *ampC* mutants.

**Figure 1:**
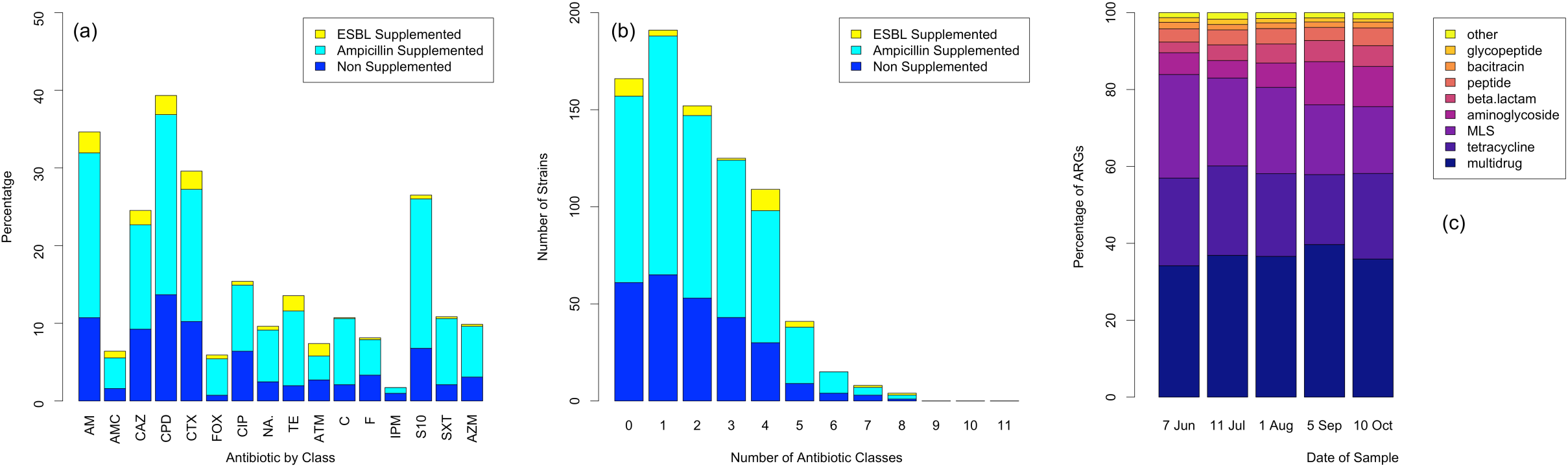
Antimicrobial resistance phenotypes and reads in the slurry tank. (a) Resistances to a panel of 16 antibiotics (Supplementary Table 2) largely do not depend on the type of supplemented media used. (b) The number of resistances per isolates; 37% of cultured isolates resistant to three or more antibiotic classes. These resistances are seen on all types of media. (c) Proportion of ARGs mapped to different antibiotic resistance classes (% reads). The metagenomic resistance profile is largely stable over time. There appears to be a gradual increase in the proportion of aminoglycoside and beta lactam resistance genes, which could be seen as consistent with antibiotic use during that period, but there is no statistical significance to the changes in proportions. ARGs are also reasonably consistent with observed phenotype data.

In main slurry tank metagenomes, eight resistance classes account for 98% of the ARGs identified in reads (Figure 1c): multidrug resistance genes (36.7%); tetracycline resistance genes (21.6%); macrolide-lincosamide-streptogramin (MLS) resistance genes (21.4%); aminoglycosides (7.3%); beta lactams (4.5%); peptides (4.0%); bacitracins (1.6%) and glycopeptides (1.2%). MRGs were also identified (*mer*: mercury; *cop*, *cus*, *pco/sil*: copper, copper/silver; *cad*, *czc:* cadmium, cadmium/zinc/cobalt; *ars*, arsenic/antimony; *pbr* lead resistance). In equivalent metagenome read assemblies, MLS and tetracycline ARGs were most frequently detected (70 and 46 contigs, respectively). Few MRGs were detected in main tank metagenome assemblies, limited to TCR copper resistance genes (5 contigs).

Overall, the identification of aminoglycoside, beta-lactam (excepting 3^rd^/4^th^ generation cephalosporins) and tetracycline resistance genes and phenotypes reflect current or recent farm antibiotic use, while the presence of zinc and copper resistance genes reflect transition metal use. The presence of sulphonamide and cephalosporin resistance genes and phenotypes may be due to historical use, or reflect widespread environmental occurrence (72). The prevalence of MLS resistance genes is unlikely to be associated with antibiotic use, as there is no recorded MLS use for milking cows.

#### Slurry tank properties and AMR remained stable due to frequent inputs

Water quality measures were largely stable (Figure 2a), with some fluctuations in July and August likely to be associated with mixing of slurry in the tank prior to spreading on fields. The relative contribution of the dominant drug-resistance categories listed above remained unchanged throughout the sampling period (Figure 1c; *p*=0.172, DirtyGenes test). Likewise, taxonomic analyses of read data showed the time-stable dominance of six bacterial phyla with at least 1% prevalence (Figure 2b; p=0.254, DirtyGenes test): Bacteroidetes (13.8%), Firmicutes (13.7%), Proteobacteria (4.7%), Spirochaetes (2.9%), Euryarcheaota (1.9%) and Tenericutes (1.4%). These phyla only account for 38% of the microbial community: there is considerable diversity in the tank with 178 phyla identified (Table S4).

**Figure 2:**
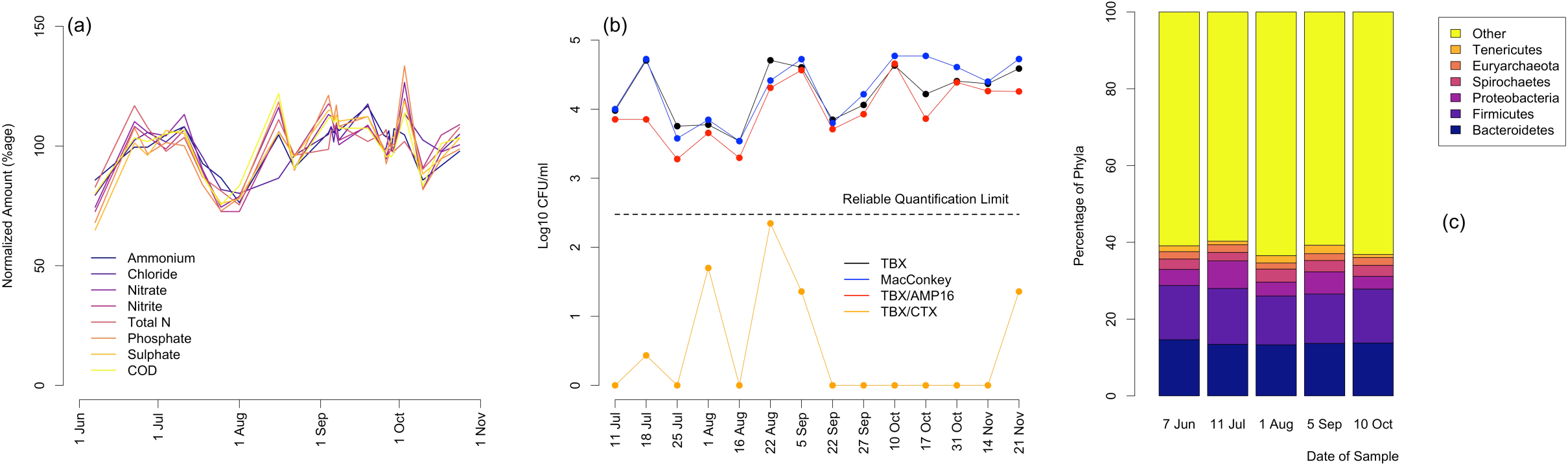
Stability of microbial ecosystem, *E. coli* counts and water quality measures. (a) water quality analysis from samples taken from the slurry tank over a five month period concurrent with microbial counts. Water quality measures are generally stable, with some fluctuations concordant with slurry use. (b) Six taxonomic groups accounting for at least 1% each of microbial reads show stable abundance in time. There is considerable diversity; these groups only account for 38% of reads, with all reads mapped to 178 different microbial phyla. (c) Counts of *E. coli* concentrations showing *E. coli* on TBX and MaConkey plates (all *E. coli*), TBX and AMP plates (*E. coli* resistant to ampicillin) and on CTX plates (ESC-R *E. coli*). Overall *E. coli* abundance is stable throughout the sampling period, as are counts of ampicillin resistant strains. CTX-resistant *E. coli* are only observed on five sampling days, and on all of those occasions at levels too low to be reliably quantified. The data for the other days are below the limit of detection of the method used and are plotted at 0 for ease of display.

The overall numbers of *E. coli* identified through culture-based enumeration also remained stable (Figure 2c), with concentrations of 4.23±0.40 (Log_10_ CFU mL^-1^) on TBX plates and 4.29±0.46 (Log_10_ CFU mL^-1^) on MacConkey media. *E. coli* strains resistant to ampicillin (TBX/Amp 16 µg mL^-1^) were stable at concentrations of 3.99±0.43 (Log_10_ CFU mL^-1^), i.e. ∼58% of cultured *E. coli* strains. *E. coli* that could be cultured on cefotaxime selective plates (TBX/CTX 2 µg L^-1^) were detected on only five of 17 sampling dates, with counts below 10 colonies per plate on all but one day (22^nd^ August). Thus, cefotaxime resistant *E. coli* were present at low levels, but could not be reliably quantified. The full AST profiles of the 811 isolates also show consistency over time, with some random variation, both on antibiotic-free and antibiotic-supplemented media (Figure 3).

**Figure 3:**
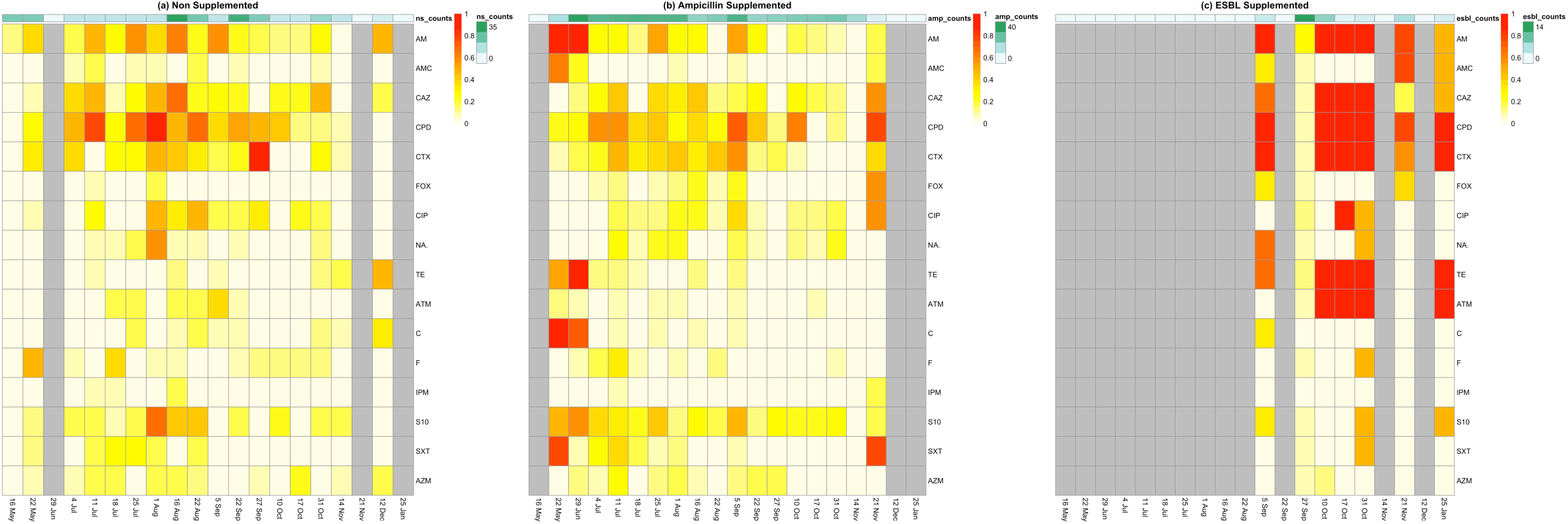
Antibiotic susceptibility testing of *E. coli* isolates shows diverse but stable range of phenotypic resistances. In each panel, the heatmap shows the proportion of strains resistant to each of 16 different antibiotics on each of the sampling dates. Grey bars indicate no use of those plate types on those dates. (a) plates without antibiotic supplement; (b) plates supplemented with ampicillin; (c) plates supplemented with cephalosporins. In all cases, the patterns of resistances are stable in time. Cephalosporin supplemented plates identify more resistant strains than other plates, including to other antibiotic classes, including tetracyclines and quinolones.

### Model predictions are consistent with microbial data

In the mathematical model, predicted resistance to penicillins fluctuated between 0.4% and 6.4% and cephalosporins between 0.5% and 7.9% (Figure 4a), i.e. both present but low, despite frequent inflow of antibiotics into the tank (Figure 4b). Resistance to tetracycline increased from low initial levels to fluctuate around ∼25% of the *E. coli* population (Figure 4a), before slowly declining over the longer term, reflecting the decline in tetracycline use later in the year. These predicted levels of tetracycline and cephalosporin resistances are concordant with the empirical phenotype above. Penicillin resistance in the model is lower than observed empirically, probably because resistance in the model is plasmid-borne, while many strains have chromosomal mutations of *ampC* or chromosomally located resistance genes that could be mobilised (e.g. IS*Ecp1*CTX-M-15 elements). The model predicts that zinc resistance is highly prevalent, rising to fluctuate around 80%, with co-occurrence of tetracycline and zinc resistance, typically fluctuating between 10 and 15%, consistent with predictions that the metal concentrations in the tank are co-selective (69).

**Figure 4:**
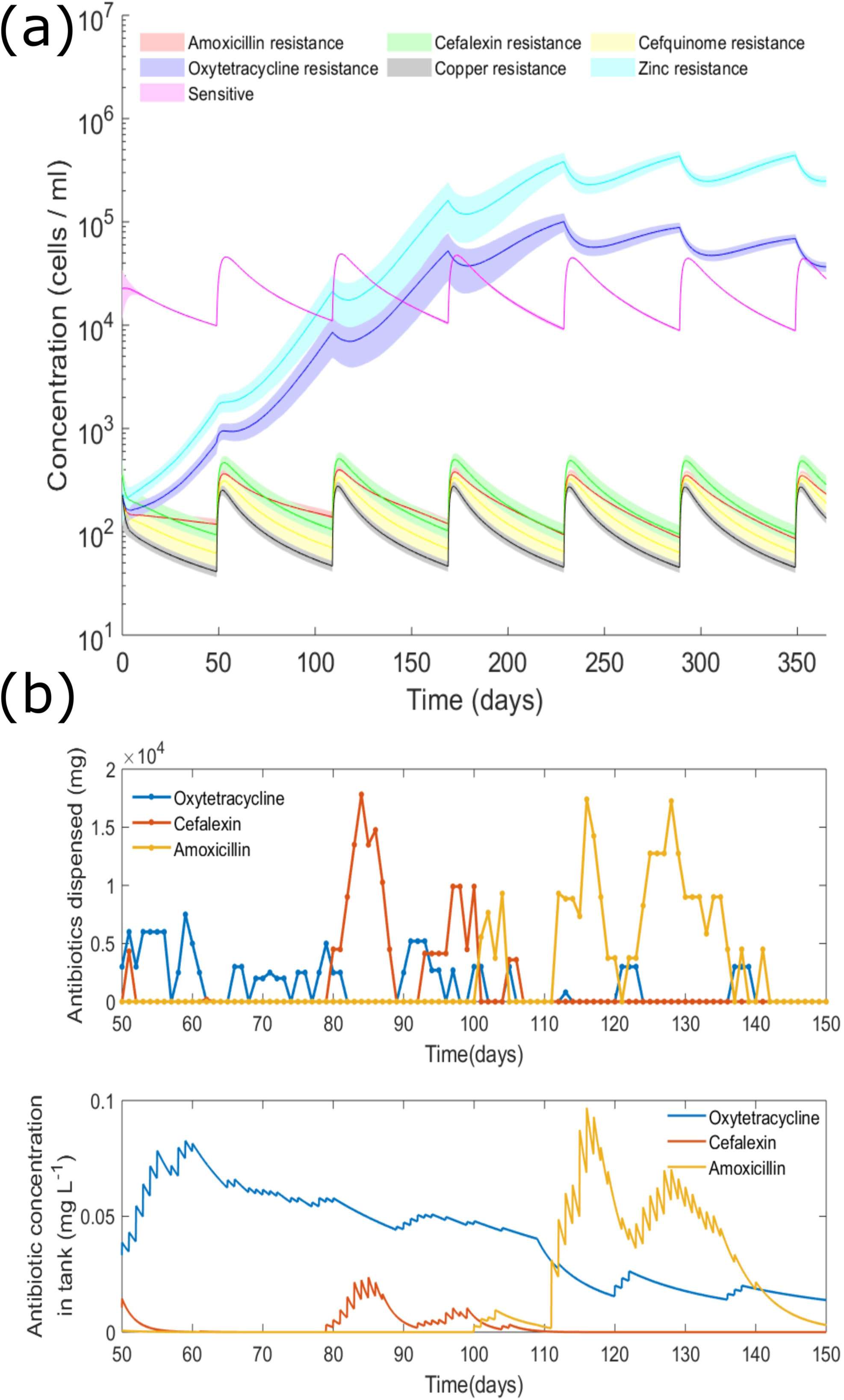
Model simulations of antimicrobials and antimicrobial resistance in the slurry tank. (a) Model prediction of resistant *E. coli* populations in the slurry tank over a year’s period, given (b) antibiotic usage on farm in 2017. Resistance groups are not mutually exclusive. The resistances are reasonably stable once the model simulation reaches its steady state, with fluctuations resulting from periodic removal of slurry for use as fertilizer. (b) Mass (in mg) of oxytetracycline, cefalexin and amoxicillin given during 2017 together with model simulation predicting concentrations (in mg L^-1^) of these antibiotics in the slurry tank over the same period. Observe that tetracycline is present in the tank, despite intermittent use, due to its high environmental stability. This explains the consistent proportion of tetracycline resistance. The two beta lactam antibiotics decay more rapidly after use.

### Associations of ARGs with other ARGs, integrons and Gram-positive taxa

Several metagenome contigs contained two or more ARGs, MRGs or associations with MGE markers in both the main tank (37 contigs) and mini-tank metagenome assemblies (101 contigs) (Figures S1 and S2). These include ARGs belonging to the same resistance gene group, e.g. *aph*3 and *aph*6 (both aminoglycoside resistance genes; Figure S3a) which were co-localised on five main-tank and eight mini-tank contigs; as well as genes associated with entirely different antibiotic resistance classes, e.g. *ant*6 and *tet*44 (aminoglycoside and tetracycline resistance, respectively; Figure S3b) were co-localised on two main-tank and eight mini-tank contigs. In other mini-tank contigs, *aph*3-*aph*6 were additionally co-resident with either a sulphonamide (*sul*2, 1 contig) or tetracycline (*tetY*, 1 contig) resistance gene. *tetM* was embedded within the widely documented Tn*916* transposon (18 *tetM* contigs in total, nine of which were linked with Tn*916* elements). The two largest Tn*916*-like contigs (18.3-18.9 kb) appear to be carried within Gram-positive bacteria, possibly *Streptococcus* spp. or *Enterococcus* spp. (NCBI-BLAST, ∼99.96% identity, ∼91% query coverage; Figure S3c). Furthermore, 21.4% (n= 6/28) of main and mini-tank contigs containing *cfxA* (class-A beta-lactamase) were co-localised with mobile elements.

Further identification of mobile resistance cassettes was through a screen of all *E. coli* strains for phenotypic mercury resistance as a marker for Tn*21* carriage. Sequence analysis of mercury resistant *E. coli* strains showed that three carried Tn*21* variants carrying the integron intI*2* conferring co-occurrence of combinations of penicillin, sulphonamide, aminoglycoside and quaternary ammonium compound resistances.

### Waste management for AMR reduction

We investigated the use of slurry storage to ameliorate resistance through a combination of empirical and modelling work. In the mini-tanks, we found that storage of slurry without inflow rapidly decreased the total concentration of cultured *E. coli* cells (Figure S6a), as well as *Escherichia, Pseudomonas and Klebsiella* spp. sequences identified by metagenomics (Figure 5). Reads assigned to gut-associated anaerobes belonging to Bacteroidetes including *Bacteroides* spp., *Alistipes* spp. and *Prevotella* spp. declined in steps. In contrast, the relative abundance of *Acinetobacter* spp. gradually increased until week four, before declining again by the end of the experiment (Figure 5).

**Figure 5:**
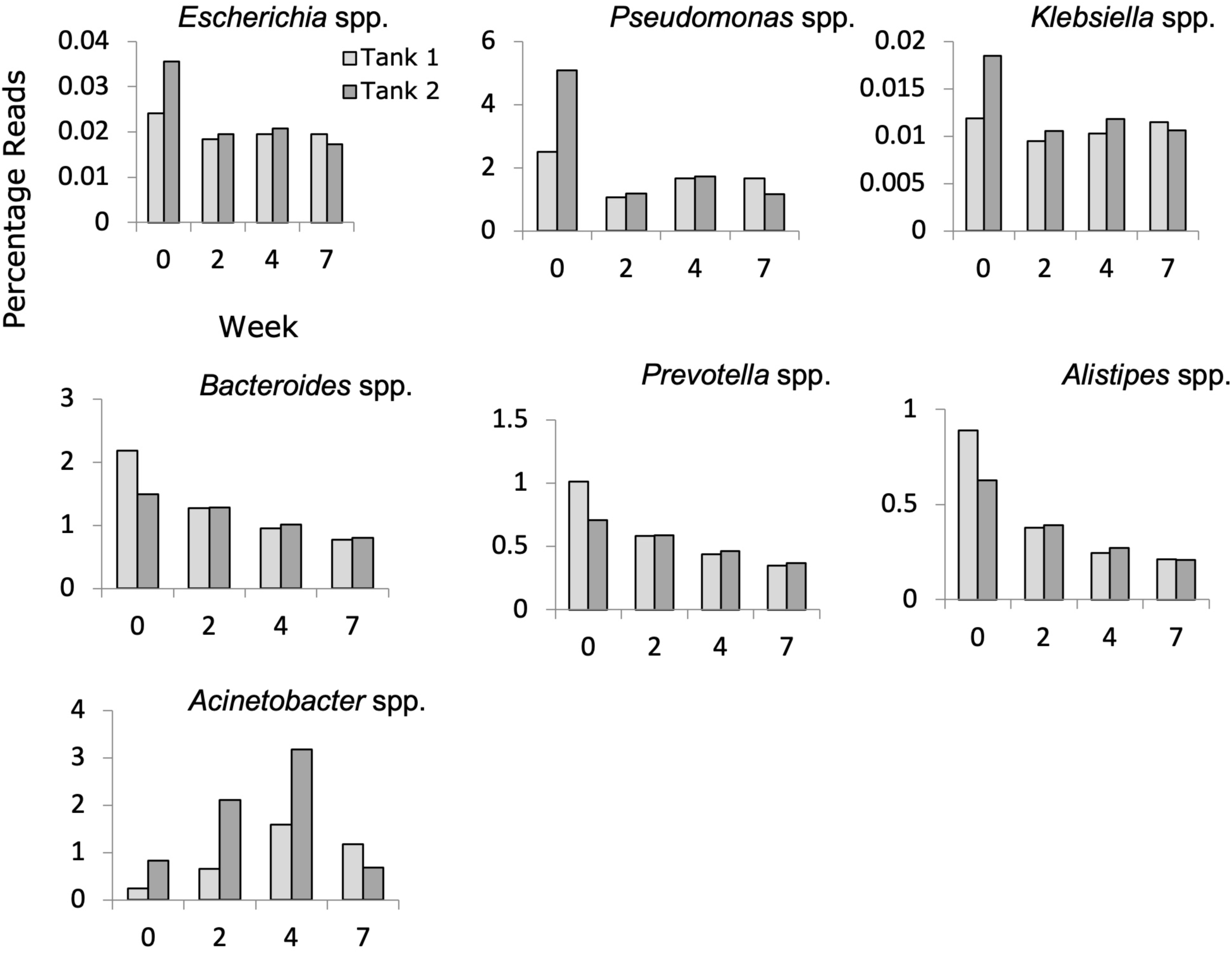
Storage without further waste-addition leads to a decline in select bacteria. Relative abundances of *Escherichia* spp., *Pseudomonas* spp., *Klebsiella* spp., *Bacteroides* spp., *Prevotella* spp. and *Alistipes* spp. in stored slurry based on metagenomic short-read data. *Escherichia* reads from metagenomics are concordant with culturing data (viable *E. coli* counts in CFU/ml over time are given in Figure S6a), both showing a stepwise decline. *Pseudomonas* and *Klebsiella* also show a stepwise decline. *Bacteroides*, *Prevotella*, and *Alistipes* show a gradual decline. *Acinetobacter* increase over the first four weeks before declining.

The prevalence of beta-lactam resistance genes declined considerably in <2 weeks (Figure 6a). The overall relative abundance of tetracycline resistance genes declined marginally over 7-weeks of storage (Figure 6b); however, different patterns were observed with different gene groups: *tetY* (Figure 6c) and *tet*40 (Figure 6d) declined sharply within two weeks, while others, e.g*. tetM* (Figure 6e) were maintained in stored slurry. According to BLAST analysis against the NCBI database, mini-tank contigs containing *tetY* (2 contigs) were likely associated with Gamma-Proteobacteria, while *tet40* (6 contigs) was consistently linked to Firmicutes. Similarly, *tetM* was typically associated with Firmicutes (7 of 16 contigs; >89% sequence coverage, >99% sequence identity), more specifically Bacilli. The proportion of MLS ARGs remained comparatively stable throughout (Figure 6f), consistent with their presence not connected with patterns of MLS use on the farm.

**Figure 6:**
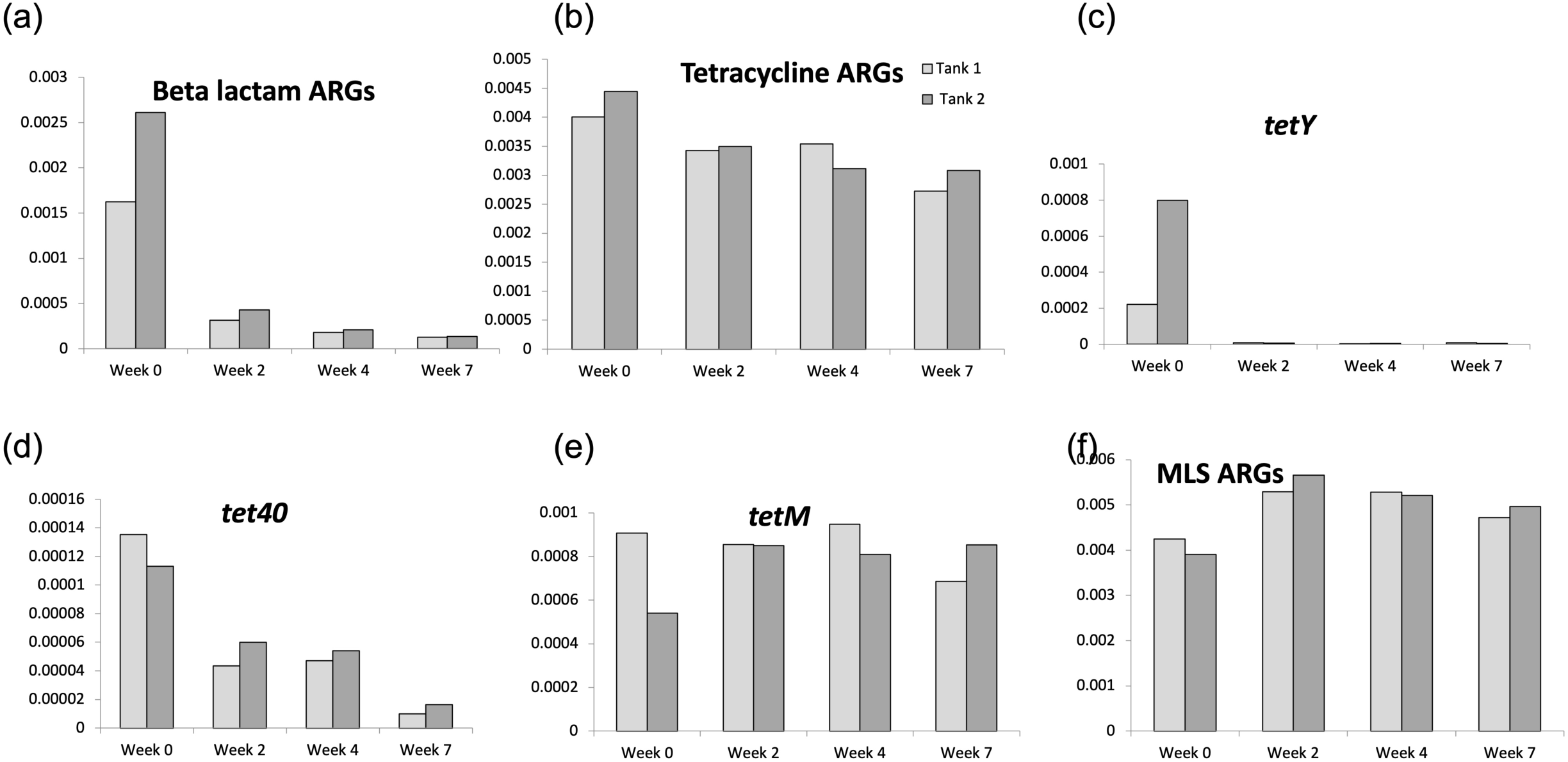
Impact of mini tank storage on selected ARGs based on DeepARG analysis. Relative abundance (percentage of reads) of (a) beta lactam ARGs; (b) tetracycline ARGs; (c) *tetY*; (d) *tet40*; (e) *tetM*; (f) MLS ARGs. The decline in beta lactam reads is consistent with other data. Tetracycline ARGs show different patterns for different genes. The persistence of MLS ARGs is consistent with their presence not related to lack of MLS use on the farm.

We implemented a two-stage in series storage mathematical model to consider whether the storage of slurry in the main tank, without fresh inputs, would reduce AMR in slurry prior to land application. The model predicted that after only four days of storage, 50% of the amoxicillin- and cefalexin-resistant *E. coli* are removed, and after 60 days of storage, only 0.29% of cefalexin-resistant and 0.00001% of amoxicillin resistant *E. coli* remained (Figure 7a). However, the model predicts that tetracycline resistant bacteria will increase over this time by 25% due to ongoing selective pressure and low fitness cost. Importantly, multidrug resistant *E. coli* become undetectable.

**Figure 7:**
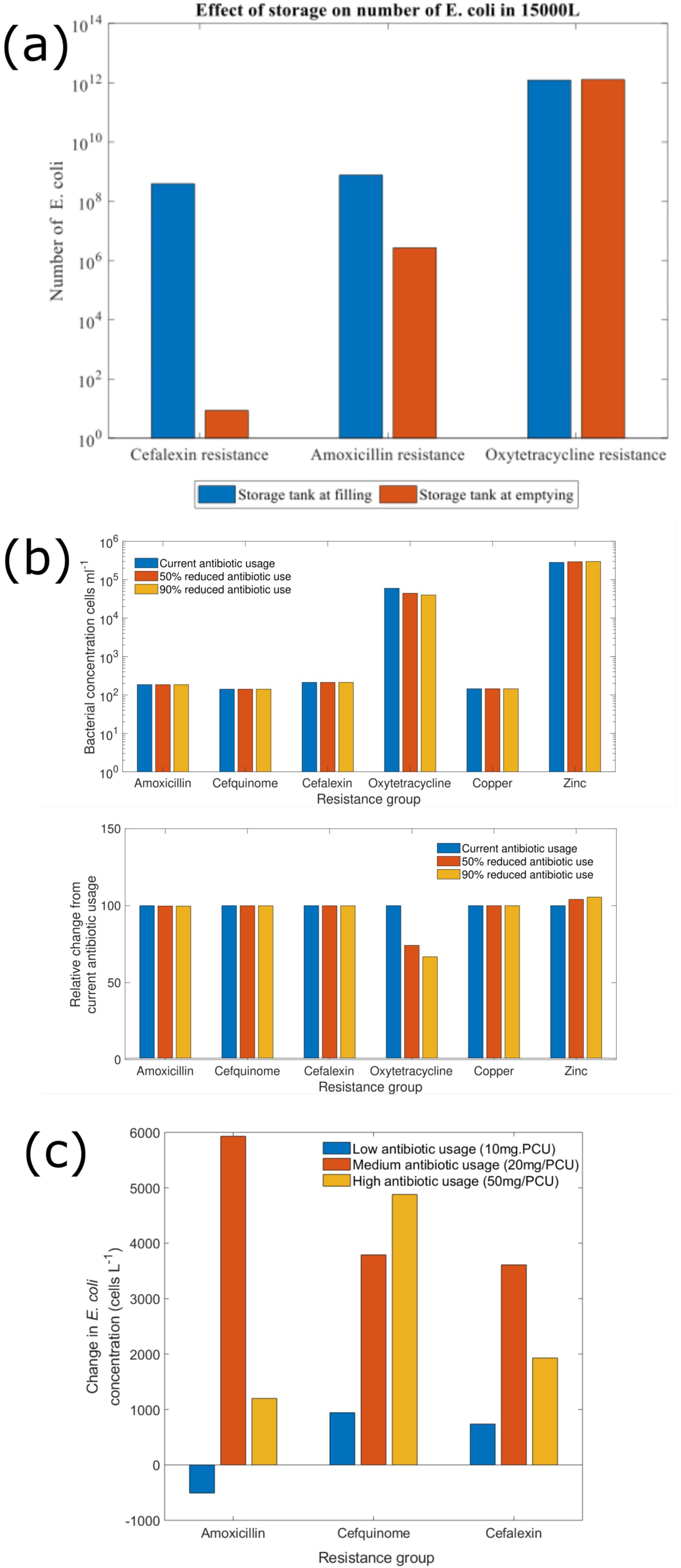
Model simulations of altered farm practise or antibiotic use. (a) Storing slurry without fresh inflow for 60 days is predicted to reduce resistance. Cephalexin resistance is reduced by more than 99.99% while amoxicillin resistance is reduced by more than 99%. (b) Model predictions of current antibiotic usage (9.7 mg/PCU) compared to a 50% reduction (4.85mg/PCU) and 90% reduction (0.97mg/PCU) show negligible impact on slurry tank resistance levels. (c) Model predictions of the change in resistant *E. coli* in the tank when using a 4^th^ generation cephalosporin instead of a 1^st^ generation cephalosporin on low, medium and high antibiotic usage farms showing increased resistance to all relevant antibiotics.

### Simulations of altered antibiotic use support criteria for responsible use

Simulations of on-farm antibiotic use (∼9.7 mg/Population Correction Unit (PCU) in 2017) result in low levels of penicillin and cephalosporin resistance, consistent with the empirical data. We simulated further reductions in antibiotics entering the tank to either 50% or 10% of current use. Neither reduction had a material impact on either resistance (Figure 7b) but there is a small reduction in tetracycline resistance (33% reduction in resistance at 10% usage) because of the reduced selective pressure for tetracycline resistance.

Very few cephalosporin resistant *E. coli* were detected in the farm samples (detailed above). Thus, we also simulated a return to use of the critically important 4^th^ generation cephalosporin (cefquinome) in place of cefalexin (1^st^ generation), assuming that cefquinome resistance also confers resistance to cefalexin. After accounting for the lower recommended dosage of cefquinome relative to cefalexin, we found cefquinome use increased resistance to both cefquinome and cefalexin of only 0.65% and 0.35% increase respectively (Figure 7c). To represent high antibiotic use following an outbreak of disease, we simulated 50 mg/PCU of cefquinome used in place of cefalexin. Such a scenario was predicted to select an increase of cefquinome resistance of only 3.55%.

## Discussion

### The slurry tank is a critical measurement and control point for AMR

The bacterial community and ARGs in the slurry tank appear to be maintained in a state of dynamic equilibrium, with a balance between input of fresh microorganisms from the cattle, and decline, as observed in the mini slurry-tank experiments. This equilibrium is also evident in the observed stability of the virome of the same tank over the same sampling period (73). The slurry tank maintains an array of ARGs, many of which have been found in other animal wastes. These include MLS genes such as *mefA* (24, 29, 74) and the *cfxA* group of beta-lactamase genes (24, 74, 75). The association of *cfxA* with Gram-positive organisms suggests that AMR phenotyping should routinely include a Gram-positive sentinel; *Enterococcus* spp. may be suitable because of their use in water quality analysis (76) and the inclusion of *E. faecium* in the ESKAPE pathogens list (77). Tetracycline resistance genes such as *tetW* and *tetM* have also been frequently found in cattle and swine waste (29, 78, 79). Although present in low quantities relative to other ARGs, *tetM* has the potential for selection and possible mobilisation (*e.g.*, IS*Ecp1* or Tn*916*-like elements). Consequently, the tank appears to be a critical sampling location, representative of the AMR status of the farm as a whole, reflecting current and previous antibiotic use. The presence of resistance genes to antibiotics with no recorded use (e.g. quinolone resistance, MLS genes) are likely to reflect broader environmental, and possibly human, input into the farm microbiome.

At a superficial level, the slurry tank appears to meet many criteria presumed to define a ‘hotspot’ for AMR, which cite a high abundance of bacterial populations and the routine presence of antimicrobial residue (80). However, the concept of an AMR ‘hotspot’, where bacterial and antimicrobial abundance are assumed to lead to increases in AMR prevalence, alongside the related concept of ‘reservoir’, assumed to represent the nascent AMR genes circulating in the environment poised to be mobilised through antimicrobial exposure, are open to critique (81). Our findings suggest that the tank, rather than generating resistance, can ameliorate resistance, depending on the waste management practice, and that slurry be stored for at least two months without fresh slurry inputs to the system/tank. Thus, the tank is neither a hotspot nor a reservoir, but, if managed appropriately, can be a critical control point for reducing the transmission of ARGs and ARB from livestock into the wider environment.

### Agricultural AMR policy should combine responsible antibiotic use with effective waste management

Policy and industry guidance to reduce AMR focus on reduced or responsible agricultural antimicrobial use (25, 82, 83), including the cessation of use of human critical antibiotics. Our findings provide evidence in support of responsible use. Simulations of reductions below the already low level of 9 mg/PCU did not predict reductions in penicillin and cephalosporin resistance below current levels. However, reduced tetracycline use led to reduced tetracycline resistance, associated with the environmental stability of this antibiotic, suggesting that prudent antibiotic use could also include antibiotic choice encouraging use for those with shorter half-lives where medically appropriate. While our findings suggested that use of 3^rd^ and 4^th^ generation cephalosporins did not lead to substantial increases associated resistances, once such resistances are established, relevant genes, e.g. CTX-M, can be selected for by 1^st^ generation use. Although UK policy initiatives have greatly reduced the use of 3^rd^/4^th^ generation cephalosporins on UK dairy farms, globally their use remains prevalent, e.g. Ceftiofur (3^rd^ generation cephalosporin) is routinely used in the US to treat metritis (84, 85) and mastitis (86). Eliminating the use of these antibiotics in agricultural production should still be an important goal of national and global policies to mitigate the environmental dissemination of AMR (87).

A policy focus on antibiotic use is limited because of the need to use antibiotics to treat sick livestock. We also showed that waste management practice provides an additional mechanism to control AMR, by reducing the prevalence of resistance genes and key microbial phyla in slurry prior to soil amendment. Specifically, secondary storage of slurry for a period of 60 days, without fresh inflow, would significantly reduce the levels of ARB within the tank, representing an opportunity for rational farm design and practice to minimize AMR outcomes. This result is also concordant with other practices for mitigating AMR on farms, including the use of anaerobic digestion (79, 88), vermicomposting and solid-liquid separation (43).

Two qPCR-based studies surveying Finnish swine and dairy farms reported that storage of animal manure slurry coincided with significant increases in select tetracycline, sulphonamide and aminoglycoside resistance genes when compared to fresh manure (28, 29). However, the farms involved in these studies used storage systems which received regular fresh inflow during the sampling period. Our metagenomic analyses of mini-tanks indicate that in the absence of fresh input a range of ARG classes decline (e.g. aminoglycoside and beta-lactam ARGs) or remain relatively stable (e.g. MLS ARGs). Moreover, culture-based results confirm an overall reduction in antibiotic resistant *E. coli* in slurry stored without inflow. Collectively, this provides empirical evidence supporting existing UK guidelines regarding the storage of slurry without further input as a means of reducing environmental exposure to AMR determinants.

### Evaluation of co-selection needs alternative approaches

Aminoglycoside, tetracycline and sulphonamide resistance genes were found on the same contigs. The result is consistent with sulphonamide resistance being co-selected by concurrent use of multiple antimicrobials because aminoglycosides and tetracycline were the two antibiotic classes used most during the sampling period. We anticipated finding evidence of co-occurrence of ARGs and MRGs in assembled metagenomic data, in accordance with other studies (19, 24, 89). However, apart from antibiotic resistance associated with Tn*21*-like elements carrying integrons, we found no evidence for such linkage in the slurry metagenomes or sequenced *E. coli* strains. This lack of evidence might not be evidence of absence of ARG-MRG co-occurrence, as these genes may not necessarily be genetically linked on a chromosome or on plasmids, and yet still be subject to co-selection if they reside in the same cell. Accordingly, the use of long-read or hybrid genome sequencing of strains selected for zinc or copper resistance may be more appropriate for detecting the co-occurrence of ARGs and MRGs (90).

## Conclusions

We have conducted a longitudinal, interdisciplinary study of the dynamics of AMR in a dairy slurry tank. The microbiota was in a state of dynamic equilibrium, with fresh input of bacteria from the animals balanced by natural decay. Antibiotic resistance was maintained, reflecting current and previous veterinary practice, as well as interaction with the broader environment. The slurry tank is therefore both a natural measurement point for on-farm resistance, as well as a control countermeasure point for resistance being released into the wider environment (land and water). The spread of antibiotic resistance into the wider environment through slurry application can be mitigated by a combination of responsible antibiotic use, including low total quantity, avoidance of human critical antibiotics, and antibiotic choice with shorter half-lives, with slurry storage. These approaches can mitigate spread of AMR into the environment from one of the world’s largest sources of AMR pollution.

## Author Contributions

Michelle Baker: Methodology, Formal Analysis, Investigation, Writing – Original Draft, Visualization

Alexander D Williams: Formal Analysis, Investigation, Data Curation, Writing – Original Draft, Visualization

Steven P.T. Hooton: Methodology, Investigation Richard Helliwell: Investigation, Writing – Original Draft Elizabeth King: Methodology, Investigation, Supervision Thomas Dodsworth: Methodology, Investigation

Rosa María Baena-Nogueras: Methodology, Investigation

Andrew Warry: Methodology, Formal Analysis, Data Curation, Visualization Catherine Ortori: Investigation

Henry Todman: Methodology, Investigation Charlotte J. Gray-Hammerton: Investigation Ryan Cook: Data Curation

Alexander C. W. Pritchard: Investigation Ethan Iles: Investigation

Richard Emes: Conceptualization, Acquisition of Funding Michael A Jones: Conceptualization, Acquisition of Funding

David A Barrett: Conceptualization, Supervision, Acquisition of Funding Theodore Kypraios: Conceptualization, Supervision, Acquisition of Funding

Stephen J Ramsden: Conceptualization, Acquisition of Funding, Writing – Reviewing and Editing

Chris Hudson: Conceptualization, Acquisition of Funding, Writing – Reviewing and Editing

Andrew D Millard: Conceptualization, Acquisition of Funding, Writing – Reviewing and Editing

Sujatha Raman: Conceptualization, Acquisition of Funding, Supervision, Writing – Reviewing and Editing

Helen West: Conceptualization, Acquisition of Funding, Supervision

Carol Morris: Conceptualization, Acquisition of Funding, Supervision, Writing – Reviewing and Editing

Rachel L Gomes: Conceptualization, Acquisition of Funding, Supervision, Writing – Reviewing and Editing

Christine E R Dodd: Conceptualization, Acquisition of Funding, Supervision, Writing – Reviewing and Editing

Jan-Ulrich Kreft: Conceptualization, Acquisition of Funding, Supervision, Writing – Reviewing and Editing

Jon L Hobman: Conceptualization, Acquisition of Funding, Supervision, Writing – Reviewing and Editing

Dov J Stekel: Conceptualization, Formal Analysis, Supervision, Writing – Reviewing and Editing, Project Administration, Acquisition of Funding

## Data Availability

Genome sequences are deposited with NCBI under BioProject PRJNA736866. Metagenome sequences are deposited with the ENA under Study Accession PRJEB38990. Mathematical models are deposited in BioModels as MODEL1909100001 and MODEL1909120002. All other data, including all details of accession numbers of genome and metagenome sequences, are on Figshare (https://figshare.com) under project number 133176.

## Acknowledgments

This work was supported by Antimicrobial Resistance Cross Council Initiative supported by the seven United Kingdom research councils (NE/N019881/1). ADW was funded by a NERC STARS PhD scholarship (NE/M009106/1). CJGH and ACWP were funded by the BBSRC Nottingham-Rothamsted Doctoral Training Partnership (BB/M008770/1). RC is supported by a scholarship from the Medical Research Foundation National PhD Training Programme in Antimicrobial Resistance Research (MRF-145-0004-TPG-AVISO). Bioinformatic analysis was made possible via the use of MRC-CLIMB (<u>MR/L015080/1</u>.) and CLIMB-BIG DATA (<u>MR/T030062/1</u>). We thank Chris Thomas, Emma Allaway and David Allaway for support with grant development. We thank Nigel Armstrong and the farm staff for their time, patience and support. We thank the external advisory board members for support, critique and feedback of our research: Nigel Brown, Brian Dalby, Gareth Hateley, Derek Armstrong, Katherine Grace, Marion Bos, Stacey Brown, Milen Georgiev, Javier Dominquez, Martin Rigley, Karen Heaton, Rupert Hough, Josh Onyango, Amreesh Mishra, Paul Wilson and Phil O’Neil. DJS thanks W Levine Stekel and CF Levine Stekel for useful conversation and advice. We thank Emma Hooley for her support throughout the entire research process.

## References

1. Van Boeckel TP, Glennon EE, Chen D, Gilbert M, Robinson TP, Grenfell BT, et al. Reducing antimicrobial use in food animals. Science. 2017;357(6358):1350–2.

2. Vogt D, Overesch G, Endimiani A, Collaud A, Thomann A, Perreten V. Occurrence and genetic characteristics of third-generation cephalosporin-resistant Escherichia coli in Swiss retail meat. Microbial drug resistance. 2014;20(5):485–94.

3. Lammie SL, Hughes JM. Antimicrobial resistance, food safety, and one health: the need for convergence. Annual review of food science and technology. 2016;7:287–312.

4. Gundogan N, Avci E. Occurrence and antibiotic resistance of E scherichia coli, S taphylococcus aureus and B acillus cereus in raw milk and dairy products in T urkey. International journal of dairy technology. 2014;67(4):562–9.

5. Silveira-Filho VM, Luz IS, Campos APF, Silva WM, Barros MPS, Medeiros ES, et al. Antibiotic resistance and molecular analysis of Staphylococcus aureus isolated from cow’s milk and dairy products in northeast Brazil. Journal of food protection. 2014;77(4):583–91.

6. Marti R, Tien Y-C, Murray R, Scott A, Sabourin L, Topp E. Safely coupling livestock and crop production systems: how rapidly do antibiotic resistance genes dissipate in soil following a commercial application of swine or dairy manure? Applied and environmental microbiology. 2014;80(10):3258–65.

7. Guron GK, Arango-Argoty G, Zhang L, Pruden A, Ponder MA. Effects of Dairy Manure-Based Amendments and Soil Texture on Lettuce-and Radish-Associated Microbiota and Resistomes. Msphere. 2019;4(3).

8. Joy SR, Bartelt-Hunt SL, Snow DD, Gilley JE, Woodbury BL, Parker DB, et al. Fate and transport of antimicrobials and antimicrobial resistance genes in soil and runoff following land application of swine manure slurry. Environmental science & technology. 2013;47(21):12081–8.

9. Neher TP, Ma L, Moorman TB, Howe A, Soupir ML. Seasonal variations in export of antibiotic resistance genes and bacteria in runoff from an agricultural watershed in Iowa. Science of The Total Environment. 2020;738:140224.

10. FAO. Livestock and environment statistics: manure and greenhouse gas emissions. Global, regional and country trends 1990–2018. 2020.

11. Willett W, Rockström J, Loken B, Springmann M, Lang T, Vermeulen S, et al. Food in the Anthropocene: the EAT–Lancet Commission on healthy diets from sustainable food systems. The Lancet. 2019;393(10170):447–92.

12. Smith K, Williams A. Production and management of cattle manure in the UK and implications for land application practice. Soil Use and Management. 2016;32:73–82.

13. DEFRA. The national inventory and map of livestock manure loadings to agricultural land (Manures-GIS) - WQ0103. 2016.

14. Oliver SP, Murinda SE, Jayarao BM. Impact of antibiotic use in adult dairy cows on antimicrobial resistance of veterinary and human pathogens: a comprehensive review. Foodborne pathogens and disease. 2011;8(3):337–55.

15. DDC. Dairy Youngstock Project-Wales, Full Report. 2015.

16. CHAWG. GB Cattle Health & Welfare Group, 4th Report. 2018.

17. CHAWG. GB Cattle Health & Welfare Group, 5th Report. 2020.

18. Griffiths BE, Dai White G, Oikonomou G. A cross-sectional study into the prevalence of dairy cattle lameness and associated herd-level risk factors in England and Wales. Frontiers in veterinary science. 2018;5:65.

19. Pal C, Bengtsson-Palme J, Kristiansson E, Larsson DJ. Co-occurrence of resistance genes to antibiotics, biocides and metals reveals novel insights into their co-selection potential. BMC genomics. 2015;16(1):964.

20. Davies R, Wales A. Antimicrobial resistance on farms: a review including biosecurity and the potential role of disinfectants in resistance selection. Comprehensive reviews in food science and food safety. 2019;18(3):753–74.

21. Seiffert SN, Hilty M, Perreten V, Endimiani A. Extended-spectrum cephalosporin-resistant Gram-negative organisms in livestock: an emerging problem for human health? Drug Resistance Updates. 2013;16(1-2):22–45.

22. Ibrahim DR, Dodd CE, Stekel DJ, Ramsden SJ, Hobman JL. Multidrug resistant, extended spectrum β-lactamase (ESBL)-producing Escherichia coli isolated from a dairy farm. FEMS microbiology ecology. 2016;92(4).

23. Wichmann F, Udikovic-Kolic N, Andrew S, Handelsman J. Diverse antibiotic resistance genes in dairy cow manure. MBio. 2014;5(2):e01017–13.

24. Zhou B, Wang C, Zhao Q, Wang Y, Huo M, Wang J, et al. Prevalence and dissemination of antibiotic resistance genes and coselection of heavy metals in Chinese dairy farms. Journal of hazardous materials. 2016;320:10–7.

25. FAO/OiE/WHO. Monitoring Global Progress on Addressing Antimicrobial Resistance, Analysis report of the second round of results of AMR country self-assessment survey. 2018.

26. UK-VARSS. Veterinary Antibiotic Resistance and Sales Surveillance Report (UK-VARSS 2019). 2020.

27. GOV.UK. Handling of manure and slurry to reduce antibiotic resistance 2016 [Available from: https://www.gov.uk/guidance/handling-of-manure-and-slurry-to-reduce-antibiotic-resistance.

28. Ruuskanen M, Muurinen J, Meierjohan A, Pärnänen K, Tamminen M, Lyra C, et al. Fertilizing with animal manure disseminates antibiotic resistance genes to the farm environment. Journal of environmental quality. 2016;45(2):488–93.

29. Muurinen J, Stedtfeld R, Karkman A, Parnanen K, Tiedje J, Virta M. Influence of manure application on the environmental resistome under Finnish agricultural practice with restricted antibiotic use. Environmental science & technology. 2017;51(11):5989–99.

30. Hurst JJ, Oliver JP, Schueler J, Gooch C, Lansing S, Crossette E, et al. Trends in antimicrobial resistance genes in manure blend pits and long-term storage across dairy farms with comparisons to antimicrobial usage and residual concentrations. Environmental science & technology. 2019;53(5):2405–15.

31. Durso LM, Harhay GP, Bono JL, Smith TP. Virulence-associated and antibiotic resistance genes of microbial populations in cattle feces analyzed using a metagenomic approach. Journal of microbiological methods. 2011;84(2):278–82.

32. Noyes NR, Yang X, Linke LM, Magnuson RJ, Cook SR, Zaheer R, et al. Characterization of the resistome in manure, soil and wastewater from dairy and beef production systems. Scientific reports. 2016;6(1):1–12.

33. Rovira P, McAllister T, Lakin SM, Cook SR, Doster E, Noyes NR, et al. Characterization of the microbial resistome in conventional and “raised without antibiotics” beef and dairy production systems. Frontiers in microbiology. 2019;10:1980.

34. Gaeta NC, Bean E, Miles AM, de Carvalho DUOG, Alemán MAR, Carvalho JS, et al. A cross-sectional study of dairy cattle metagenomes reveals increased antimicrobial resistance in animals farmed in a heavy metal contaminated environment. Frontiers in microbiology. 2020;11.

35. Himathongkham S, Bahari S, Riemann H, Cliver D. Survival of Escherichia coli O157: H7 and Salmonella typhimurium in cow manure and cow manure slurry. FEMS Microbiology Letters. 1999;178(2):251–7.

36. Placha I, Venglovský J, Sasakova N, Svoboda I. The effect of summer and winter seasons on the survival of Salmonella typhimurium and indicator micro-organisms during the storage of solid fraction of pig slurry. Journal of Applied Microbiology. 2001;91(6):1036–43.

37. Nicholson FA, Groves SJ, Chambers BJ. Pathogen survival during livestock manure storage and following land application. Bioresource technology. 2005;96(2):135–43.

38. Sharma M, Reynnells R. Importance of Soil Amendments: Survival of Bacterial Pathogens in Manure and Compost Used as Organic Fertilizers. Microbiology Spectrum. 2016;4(4).

39. Schubert H, Morley K, Puddy EF, Arbon R, Findlay J, Mounsey O, et al. Reduced Antibacterial Drug Resistance and blaCTX-M β-Lactamase Gene Carriage in Cattle-Associated Escherichia coli at Low Temperatures, at Sites Dominated by Older Animals, and on Pastureland: Implications for Surveillance. Applied and Environmental Microbiology. 2021;87(6).

40. Halbert LW, Kaneene JB, Ruegg PL, Warnick LD, Wells SJ, Mansfield LS, et al. Evaluation of antimicrobial susceptibility patterns in Campylobacter spp isolated from dairy cattle and farms managed organically and conventionally in the midwestern and northeastern United States. Journal of the American Veterinary Medical Association. 2006;228(7):1074–81.

41. Feng X, Littier HM, Knowlton KF, Garner E, Pruden A. The impacts of feeding milk with antibiotics on the fecal microbiome and antibiotic resistance genes in dairy calves. Canadian Journal of Animal Science. 2019;100(1):69–76.

42. Feng X, Chambers LR, Knowlton KF. Antibiotic resistance genes in the faeces of dairy cows following short-term therapeutic and prophylactic antibiotic administration. Journal of Applied Animal Research. 2020;48(1):34–7.

43. Oliver JP, Hurst JJ, Gooch CA, Stappenbeck A, Sassoubre L, Aga DS. On-farm screw-press/rotary drum treatment of dairy manure associated antibiotic residues and resistance. Wiley Online Library; 2020. Report No.: 0047-2425.

44. CLSI. Performance Standards for Antimicrobial Susceptibility Testing-M100S. 2016.

45. Yang H, Wei S-H, Hobman JL, Dodd CE. Antibiotic and metal resistance in Escherichia coli isolated from pig slaughterhouses in the United Kingdom. Antibiotics. 2020;9(11):746.

46. Hooton SP, Pritchard AC, Asiani K, Gray-Hammerton CJ, Stekel DJ, Crossman LC, et al. Laboratory stock variants of the archetype silver resistance plasmid pMG101 demonstrate plasmid fusion, loss of transmissibility, and transposition of Tn7/pco/sil Into the host chromosome. Frontiers in Microbiology. 2021;12.

47. Seemann T. Prokka: rapid prokaryotic genome annotation. Bioinformatics. 2014;30(14):2068–9.

48. Chen K-T, Lu CL. CSAR-web: a web server of contig scaffolding using algebraic rearrangements. Nucleic acids research. 2018;46(W1):W55–W9.

49. Bortolaia V, Kaas RS, Ruppe E, Roberts MC, Schwarz S, Cattoir V, et al. ResFinder 4.0 for predictions of phenotypes from genotypes. Journal of Antimicrobial Chemotherapy. 2020;75(12):3491–500.

50. Carattoli A, Zankari E, Garcìa-Fernandez A, Larsen MV, Lund O, Villa L, et al. PlasmidFinder and pMLST: in silico detection and typing of plasmids. Antimicrobial agents and chemotherapy. 2014.

51. Baena-Nogueras R, Ortori C, Barrett D, Gomes R, editors. Analysis of veterinary antibiotics in dairy environments by liquid chromatography–mass spectrometry. 15th International Conference on Environmental Science and Technology Rhodes, Greece; 2017.

52. Martin M. Cutadapt removes adapter sequences from high-throughput sequencing reads. EMBnet journal. 2011;17(1):10–2.

53. Joshi N, Fass J. Sickle: A sliding-window, adaptive, quality-based trimming tool for FastQ files (Version 1.33)[Software]. 2011.

54. Chen S, Zhou Y, Chen Y, Gu J. fastp: an ultra-fast all-in-one FASTQ preprocessor. Bioinformatics. 2018;34(17):i884–i90.

55. Li D, Liu C-M, Luo R, Sadakane K, Lam T-W. MEGAHIT: an ultra-fast single-node solution for large and complex metagenomics assembly via succinct de Bruijn graph. Bioinformatics. 2015;31(10):1674–6.

56. Arango-Argoty G, Garner E, Pruden A, Heath LS, Vikesland P, Zhang L. DeepARG: a deep learning approach for predicting antibiotic resistance genes from metagenomic data. Microbiome. 2018;6(1):1–15.

57. Seemann T. Abricate. Github2020.

58. Doster E, Lakin SM, Dean CJ, Wolfe C, Young JG, Boucher C, et al. MEGARes 2.0: a database for classification of antimicrobial drug, biocide and metal resistance determinants in metagenomic sequence data. Nucleic acids research. 2020;48(D1):D561–D9.

59. Leplae R, Lima-Mendez G, Toussaint A. ACLAME: a CLAssification of Mobile genetic Elements, update 2010. Nucleic acids research. 2010;38(suppl_1):D57–D61.

60. Wang LYR, Jokinen CC, Laing CR, Johnson RP, Ziebell K, Gannon VP. Multi-year persistence of verotoxigenic Escherichia coli (VTEC) in a closed Canadian beef herd: a cohort study. Frontiers in microbiology. 2018;9:2040.

61. Pal C, Bengtsson-Palme J, Rensing C, Kristiansson E, Larsson DJ. BacMet: antibacterial biocide and metal resistance genes database. Nucleic acids research. 2014;42(D1):D737–D43.

62. Hyatt D, Chen G-L, LoCascio PF, Land ML, Larimer FW, Hauser LJ. Prodigal: prokaryotic gene recognition and translation initiation site identification. BMC bioinformatics. 2010;11(1):1–11.

63. Menzel P, Ng KL, Krogh A. Fast and sensitive taxonomic classification for metagenomics with Kaiju. Nature communications. 2016;7(1):1–9.

64. NCBI. National Center for Biotechnology Information: Bethesda (MD): National Library of Medicine (US), National Center for Biotechnology Information; [1988] [Available from: https://www.ncbi.nlm.nih.gov/.

65. Altschul SF, Gish W, Miller W, Myers EW, Lipman DJ. Basic local alignment search tool. Journal of molecular biology. 1990;215(3):403–10.

66. Morgulis A, Coulouris G, Raytselis Y, Madden TL, Agarwala R, Schäffer AA. Database indexing for production MegaBLAST searches. Bioinformatics. 2008;24(16):1757–64.

67. Shaw LM, Blanchard A, Chen Q, An X, Davies P, Tötemeyer S, et al. DirtyGenes: testing for significant changes in gene or bacterial population compositions from a small number of samples. Scientific reports. 2019;9(1):1–10.

68. Baker M, Hobman JL, Dodd CE, Ramsden SJ, Stekel DJ. Mathematical modelling of antimicrobial resistance in agricultural waste highlights importance of gene transfer rate. FEMS Microbiology Ecology. 2016;92(4).

69. Arya S, Williams A, Reina SV, Knapp CW, Kreft J-U, Hobman JL, et al. Towards a general model for predicting minimal metal concentrations co-selecting for antibiotic resistance plasmids. Environmental Pollution. 2021;275:116602.

70. Malik-Sheriff RS, Glont M, Nguyen TV, Tiwari K, Roberts MG, Xavier A, et al. BioModels—15 years of sharing computational models in life science. Nucleic acids research. 2020;48(D1):D407–D15.

71. Magiorakos A-P, Srinivasan A, Carey R, Carmeli Y, Falagas M, Giske C, et al. Multidrug-resistant, extensively drug-resistant and pandrug-resistant bacteria: an international expert proposal for interim standard definitions for acquired resistance. Clinical microbiology and infection. 2012;18(3):268–81.

72. Byrne-Bailey K, Gaze W, Kay P, Boxall A, Hawkey P, Wellington E. Prevalence of sulfonamide resistance genes in bacterial isolates from manured agricultural soils and pig slurry in the United Kingdom. Antimicrobial Agents and Chemotherapy. 2009;53(2):696–702.

73. Cook R, Hooton S, Trivedi U, King L, Dodd CE, Hobman JL, et al. Hybrid assembly of an agricultural slurry virome reveals a diverse and stable community with the potential to alter the metabolism and virulence of veterinary pathogens. Microbiome. 2021;9(1):1–17.

74. Li B, Yang Y, Ma L, Ju F, Guo F, Tiedje JM, et al. Metagenomic and network analysis reveal wide distribution and co-occurrence of environmental antibiotic resistance genes. The ISME journal. 2015;9(11):2490–502.

75. Zhang Y-J, Hu H-W, Gou M, Wang J-T, Chen D, He J-Z. Temporal succession of soil antibiotic resistance genes following application of swine, cattle and poultry manures spiked with or without antibiotics. Environmental Pollution. 2017;231:1621–32.

76. Bartram J, Pedley S. Microbiological Analyses. In: Bartram J, Ballance R, editors. Water quality monitoring: a practical guide to the design and implementation of freshwater quality studies and monitoring programmes: CRC Press; 1996.

77. Rice LB. Federal funding for the study of antimicrobial resistance in nosocomial pathogens: no ESKAPE. The University of Chicago Press; 2008.

78. Kyselková M, Jirout J, Vrchotová N, Schmitt H, Elhottová D. Spread of tetracycline resistance genes at a conventional dairy farm. Frontiers in microbiology. 2015;6:536.

79. Zhang J, Wang Z, Wang Y, Zhong H, Sui Q, Zhang C, et al. Effects of graphene oxide on the performance, microbial community dynamics and antibiotic resistance genes reduction during anaerobic digestion of swine manure. Bioresource technology. 2017;245:850–9.

80. Lima T, Domingues S, Da Silva GJ. Manure as a Potential Hotspot for Antibiotic Resistance Dissemination by Horizontal Gene Transfer Events. Veterinary Sciences. 2020;7(3):110.

81. Helliwell R, Raman S, Morris C. Environmental imaginaries and the environmental sciences of antimicrobial resistance. Environment and Planning E: Nature and Space. 2020:2514848620950752.

82. O’Neill J. Antimicrobial Resistance: Tackling a crisis for the health and wealth of nations. 2014.

83. WHO. Global Action Plan on Antimicrobial Resistance. 2015.

84. Espadamala A, Pereira R, Pallares P, Lago A, Silva-Del-Rio N. Metritis diagnosis and treatment practices in 45 dairy farms in California. Journal of dairy science. 2018;101(10):9608–16.

85. Silva T, de Oliveira E, Pérez-Báez J, Risco C, Chebel R, Cunha F, et al. Economic comparison between ceftiofur-treated and nontreated dairy cows with metritis. Journal of Dairy Science. 2021.

86. USDA A, VS, CEAH, NAHMS. Dairy 2014 Milk Quality, Milking Procedures, and Mastitis on U.S. Dairies, 2014. 2014.

87. Hughes A, Roe E, Hocknell S. Food supply chains and the antimicrobial resistance challenge: On the framing, accomplishments and limitations of corporate responsibility. Environment and Planning A: Economy and Space. 2021:0308518X211015255.

88. Sui Q, Zhang J, Chen M, Tong J, Wang R, Wei Y. Distribution of antibiotic resistance genes (ARGs) in anaerobic digestion and land application of swine wastewater. Environmental Pollution. 2016;213:751–9.

89. Li X, Rensing C, Vestergaard G, Nesme J, Gupta S, Arumugam M, et al. Metagenomic evidence for co-occurrence of antibiotic, biocide and metal resistance genes in pigs. bioRxiv. 2021.

90. Brown CL, Keenum IM, Dai D, Zhang L, Vikesland PJ, Pruden A. Critical evaluation of short, long, and hybrid assembly for contextual analysis of antibiotic resistance genes in complex environmental metagenomes. Scientific reports. 2021;11(1):1–12.

