## Supplementary Data for "Antimicrobial resistance in dairy slurry tanks: a critical point for measurement and control"

Baker M, Williams AD, Hooton SPT et al.

**This document contains supplementary information not supplied in the main text. Specifically:**

Supplementary Text 1: Full description of model

Supplementary Text 2: Model equations

Supplementary Text 3: Supplementary Experimental Methods for Parameter Estimation

Supplementary Table S1: Sampling dates for microbiology and water quality analyses.

Supplementary Table S2: Details of antibiotic disks used for AST analysis

Supplementary Table S3: Recorded antibiotic use for dairy cattle on the farm

Supplementary Table S4: Bacterial phyla identified in the slurry

Supplementary Table S5: Model parameters

Supplementary Table S6: Bayes factors for microbial growth model fits

References (for Supplementary Text 1 and Table S5)

Supplementary Figure S1: ARG and MRG collocation on slurry tank contigs

Supplementary Figure S2: ARG and MRG collocation on mini tank contigs

Supplementary Figure S3: Contigs containing co-located ARGs

Supplementary Figure S4: Full Model schematic

Supplementary Figure S5: Model fit to antibiotic data

Supplementary Figure S6: Model fit to *E. coli* count data

Supplementary Figure S7: Mini-tank Model Schematic

Supplementary Figure S8: Model workflow

Supplementary Figure S9: Storage Methods Schematic

Supplementary Figure S10: Impact on AMR of different storage methods

### Supplementary Text 1: Model Details

#### Model development

The ODE model (Supplementary Text 2) consists of a multidimensional ODE with variables for each of the *E. coli* subpopulations in the slurry that are sensitive or resistant to each of the six antimicrobials (64 equations), plus variables for the antibiotic concentrations in slurry (penicillin, tetracycline, cefalexin, cefquinome) and a variable for slurry volume (1 equation). A model schematic is shown in Supplementary Figure S4 and the full model parameter values can be found in Supplementary Table S4.

We assume slurry flows into the main slurry tank at a constant rate ($\lambda$), and contains a constant concentration of *E. coli*. We estimate the flow rate of slurry to be 1480 litres per hour, calculated based on the number of cattle housed on the farm and expected levels of faecal and effluent waste. The main slurry tank has a maximum capacity of 3 million litres, hence would fill every 85 days and requires regular emptying. We simulate the tank being emptied every 60 days (removing 90% of the slurry) which allows it to fill to almost maximum capacity and is consistent with farm records that show slurry being spread on fields every 1-2 months in spreading season. Due to the farm infrastructure, all slurry passes through the main tank so we have assumed that even during periods when no slurry is spread on fields due to government regulation, the tank will still be filled and emptied on a regular basis into the lagoon for storage.

We assume that the *E. coli* concentration in the inflow pipe (v) is $2.16 \times 10^8$ CFU $L^{-1}$ ($2.16 \times 10^5$ CFU $mL^{-1}$). This is based on microbial counts from three independent samples taken on different days from the underground reservoir on the farm, which feeds directly into the slurry inflow pipe.

We assume metal concentrations of copper and zinc are constant within the slurry inflow and the tank, based on ICP-MS results from slurry tank samples taken in 2015, the concentrations used are 22.317 mg $L^{-1}$ (Cu) and 32.158 mg $L^{-1}$ (Zn) (Arya *et al.* 2021). We neglect any settling of the heavy metals within the tank and assume that the stirrer ensures a homogeneous distribution of all content. We have calculated the death rates $\delta_{Cu}$ and $\delta_{Zn}$ at these constant metal concentrations using metal toxicity data (Ivask et al. 2009).

Cephqinome and Cefalexin were assumed to exponentially decay over time within the slurry tank and that the decay rate constants could be estimated independently from the full model as they are calculated in uncoupled equations. The decay rate constants were estimated directly from measurements of the levels of these antibiotics during mini-tank experiments (Supplementary Text 3). To use these estimates as data for parameter estimation we assumed that the pre-spiked samples represented levels just before sampling, and the post spiked samples represented levels 24 hours after the sampling time. Assuming a simple decay model we estimated the decay rate of both antibiotics in this realistic slurry environment using a Bayesian MCMC Metropolis-Hastings algorithm, running for 100000 iterations and discarding 25% as burn-in. This gave a satisfactory trace of each parameter and good sampling of the posterior distribution (data not shown). Using the mean of the posterior as the model parameters the experimental data gave a good fit with the predicted values within the experimental error bars for all time points (Supplementary Figure S6). The decay rate constants for amoxicillin and tetracycline were estimated based on values from literature (Supplementary Table S5).

The parameters (δ, $\alpha_{CEF}$, $\alpha_{CEX}$, $\alpha_{Cu}$, $\alpha_{Z}$, and $\rho$) were estimated simultaneously using MCMC estimation, using *E. coli* count data from the control tanks, and tanks with metal added, either alone or with cefquinome (Supplementary Figure S4). For the purposes of parameter estimation. A model variant for the mini tanks (Schematic in Supplementary Figure S7) allows the investigation of possible interventions. In this model, resistance to only three antimicrobials is considered because we use the model to estimate the parameters associated with the single added antibiotic, alongside copper and zinc concentrations.

All parameters were run for 50000 iterations (10% burn-in) to give posterior distribution estimates for each parameter. The estimation algorithm was verified by relaxing the prior distributions in the model and convergence was still observed. Joint posterior distributions of the estimated parameters show no major correlation between the parameters (data not shown). The posterior distribution for each parameter was then used to randomly select a parameter value for multiple runs of the mathematical model.

Given the long time periods over which the slurry is being stored it is likely that slurry tanks experience temperature variation. This will consist of diurnal temperature fluctuations and longer term daily or seasonal temperature change. To relate the temperature in slurry to a specific growth rate parameter a small bacterial growth experiment was performed (Supplementary Text 3). The rate of growth was estimated over the first 5 hours of the experiment, before a net loss of *E. coli* was observed. This gave a specific growth rate of $0.08h^{-1}$ without nutrients and $0.183h^{-1}$ with nutrients (1% UHT milk). To take account of the effect of temperature on growth we considered the relationship between the specific growth rate, *r* and the temperature to be linear and using our estimates for *r* at 37°C and a cut-off of 5°C where we expected no growth, we estimated the gradient and intercept of the equation relating temperature to growth. We considered both a fixed growth rate over the course of the experiment and a variable daily rate. This gave four alternative sets of growth rates, r, over the course of the experiment. It was not possible to get an estimate of the environmental death rate, δ, from the experimental work so we considered three different possibilities for the relationship between the death rate and temperature: (i) a constant value of δ, (ii) that δ is variable with a relative difference to *r* and thirdly that δ is variable with an absolute difference to *r*. MCMC was used to estimate parameters for each model variant and select the best model. The MCMC would not converge to the model with a daily variable rate of *r* and constant δ so this left 6 different model variants to consider. To choose which model variant best fitted the experimental data Bayesian competition was performed using Bayes factor analysis (Supplementary Table S6), with thermodynamic integration, this showed that a constant growth and death rates were in low nutrients gave the best model fit.

Bacteria growth is modelled logistically with a constant maximum specific growth rate (r) of 0.08 $h^{-1}$. The actual rate will be reduced in individual subspecies by fitness costs and bacteriostatic agents. Following Baker *et al.* 2016, the carrying capacity ($N_{max}$) is of 1 x $10^{10}$ cells $L^{-1}$. We considered models of *E. coli* growth linked to the recorded atmospheric temperature, but model selection procedures predicted that a constant temperature model gave the best fit to experimental data. All bacteria carrying resistance genes have an associated fitness costs estimated from literature or data (Supplementary Table 7). The fitness costs are considered additive except for resistance to cephalosporins. Since resistance to, for example, a 3rd generation cephalosporin would confer resistance to 2nd and 1st generation cephalosporins there would be no selective advantage to holding both resistances, hence we consider the fitness cost of these to be the maximum of the individual costs.

Each subpopulation is associated with several death rates. All populations are subject to a baseline constant environmental death rate, accounting for death due to the hostile conditions including unsuitable oxygen levels; pH or temperature; predation by protozoa, bacteria, or phages . The MCMC (detailed above) predicted an approximately Normal distribution for the baseline death rate, therefore in repeats of model simulations we randomly select a value from this distribution. Additional death will be due to antimicrobial agents in the tank, with populations sensitive to a particular agent affected by that death rate, dependent on the concentration present in the tank. The death rate for these is assumed to depend on the MIC following a Hill function according to either,

$$\delta_A = 1 - 2\frac{\left(\frac{A}{V}\right)^2}{MIC_A{}^2 + \left(\frac{A}{V}\right)^2}$$

for bactericidal antibiotics or,

$$\frac{\left(\frac{A}{V}\right)^2}{MIC_A{}^2 + \left(\frac{A}{V}\right)^2}$$

for bacteriostatic antibiotics.

We allow horizontal gene transfer of resistance between all subpopulations of bacteria according to the level of contact between the bacteria, with a single gene transfer rate parameter (β) that we set at 1 x $10^{-6}$ $h^{-1}$, following Baker *et al.* 2016.

The mathematical model is an ODE model with a single simulation running over the timescale of 365 days. To account for the input of antibiotics (for which we have daily experimental data) and slurry tank emptying (taking place every 60 days) we serialise the model on a daily basis as shown in the workflow (Supplementary Figure S8). To properly account for variation in the parameters that we found in the MCMC parameter estimation the mathematical ODE model was run for 1000 independent simulations and the median and median absolute deviation were calculated.

This model was deposited in BioModels (Malik-Sheriff *et al.* 2020) and assigned the identifier MODEL1909100001. The secondary storage model is also deposited under the identifier MODEL1909120002 and follows a similar workflow.

**Secondary Storage Models**

A secondary storage intervention could be implemented in several ways: we assessed two different methods (Supplementary Figure S9). One method uses two connected tanks, with the first always taking inflow from the animals, and the second store taking inflow from the first tank. The alternative has two disconnected tanks, with one tank taking inflow whilst the other is isolated, then alternating. In both cases, there are differential equations for bacterial populations and antiobiotic concentrations for each tank. The model predicts that use of both of these methods results in very similar reductions in resistance, with only marginal differences due to the irregularity of antibiotic inflow (Supplementary Figure S10).

**Supplementary Text 2:** Full Model Equations

$$
\begin{aligned}
\frac{dS}{dt} = r\left(1-\frac{N}{N_{max}}\right) S\delta_o + \lambda\rho_S v - S(\delta+\delta_a+\delta_c+\delta_{cq}+\delta_{cu}+\delta_{zn}) + \beta S(-R_{o,c,cq,a,cu,zn}/(S+R_{o,c,cq,a,cu,zn})\\
+ R_{o,c,cq,cu,zn}/(S+R_{o,c,cq,cu,zn}) + R_{o,cq,a,cu,zn}/(S+R_{o,cq,a,cu,zn}) + R_{c,cq,a,cu,zn}/(S+R_{c,cq,a,cu,zn})\\
+ R_{o,c,cq,a,zn}/(S+R_{o,c,cq,a,zn}) + R_{o,c,cq,a,cu}/(S+R_{o,c,cq,a,cu}) + R_{o,c,a,cu,zn}/(S+R_{o,c,a,cu,zn})\\
+ R_{o,cq,cu,zn}/(S+R_{o,cq,cu,zn}) + R_{c,cq,cu,zn}/(S+R_{c,cq,cu,zn}) + R_{c,cq,a,zn}/(S+R_{c,cq,a,zn})\\
+ R_{c,cq,a,cu}/(S+R_{c,cq,a,cu}) + R_{o,c,cq,a}/(S+R_{o,c,cq,a}) + R_{cq,a,cu,zn}/(S+R_{cq,a,cu,zn})\\
+ R_{o,c,cq,cu}/(S+R_{o,c,cq,cu}) + R_{o,c,cq,zn}/(S+R_{o,c,cq,zn}) + R_{o,cq,a,zn}/(S+R_{o,cq,a,zn})\\
+ R_{o,cq,a,cu}/(S+R_{o,cq,a,cu}) + R_{c,a,cu,zn}/(S+R_{c,a,cu,zn}) + R_{o,a,cu,zn}/(S+R_{o,a,cu,zn})\\
+ R_{o,c,cu,zn}/(S+R_{o,c,cu,zn}) + R_{o,c,a,zn}/(S+R_{o,c,a,zn}) + R_{o,c,a,cu}/(S+R_{o,c,a,cu})\\
+ R_{cq,cu,zn}/(S+R_{cq,cu,zn}) + R_{o,cq,zn}/(S+R_{o,cq,zn}) + R_{o,cq,cu}/(S+R_{o,cq,cu}) + R_{c,cq,zn}/(S+R_{c,cq,zn})\\
+ R_{c,cq,cu}/(S+R_{c,cq,cu}) + R_{o,c,cq}/(S+R_{o,c,cq}) + R_{cq,a,zn}/(S+R_{cq,a,zn}) + R_{cq,a,cu}/(S+R_{cq,a,cu})\\
+ R_{o,cq,a}/(S+R_{o,cq,a}) + R_{c,cq,a}/(S+R_{c,cq,a}) + R_{a,cu,zn}/(S+R_{a,cu,zn}) + R_{c,cu,zn}/(S+R_{c,cu,zn})\\
+ R_{c,a,zn}/(S+R_{c,a,zn}) + R_{c,a,cu}/(S+R_{c,a,cu}) + R_{o,cu,zn}/(S+R_{o,cu,zn}) + R_{o,a,zn}/(S+R_{o,a,zn})\\
+ R_{o,a,cu}/(S+R_{o,a,cu}) + R_{o,c,zn}/(S+R_{o,c,zn}) + R_{o,c,cu}/(S+R_{o,c,cu}) + R_{o,c,a}/(S+R_{o,c,a})\\
+ R_{cq,zn}/(S+R_{cq,zn}) + R_{cq,cu}/(S+R_{cq,cu}) + R_{c,cq}/(S+R_{c,cq}) + R_{cq,a}/(S+R_{cq,a})\\
+ R_{o,cq}/(S+R_{o,cq}) + R_{cu,zn}/(S+R_{cu,zn}) + R_{a,zn}/(S+R_{a,zn}) + R_{a,cu}/(S+R_{a,cu})\\
+ R_{c,zn}/(S+R_{c,zn}) + R_{c,cu}/(S+R_{c,cu}) + R_{c,a}/(S+R_{c,a}) + R_{o,zn}/(S+R_{o,zn})\\
+ R_{o,cu}/(S+R_{o,cu}) + R_{o,a}/(S+R_{o,a}) + R_{o,c}/(S+R_{o,c}) + R_{cq}/(S+R_{cq})\\
+ R_{zn}/(S+R_{zn}) + R_{cu}/(S+R_{cu}) + R_a/(S+R_a) + R_c/(S+R_c) + R_o/(S+R_o))
\end{aligned}
$$

$$
\begin{aligned}
\frac{dR_o}{dt} = r\left(1-\frac{N}{N_{max}}\right)(1-\alpha_o xy)R_o + \lambda\rho_R v - R_o(\delta+\delta_a+\delta_{cq}+\delta_c+\delta_{cu}+\delta_{zn}) + \beta R_o(-R_{c,cq,a,cu,zn}/(R_{c,cq,a,cu,zn}+R_o)\\
- R_{c,a,cu,zn}/(R_{c,a,cu,zn}+R_o) - R_{c,cq,cu,zn}/(R_{c,cq,cu,zn}+R_o) - R_{cq,a,cu,zn}/(R_{cq,a,cu,zn}+R_o)\\
- R_{c,cq,a,zn}/(R_{c,cq,a,zn}+R_o) - R_{c,cq,a,cu}/(R_{c,cq,a,cu}+R_o) - R_{cq,cu,zn}/(R_{cq,cu,zn}+R_o)\\
- R_{c,cq,a}/(R_{c,cq,a}+R_o) - R_{c,cq,cu}/(R_{c,cq,cu}+R_o) - R_{c,cq,zn}/(R_{c,cq,zn}+R_o)\\
- R_{cq,a,zn}/(R_{cq,a,zn}+R_o) - R_{cq,a,cu}/(R_{cq,a,cu}+R_o) - R_{c,cu,zn}/(R_{c,cu,zn}+R_o)\\
- R_{a,cu,zn}/(R_{a,cu,zn}+R_o) + S/(S+R_o) - R_{c,a,cu}R_o/(R_{c,a,cu}+R_o) - R_{c,a,zn}/(R_{c,a,zn}+R_o)\\
- R_{cq,zn}/(R_{cq,zn}+R_o) - R_{cq,cu}/(R_{cq,cu}+R_o) - R_{c,cq}/(R_{c,cq}+R_o)\\
- R_{cq,a}R_{cq}/(R_{cq,a}+R_o) - R_{cu,zn}/(R_{cu,zn}+R_o) - R_{a,cu}/(R_{a,cu}+R_o)\\
- R_{a,zn}/(R_{a,zn}+R_o) - R_{c,zn}/(R_{c,zn}+R_o) - R_c/(R_o+R_c) - R_{c,cu}/(R_{c,cu}+R_o)\\
- R_{c,a}/(R_{c,a}+R_o) - R_a/(R_o+R_a) - R_{cu}/(R_o+R_{cu}) - R_{zn}/(R_o+R_{zn}) - R_{cq}/(R_o+R_{cq}))
\end{aligned}
$$

$$
\begin{aligned}
\frac{dR_c}{dt} = r\left(1-\frac{N}{N_{max}}\right)(1-\alpha_c ex)R_c\delta_o + \lambda\rho_R v - R_c(\delta+\delta_a+\delta_{cq}+\delta_{cu}+\delta_{zn}) + \beta R_c(-R_{o,cq,a,cu,zn}/(R_{o,cq,a,cu,zn}+R_c)\\
- R_{o,a,cu,zn}/(R_{o,a,cu,zn}+R_c) - R_{o,cq,cu,zn}/(R_{o,cq,cu,zn}+R_c) - R_{cq,a,cu,zn}/(R_{cq,a,cu,zn}+R_c)\\
- R_{o,cq,a,zn}/(R_{o,cq,a,zn}+R_c) - R_{o,cq,a,cu}/(R_{o,cq,a,cu}+R_c) - R_{cq,cu,zn}/(R_{cq,cu,zn}+R_c) - R_{cq,a,zn}/(R_{cq,a,zn}+R_c)\\
- R_{cq,a,cu}/(R_{cq,a,cu}+R_c) - R_{o,cq,zn}/(R_{o,cq,zn}+R_c) - R_{o,cq,a}/(R_{o,cq,a}+R_c) - R_{o,cq,cu}/(R_{o,cq,cu}+R_c)\\
- R_{a,cu,zn}/(R_{a,cu,zn}+R_c) - R_{o,cu,zn}/(R_{o,cu,zn}+R_c) + S/(S+R_c) - R_{o,a,cu}/(R_{o,a,cu}+R_c) - R_{o,a,zn}/(R_{o,a,zn}+R_c)\\
- R_{cq,zn}/(R_{cq,zn}+R_c) - R_{cq,cu}/(R_{cq,cu}+R_c) - R_{o,cq}/(R_{o,cq}+R_c) - R_{cq,a}/(R_{cq,a}+R_c)\\
- R_{cu,zn}/(R_{cu,zn}+R_c) - R_{a,zn}/(R_{a,zn}+R_c) - R_{a,cu}/(R_{a,cu}+R_c) - R_{o,zn}/(R_{o,zn}+R_c)\\
- R_o/(R_o+R_c) - R_{o,cu}/(R_{o,cu}+R_c) - R_{o,a}/(R_{o,a}+R_c) - R_{cq}/(R_c+R_{cq}) - R_a/(R_c+R_a)\\
- R_{cu}/(R_c+R_{cu}) - R_{zn}/(R_c+R_{zn}))
\end{aligned}
$$

$$\frac{dR_a}{dt} = r\left(1-\frac{N}{N_{max}}\right)(1-\alpha_a mox)R_a\delta_o + \lambda\rho_R v - R_a(\delta+\delta_c+\delta_{cu}++\delta_{cq}+\delta_{zn})$$
$$+\beta R_a(-R_{o,c,cq,cu,zn}/(R_{o,c,cq,cu,zn}+R_a) - R_{o,c,cu,zn}/(R_{o,c,cu,zn}+R_a) - R_{o,cq,cu,zn}/(R_{o,cq,cu,zn}+R_a)$$
$$- R_{c,cq,cu,zn}/(R_{c,cq,cu,zn}+R_a) - R_{o,c,cq,zn}/(R_{o,c,cq,zn}+R_a) - R_{o,c,cq,cu}/(R_{o,c,cq,cu}+R_a)$$
$$- R_{c,cq,zn}/(R_{c,cq,zn}+R_a) - R_{c,cq,cu}/(R_{c,cq,cu}+R_a) - R_{o,c,cq}/(R_{o,c,cq}+R_a) + S/(S+R_a) - R_{cq,cu,zn}/(R_{cq,cu,zn}+R_a)$$
$$- R_{o,cq,zn}/(R_{o,cq,zn}+R_a) - R_{o,cq,zn}/(R_{o,cq,zn}+R_a) - R_{o,cq,cu}/(R_{o,cq,cu}+R_a) - R_{c,cu,zn}/(R_{c,cu,zn}+R_a)$$
$$- R_{o,cu,zn}/(R_{o,cu,zn}+R_a) - R_{o,c,cu}/(R_{o,c,cu}+R_a) - R_{o,c,zn}/(R_{o,c,zn}+R_a) - R_{cq,zn}/(R_{cq,zn}+R_a)$$
$$- R_{cq,cu}/(R_{cq,cu}+R_a) - R_{o,cq}/(R_{o,cq}+R_a) - R_{cu,zn}/(R_{cu,zn}+R_a) - R_{c,cq}/(R_{c,cq}+R_a) - R_{c,zn}/(R_{c,zn}+R_a)$$
$$- R_{o,zn}/(R_{o,zn}+R_a) - R_{c,cu}/(R_{c,cu}+R_a) - R_{o,cu}/(R_{o,cu}+R_a) - R_o/(R_o+R_a) - R_{o,c}/(R_{o,c}+R_a)$$
$$- R_c/(R_c+R_a) - R_{cq}/(R_{cq}+R_a) - R_{cu}/(R_a+R_{cu}) - R_{zn}/(R_a+R_{zn}))$$
$$\frac{dR_{cu}}{dt} = r\left(1-\frac{N}{N_{max}}\right)(1-\alpha_c u)R_{cu}\delta_o + \lambda\rho_R v - R_{cu}(\delta+\delta_a+\delta_{cq}+\delta_c+\delta_{zn})$$
$$+\beta R_{cu}(-R_{o,c,cq,a,zn}/(R_{o,c,cq,a,zn}+R_{cu}) - R_{o,c,a,zn}/(R_{o,c,a,zn}+R_{cu}) - R_{o,c,cq,zn}/(R_{o,c,cq,zn}+R_{cu})$$
$$- R_{o,cq,a,zn}/(R_{o,cq,a,zn}+R_{cu}) - R_{c,cq,a,zn}/(R_{c,cq,a,zn}+R_{cu}) - R_{o,c,cq,a}/(R_{o,c,cq,a}+R_{cu}) + S/(S+R_{cu})$$
$$- R_{o,cq,zn}/(R_{o,cq,zn}+R_{cu}) - R_{c,cq,zn}/(R_{c,cq,zn}+R_{cu}) - R_{c,cq,a}/(R_{c,cq,a}+R_{cu}) - R_{cq,a,zn}/(R_{cq,a,zn}$$
$$+R_{cu}) - \beta R_{o,c,cq}R_{cu}/(R_{o,c,cq}+R_{cu}) - \beta R_{o,cq,a}R_{cu}/(R_{o,cq,a}+R_{cu}) - R_{c,a,zn}/(R_{c,a,zn}+R_{cu})$$
$$- R_{o,a,zn}/(R_{o,a,zn}+R_{cu}) - R_{o,c,zn}/(R_{o,c,zn}+R_{cu}) - R_{o,c,a}/(R_{o,c,a}+R_{cu}) - R_{cq,zn}/(R_{cq,zn}+R_{cu})$$
$$- R_{o,cq}/(R_{o,cq}+R_{cu}) - R_{c,cq}/(R_{c,cq}+R_{cu}) - R_{cq,a}/(R_{cq,a}+R_{cu}) - R_{a,zn}/(R_{a,zn}+R_{cu}) - R_{c,zn}/(R_{c,zn}+R_{cu})$$
$$- R_{c,a}/(R_{c,a}+R_{cu}) - R_{o,zn}/(R_{o,zn}+R_{cu}) - R_o/(R_o+R_{cu}) - R_{o,a}/(R_{o,a}+R_{cu}) - R_{o,c}/(R_{o,c}+R_{cu})$$
$$- R_{cq}/(R_{cq}+R_{cu}) - R_c/(R_c+R_{cu}) - R_a/(R_a+R_{cu}) - R_{zn}/(R_{cu}+R_{zn}))$$
$$\frac{dR_{zn}}{dt} = r\left(1-\frac{N}{N_{max}}\right)(1-\alpha_z n)R_{zn}\delta_o + \lambda\rho_R v - R_{zn}(\delta+\delta_a+\delta_{cq}+\delta_c+\delta_{cu})$$
$$+\beta R_{zn}(-R_{o,c,cq,a,cu}/(R_{o,c,cq,a,cu}+R_{zn}) - R_{o,c,a,cu}/(R_{o,c,a,cu}+R_{zn}) - R_{o,c,cq,cu}/(R_{o,c,cq,cu}+R_{zn})$$
$$- R_{o,cq,a,cu}/(R_{o,cq,a,cu}+R_{zn}) - R_{c,cq,a,cu}/(R_{c,cq,a,cu}+R_{zn}) - R_{o,c,cq,a}/(R_{o,c,cq,a}+R_{zn}) + S/(S+R_{zn})$$
$$- R_{o,cq,cu}/(R_{o,cq,cu}+R_{zn}) - R_{c,cq,cu}/(R_{c,cq,cu}+R_{zn}) - R_{c,cq,a}/(R_{c,cq,a}+R_{zn}) - R_{cq,a,cu}/(R_{cq,a,cu}+R_{zn})$$
$$- R_{o,c,cq}/(R_{o,c,cq}+R_{zn}) - R_{o,cq,a}/(R_{o,cq,a}+R_{zn}) - R_{c,a,cu}/(R_{c,a,cu}+R_{zn}) - R_{o,a,cu}/(R_{o,a,cu}+R_{zn})$$
$$- R_{o,c,cu}/(R_{o,c,cu}+R_{zn}) - R_{o,c,a}/(R_{o,c,a}+R_{zn}) - R_{cq,cu}/(R_{cq,cu}+R_{zn}) - R_{o,cq}/(R_{o,cq}+R_{zn})$$
$$- R_{cq,a}/(R_{cq,a}+R_{zn}) - R_{c,cq}/(R_{c,cq}+R_{zn}) - R_{a,cu}/(R_{a,cu}+R_{zn}) - R_{c,cu}/(R_{c,cu}+R_{zn})$$
$$- R_{c,a}/(R_{c,a}+R_{zn}) - R_{o,cu}/(R_{o,cu}+R_{zn}) - R_{o,c}/(R_{o,c}+R_{zn}) - R_{o,a}/(R_{o,a}+R_{zn}) - R_o/(R_o+R_{zn})$$
$$R_{cq}/(R_{cq}+R_{zn}) - R_c/(R_c+R_{zn}) - R_a/(R_a+R_{zn}) - R_{cu}/(R_{cu}+R_{zn}))$$
$$\frac{dR_{cq}}{dt} = r\left(1-\frac{N}{N_{max}}\right)(1-\alpha_c ef)R_{cq}\delta_o + \lambda\rho_R v - R_{cq}(\delta+\delta_a+\delta_{cu}+\delta_{zn})$$
$$+\beta R_{cq}(-R_{o,c,a,cu,zn}/(R_{o,c,a,cu,zn}+R_{cq}) - R_{o,c,cu,zn}/(R_{o,c,cu,zn}+R_{cq}) - R_{o,a,cu,zn}/(R_{o,a,cu,zn}+R_{cq})$$
$$- R_{c,a,cu,zn}/(R_{c,a,cu,zn}+R_{cq}) + S/(S+R_{cq}) - R_{o,c,a,zn}/(R_{o,c,a,zn}+R_{cq}) - R_{o,c,a,cu}/(R_{o,c,a,cu}+R_{cq})$$
$$- R_{o,cu,zn}/(R_{o,cu,zn}+R_{cq}) - R_{c,cu,zn}/(R_{c,cu,zn}+R_{cq}) - R_{c,a,zn}/(R_{c,a,zn}+R_{cq}) - R_{c,a,cu}/(R_{c,a,cu}+R_{cq})$$
$$- R_{o,c,a}/(R_{o,c,a}+R_{cq}) - R_{a,cu,zn}/(R_{a,cu,zn}+R_{cq}) - R_{o,c,cu}/(R_{o,c,cu}+R_{cq}) - R_{o,c,zn}/(R_{o,c,zn}+R_{cq})$$
$$- R_{o,a,zn}/(R_{o,a,zn}+R_{cq}) - R_{o,a,cu}/(R_{o,a,cu}+R_{cq}) - R_{cu,zn}/(R_{cu,zn}+R_{cq}) - R_{o,zn}/(R_{o,zn}+R_{cq})$$
$$- R_{o,cu}/(R_{o,cu}+R_{cq}) - R_{c,cu}/(R_{c,cu}+R_{cq}) - R_{c,zn}/(R_{c,zn}+R_{cq}) - R_{o,c}/(R_{o,c}+R_{cq})$$
$$- R_{a,zn}/(R_{a,zn}+R_{cq}) - R_{a,cu}/(R_{a,cu}+R_{cq}) - R_{o,a}/(R_{o,a}+R_{cq}) - R_{c,a}/(R_{c,a}+R_{cq}) - R_o/(R_o+R_{cq})$$
$$- R_{zn}/(R_{cq}+R_{zn}) - R_{cu}/(R_{cq}+R_{cu}) - R_c/(R_c+R_{cq}) - R_a/(R_{cq}+R_a))$$
$$\frac{dR_{o,c}}{dt} = r\left(1-\frac{N}{N_{max}}\right)(1-\alpha_{o,c})R_{o,c} + \lambda\rho_R v - R_{o,c}(\delta+\delta_a+\delta_{cq}+\delta_{cu}+\delta_{zn})$$
$$+\beta R_{o,c}(-R_{cq,a,cu,zn}/(R_{cq,a,cu,zn}+R_{o,c}) - R_{a,cu,zn}/(R_{a,cu,zn}+R_{o,c}) - R_{cq,cu,zn}/(R_{cq,cu,zn}+R_{o,c})$$
$$- R_{cq,a,zn}/(R_{cq,a,zn}+R_{o,c}) - R_{cq,a}/(R_{cq,a}+R_{o,c}) - R_{cq,a,cu}/(R_{cq,a,cu}+R_{o,c}) - R_{cq,cu}/(R_{o,c}+R_{cq,cu})$$
$$- R_{cq,zn}/(R_{cq,zn}+R_{o,c}) + S/(S+R_{o,c}) - R_{cu,zn}/(R_{cu,zn}+R_{o,c}) - R_{a,cu}/(R_{o,c}+R_{a,cu}) - R_{cq}/(R_{o,c}+R_{cq})$$
$$- R_{zn}/(R_{o,c}+R_{zn}) - R_{cu}/(R_{o,c}+R_{cu}) - R_a/(R_{o,c}+R_a)) + \beta 2R_oR_c/(R_o+R_c)$$

$$
\begin{aligned}
\frac{dR_{o,a}}{dt} &= r\left(1-\frac{N}{N_{max}}\right)(1-\alpha_{o,a})R_{o,a}+\lambda\rho_R v-R_{o,a}(\delta+\delta_c+\delta_{cq}+\delta_{cu}+\delta_{zn})\\
&+\beta R_{o,a}(-R_{c,cq,cu,zn}/(R_{c,cq,cu,zn}+R_{o,a})-R_{c,cu,zn}/(R_{c,cu,zn}+R_{o,a})+S/(S+R_{o,a})-R_{cq,cu,zn}/(R_{cq,cu,zn}+R_{o,a})\\
&-R_{c,cq,zn}/(R_{c,cq,zn}+R_{o,a})-R_{c,cq,cu}/(R_{c,cq,cu}+R_{o,a})-R_{c,cq}/(R_{c,cq}+R_{o,a})-R_{cq,zn}/(R_{cq,zn}+R_{o,a})\\
&-R_{cq,cu}/(R_{o,a}+R_{cq,cu})-R_{cu,zn}/(R_{o,a}+R_{cu,zn})-R_{c,cu}/(R_{o,a}+R_{c,cu})-R_{c,zn}/(R_{o,a}+R_{c,zn})\\
&-R_{cq}/(R_{o,a}+R_{cq})-R_{zn}/(R_{o,a}+R_{zn})-R_{cu}/(R_{o,a}+R_{cu})-R_c/(R_{o,a}+R_c))+\beta 2R_oR_a/(R_o+R_a)\\
\frac{dR_{o,cu}}{dt} &= r\left(1-\frac{N}{N_{max}}\right)(1-\alpha_{o,cu})R_{o,cu}+\lambda\rho_R v-R_{o,cu}(\delta+\delta_a+\delta_{cq}+\delta_c+\delta_{zn})\\
&+\beta R_{o,cu}(-R_{c,cq,a,zn}/(R_{c,cq,a,zn}+R_{o,cu})-R_{c,a,zn}/(R_{c,a,zn}+R_{o,cu})-R_{c,cq,zn}/(R_{c,cq,zn}+R_{o,cu})\\
&-R_{cq,a,zn}/(R_{cq,a,zn}+R_{o,cu})+S/(S+R_{o,cu})-R_{c,cq,a}/(R_{c,cq,a}+R_{o,cu})-R_{cq,zn}/(R_{cq,zn}+R_{o,cu})\\
&-R_{c,cq}/(R_{o,cu}+R_{c,cq})-R_{cq,a}/(R_{o,cu}+R_{cq,a})-R_{a,zn}/(R_{o,cu}+R_{a,zn})-R_{c,zn}/(R_{c,zn}+R_{o,cu})\\
&-R_{c,a}/(R_{c,a}+R_{o,cu})-R_{cq}/(R_{o,cu}+R_{cq})-R_{zn}/(R_{o,cu}+R_{zn})-R_a/(R_{o,cu}+R_a)-R_c/(R_{o,cu}+R_c))\\
&+\beta 2R_oR_{cu}/(R_o+R_{cu})\\
\frac{dR_{o,zn}}{dt} &= r\left(1-\frac{N}{N_{max}}\right)(1-\alpha_{o,zn})R_{o,zn}+\lambda\rho_R v-R_{o,zn}(\delta+\delta_a+\delta_{cq}+\delta_c+\delta_{cu})\\
&+\beta R_{o,zn}(-R_{c,cq,a,cu}/(R_{c,cq,a,cu}+R_{o,zn})-R_{c,a,cu}/(R_{c,a,cu}+R_{o,zn})-R_{c,cq,cu}/(R_{c,cq,cu}+R_{o,zn})\\
&-R_{cq,a,cu}/(R_{cq,a,cu}+R_{o,zn})-R_{c,cq,a}/(R_{c,cq,a}+R_{o,zn})-R_{c,cq}/(R_{c,cq}+R_{o,zn})-R_{cq,cu}/(R_{cq,cu}+R_{o,zn})\\
&-R_{cq,a}/(R_{cq,a}+R_{o,zn})+S/(S+R_{o,zn})-R_{a,cu}/(R_{o,zn}+R_{a,cu})-R_{c,cu}/(R_{c,cu}+R_{o,zn})\\
&-R_{c,a}/(R_{o,zn}+R_{c,a})-R_{cq}/(R_{o,zn}+R_{cq})-R_{cu}/(R_{o,zn}+R_{cu})-R_a/(R_{o,zn}+R_a)\\
&-R_c/(R_{o,zn}+R_c)))+\beta 2R_oR_{zn}/(R_o+R_{zn})\\
\frac{dR_{c,a}}{dt} &= r\left(1-\frac{N}{N_{max}}\right)(1-\alpha_{c,a})R_{c,a}\delta_o+\lambda\rho_R v-R_{c,a}(\delta+\delta_{cq}+\delta_{cu}+\delta_{zn})\\
&+\beta R_{c,a}(-R_{o,cq,zn}/(R_{o,cq,zn}+R_{c,a})-R_{o,cq,cu,zn}/(R_{o,cq,cu,zn}+R_{c,a})+S/(S+R_{c,a})-R_{o,cu,zn}/(R_{o,cu,zn}+R_{c,a})\\
&-R_{o,cq,cu}/(R_{o,cq,cu}+R_{c,a})-R_{cq,zn}/(R_{cq,zn}+R_{c,a})-R_{cq,cu}/(R_{cq,cu}+R_{c,a})-R_{o,cq}/(R_{c,a}+R_{o,cq})\\
&-R_{cu,zn}/(R_{c,a}+R_{cu,zn})-R_{o,zn}/(R_{o,zn}+R_{c,a})-R_{cq}/(R_{c,a}+R_{cq})-R_{zn}/(R_{c,a}+R_{zn})-R_{cu}/(R_{c,a}+R_{cu})\\
&-R_o/(R_{c,a}+R_o))+\beta 2R_cR_a/(R_c+R_a)\\
\frac{dR_{c,cu}}{dt} &= r\left(1-\frac{N}{N_{max}}\right)(1-\alpha_{c,cu})R_{c,cu}\delta_o+\lambda\rho_R v-R_{c,cu}(\delta+\delta_{cq}+\delta_a+\delta_{zn})\\
&+\beta R_{c,cu}(-R_{o,cq,a,zn}/(R_{o,cq,a,zn}+R_{c,cu})-R_{o,a,zn}/(R_{o,a,zn}+R_{c,cu})-R_{o,cq,zn}/(R_{o,cq,zn}+R_{c,cu})\\
&-R_{cq,a,zn}/(R_{cq,a,zn}+R_{c,cu})+S/(S+R_{c,cu})-R_{o,cq,a}/(R_{o,cq,a}+R_{c,cu})-R_{cq,zn}/(R_{cq,zn}+R_{c,cu})\\
&-R_{cq,a}/(R_{c,cu}+R_{cq,a})-R_{o,cq}/(R_{o,cq}+R_{c,cu})-R_{a,zn}/(R_{c,cu}+R_{a,zn})-R_{o,zn}/(R_{c,cu}+R_{o,zn})\\
&-R_{o,a}/(R_{o,a}+R_{c,cu})-R_{cq}/(R_{c,cu}+R_{cq})-R_{zn}/(R_{c,cu}+R_{zn})-R_a/(R_{c,cu}+R_a)-R_o/(R_{c,cu}+R_o))\\
&+\beta 2R_cR_{cu}/(R_c+R_{cu})\\
\frac{dR_{c,zn}}{dt} &= r\left(1-\frac{N}{N_{max}}\right)(1-\alpha_{c,zn})R_{c,zn}\delta_o+\lambda\rho_R v-R_{c,zn}(\delta+\delta_{cq}+\delta_a+\delta_{cu})\\
&+\beta R_{c,zn}(-R_{o,cq,a,cu}/(R_{o,cq,a,cu}+R_{c,zn})-R_{o,a,cu}/(R_{o,a,cu}+R_{c,zn})-R_{o,cq,cu}/(R_{o,cq,cu}+R_{c,zn})\\
&-R_{cq,a,cu}/(R_{cq,a,cu}+R_{c,zn})-R_{o,cq,a}/(R_{o,cq,a}+R_{c,zn})-R_{cq,a}/(R_{c,zn}+R_{cq,a})-R_{cq,cu}/(R_{c,zn}+R_{cq,cu})\\
&-R_{o,cq}/(R_{c,zn}+R_{o,cq})+S/(S+R_{c,zn})-R_{a,cu}/(R_{c,zn}+R_{a,cu})-R_{o,cu}/(R_{c,zn}+R_{o,cu})\\
&-R_{o,a}/(R_{o,a}+R_{c,zn})-R_{cq}/(R_{c,zn}+R_{cq})-R_{cu}/(R_{c,zn}+R_{cu})-R_a/(R_{c,zn}+R_a)-R_o/(R_{c,zn}+R_o))\\
&+\beta 2R_cR_{zn}/(R_c+R_{zn})\\
\frac{dR_{a,cu}}{dt} &= r\left(1-\frac{N}{N_{max}}\right)(1-\alpha_{a,cu})R_{a,cu}\delta_o+\lambda\rho_R v-R_{a,cu}(\delta+\delta_{cq}+\delta_c+\delta_{zn})\\
&+\beta R_{a,cu}(R_{o,c,cq,zn}/(-R_{o,c,cq,zn}+R_{a,cu})-R_{o,c,zn}/(R_{o,c,zn}+R_{a,cu})-R_{o,cq,zn}/(R_{o,cq,zn}+R_{a,cu})\\
&-R_{c,cq,zn}/(R_{c,cq,zn}+R_{a,cu})+S/(S+R_{a,cu})-R_{o,c,cq}/(R_{o,c,cq}+R_{a,cu})-R_{c,cq}/(R_{a,cu}+R_{c,cq})\\
&-R_{cq,zn}/(R_{cq,zn}+R_{a,cu})-R_{o,cq}/(R_{o,cq}+R_{a,cu})-R_{c,zn}/(R_{c,zn}+R_{a,cu})-R_{o,zn}/(R_{o,zn}+R_{a,cu})\\
&-R_{o,c}/(R_{o,c}+R_{a,cu})-R_{cq}/(R_{a,cu}+R_{cq})-R_{zn}/(R_{a,cu}+R_{zn})-R_c/(R_{a,cu}+R_c)-R_o/(R_{a,cu}+R_o))\\
&+\beta 2R_aR_{cu}/(R_a+R_{cu})
\end{aligned}
$$

$$
\begin{aligned}
\frac{dR_{a,zn}}{dt} &= r\left(1-\frac{N}{N_{max}}\right)(1-\alpha_{a,zn})R_{a,zn}\delta_o + \lambda\rho_R v - R_{a,zn}(\delta+\delta_c+\delta_{cq}+\delta_{cu}) \\
&+ \beta R_{a,zn}(-R_{o,c,cq,cu}/(R_{o,c,cq,cu}+R_{a,zn}) - R_{o,c,cu}/(R_{o,c,cu}+R_{a,zn}) - R_{o,cq,cu}/(R_{o,cq,cu}+R_{a,zn}) \\
&- R_{c,cq,cu}/(R_{c,cq,cu}+R_{a,zn}) - R_{o,c,cq}/(R_{o,c,cq}+R_{a,zn}) + S/(S+R_{a,zn}) - R_{c,cq}/(R_{a,zn}+R_{c,cq}) \\
&- R_{cq,cu}/(R_{cq,cu}+R_{a,zn}) - R_{o,cq}/(R_{a,zn}+R_{o,cq}) - R_{c,cu}/(R_{c,cu}+R_{a,zn}) - R_{cq}/(R_{a,zn}+R_{cq}) \\
&- R_{cu}/(R_{a,zn}+R_{cu}) - R_c/(R_{a,zn}+R_c) - R_o/(R_{a,zn}+R_o)) + \beta 2R_aR_{zn}/(R_a+R_{zn}) \\
\frac{dR_{cu,zn}}{dt} &= r\left(1-\frac{N}{N_{max}}\right)(1-\alpha_{cu,zn})R_{cu,zn}\delta_o + \lambda\rho_R v - R_{cu,zn}(\delta+\delta_{cq}+\delta_a+\delta_c) \\
&+ \beta R_{cu,zn}(-R_{o,c,cq,a}/(R_{o,c,cq,a}+R_{cu,zn}) - R_{o,c,a}/(R_{o,c,a}+R_{cu,zn}) - R_{o,c,cq}/(R_{o,c,cq}+R_{cu,zn}) \\
&- R_{o,cq,a}/(R_{o,cq,a}+R_{cu,zn}) - R_{c,cq,a}/(R_{c,cq,a}+R_{cu,zn}) + S/(S+R_{cu,zn}) - R_{o,cq}/(R_{cu,zn}+R_{o,cq}) \\
&- R_{c,cq}/(R_{cu,zn}+R_{c,cq}) - R_{cq,a}/(R_{cq,a}+R_{cu,zn}) - R_{c,a}/(R_{c,a}+R_{cu,zn}) - R_{o,a}/(R_{o,a}+R_{cu,zn}) \\
&- R_{o,c}/(R_{cu,zn}+R_{o,c}) - R_{cq}/(R_{cu,zn}+R_{cq}) - R_a/(R_{cu,zn}+R_a) - R_c/(R_{cu,zn}+R_c) - R_o/(R_{cu,zn}+R_o)) \\
&+ \beta 2R_{cu}R_{zn}/(R_{cu}+R_{zn}) \\
\frac{dR_{o,cq}}{dt} &= r\left(1-\frac{N}{N_{max}}\right)(1-\alpha_{o,cq})R_{o,cq} + \lambda\rho_R v - R_{o,cq}(\delta+\delta_a+\delta_{cu}+\delta_{zn}) \\
&+ \beta R_{o,cq}(-R_{c,a,cu,zn}/(R_{c,a,cu,zn}+R_{o,cq}) - R_{c,cu,zn}/(R_{c,cu,zn}+R_{o,cq}) - R_{a,cu,zn}/(R_{a,cu,zn}+R_{o,cq}) \\
&- R_{c,a,zn}/(R_{c,a,zn}+R_{o,cq}) - R_{c,a,cu}/(R_{c,a,cu}+R_{o,cq}) - R_{cu,zn}/(R_{cu,zn}+R_{o,cq}) - R_{c,a}/(R_{c,a}+R_{o,cq}) \\
&- R_{c,cu}/(R_{o,cq}+R_{c,cu}) - R_{c,zn}/(R_{c,zn}+R_{o,cq}) - R_{a,zn}/(R_{a,zn}+R_{o,cq}) - R_{a,cu}/(R_{o,cq}+R_{a,cu}) + S/(S+R_{o,cq}) \\
&- R_{zn}/(R_{o,cq}+R_{zn}) - R_{cu}/(R_{o,cq}+R_{cu}) - R_c/(R_{o,cq}+R_c) - R_a/(R_{o,cq}+R_a)) + \beta 2R_oR_{cq}/(R_o+R_{cq}) \\
\frac{dR_{cq,a}}{dt} &= r\left(1-\frac{N}{N_{max}}\right)(1-\alpha_{cq,a})R_{cq,a}\delta_o + \lambda\rho_R v - R_{cq,a}(\delta+\delta_{cu}+\delta_{zn}) \\
&+ \beta R_{cq,a}(-R_{o,c,cu,zn}/(R_{o,c,cu,zn}+R_{cq,a}) + S/(S+R_{cq,a}) - R_{o,cu,zn}/(R_{o,cu,zn}+R_{cq,a}) \\
&- R_{c,cu,zn}/(R_{c,cu,zn}+R_{cq,a}) - R_{o,c,zn}/(R_{o,c,zn}+R_{cq,a}) - R_{o,c,cu}/(R_{o,c,cu}+R_{cq,a}) - R_{c,zn}/(R_{c,zn}+R_{cq,a}) \\
&- R_{c,cu}/(R_{c,cu}+R_{cq,a}) - R_{o,c}/(R_{cq,a}+R_{o,c}) - R_{cu,zn}/(R_{cq,a}+R_{cu,zn}) - R_{o,zn}/(R_{cq,a}+R_{o,zn}) \\
&- R_{o,cu}/(R_{o,cu}+R_{cq,a}) - R_{zn}/(R_{cq,a}+R_{zn}) - R_{cu}/(R_{cq,a}+R_{cu}) - R_{cq}/(R_{cq,a}+R_o) -\} R_c/(R_{cq,a}+R_c)) \\
&+ \beta 2R_{cq}R_a/(R_{cq}+R_a) \\
\frac{dR_{c,cq}}{dt} &= r\left(1-\frac{N}{N_{max}}\right)(1-\alpha_{c,cq})R_{c,cq}\delta_o + \lambda\rho_R v - R_{c,cq}(\delta+\delta_{cu}+\delta_{zn}) \\
&+ \beta R_{c,cq}(-R_{o,a,cu,zn}/(R_{o,a,cu,zn}+R_{c,cq}) - R_{o,cu,zn}/(R_{o,cu,zn}+R_{c,cq}) - R_{a,cu,zn}/(R_{a,cu,zn}+R_{c,cq}) \\
&- R_{o,a,zn}/(R_{o,a,zn}+R_{c,cq}) + S/(S+R_{c,cq}) - R_{o,a,cu}/(R_{o,a,cu}+R_{c,cq}) - R_{cu,zn}/(R_{cu,zn}+R_{c,cq}) \\
&- R_{a,zn}/(R_{a,zn}+R_{c,cq}) - R_{a,cu}/(R_{a,cu}+R_{c,cq}) - R_{o,a}/(R_{c,cq}+R_{o,a}) - R_{o,cu}/(R_{o,cu}+R_{c,cq}) \\
&- R_{o,zn}/(R_{c,cq}+R_{o,zn}) - R_{zn}/(R_{c,cq}+R_{zn}) - R_{cu}/(R_{c,cq}+R_{cu}) - R_o/(R_{c,cq}+R_o) - R_a/(R_{c,cq}+R_a)) \\
&+ \beta 2R_cR_{cq}/(R_c+R_{cq}) \\
\frac{dR_{cq,cu}}{dt} &= r\left(1-\frac{N}{N_{max}}\right)(1-\alpha_{cq,cu})R_{cq,cu}\delta_o + \lambda\rho_R v - R_{cq,cu}(\delta+\delta_a+\delta_{zn}) \\
&+ \beta R_{cq,cu}(-R_{o,c,a,zn}/(R_{o,c,a,zn}+R_{cq,cu}) - R_{o,c,zn}/(R_{o,c,zn}+R_{cq,cu}) - R_{o,a,zn}/(R_{o,a,zn}+R_{cq,cu}) \\
&- R_{c,a,zn}/(R_{c,a,zn}+R_{cq,cu}) + S/(S+R_{cq,cu}) - R_{o,c,a}/(R_{o,c,a}+R_{cq,cu}) - R_{o,zn}/(R_{cq,cu}+R_{o,zn}) \\
&- R_{c,zn}/(R_{c,zn}+R_{cq,cu}) - R_{c,a}/(R_{cq,cu}+R_{c,a}) - R_{a,zn}/(R_{cq,cu}+R_{a,zn}) - R_{o,c}/(R_{o,c}+R_{cq,cu}) \\
&- R_{o,a}/(R_{o,a}+R_{cq,cu}) - R_{zn}/(R_{cq,cu}+R_{zn}) - R_o/(R_{cq,cu}+R_o) - R_c/(R_{cq,cu}+R_c) - R_a/(R_{cq,cu}+R_a)) \\
&+ \beta 2R_{cq}R_{cu}/(R_{cq}+R_{cu}) \\
\frac{dR_{cq,zn}}{dt} &= r\left(1-\frac{N}{N_{max}}\right)(1-\alpha_{cq,zn})R_{cq,zn}\delta_o + \lambda\rho_R v - R_{cq,zn}(\delta+\delta_a+\delta_{cu}) \\
&+ \beta R_{cq,zn}(-R_{o,c,a,cu}/(R_{o,c,a,cu}+R_{cq,zn}) - R_{o,c,cu}/(R_{o,c,cu}+R_{cq,zn}) - R_{o,a,cu}/(R_{o,a,cu}+R_{cq,zn}) \\
&- R_{c,a,cu}/(R_{c,a,cu}+R_{cq,zn}) + S/(S+R_{cq,zn}) - R_{o,c,a}/(R_{o,c,a}+R_{cq,zn}) - R_{o,cu}/(R_{cq,zn}+R_{o,cu}) \\
&- R_{cq,cu}/(R_{c,zn}+R_{cq,cu}) - R_{c,cu}/(R_{cq,zn}+R_{c,cu}) - R_{c,a}/(R_{cq,zn}+R_{c,a}) - R_{a,cu}/(R_{cq,zn}+R_{a,cu}) \\
&- R_{o,c}/(R_{cq,zn}+R_{o,c}) - R_{o,a}/(R_{cq,zn}+R_{o,a}) - R_{cu}/(R_{cq,zn}+R_{cu}) - R_o/(R_{cq,zn}+R_o) - R_c/(R_{cq,zn}+R_c) \\
&- R_a/(R_{cq,zn}+R_a)) + \beta 2R_{cq}R_{zn}/(R_{cq}+R_{zn})
\end{aligned}
$$

$$\begin{aligned}
\frac{dR_{o,c,a}}{dt} &= r\left(1-\frac{N}{N_{max}}\right)(1-\alpha_{o,c,a})R_{o,c,a} + \lambda\rho_R v - R_{o,c,a}(\delta+\delta_{cq}+\delta_{cu}+\delta_{zn}) \\
&+ \beta R_{o,c,a}(-R_{cq,cu,zn}/(R_{o,c,a}+R_{cq,cu,zn}) - R_{cu,zn}/(R_{o,c,a}+R_{cu,zn}) - R_{cq,zn}/(R_{o,c,a}+R_{cq,zn}) \\
&- R_{cq,cu}/(R_{o,c,a}+R_{cq,cu}) - R_{cq}/(R_{o,c,a}+R_{cq}) - R_{cu}/(R_{o,c,a}+R_{cu}) - R_{zn}/(R_{o,c,a}+R_{zn}) + S/(S+R_{o,c,a})) \\
&+ \beta 2(R_{o,c}R_a/(R_{o,c}+R_a) + R_{c,a}R_o/(R_{c,a}+R_o) + R_{o,a}R_c/(R_{o,a}+R_c)) \\
\frac{dR_{o,c,cu}}{dt} &= r\left(1-\frac{N}{N_{max}}\right)(1-\alpha_{o,c,cu})R_{o,c,cu} + \lambda\rho_R v - R_{o,c,cu}(\delta+\delta_{cq}+\delta_{a}+\delta_{zn}) \\
&+ \beta R_{o,c,cu}(-R_{cq,a,zn}/(R_{o,c,cu}+R_{cq,a,zn}) - R_{a,zn}/(R_{o,c,cu}+R_{a,zn}) - R_{cq,zn}/(R_{o,c,cu}+R_{cq,zn}) \\
&- R_{cq,a}/(R_{o,c,cu}+R_{cq,a}) - R_{zn}/(R_{o,c,cu}+R_{zn}) - R_{cq}/(R_{o,c,cu}+R_{cq}) - R_{a}/(R_{o,c,cu}+R_{a}) + S/(S+R_{o,c,cu})) \\
&+ \beta 2(R_{c,cu}R_o/(R_{c,cu}+R_o) + R_{o,c}R_{cu}/(R_{o,c}+R_{cu}) + R_{o,cu}R_c/(R_{o,cu}+R_c)) \\
\frac{dR_{o,c,zn}}{dt} &= r\left(1-\frac{N}{N_{max}}\right)(1-\alpha_{o,c,zn})R_{o,c,zn} + \lambda\rho_R v - R_{o,c,zn}(\delta+\delta_{cq}+\delta_{a}+\delta_{cu}) \\
&+ \beta R_{o,c,zn}(-R_{cq,a,cu}/(R_{o,c,zn}+R_{cq,a,cu}) - R_{a,cu}/(R_{o,c,zn}+R_{a,cu}) - R_{cq,cu}/(R_{o,c,zn}+R_{cq,cu}) \\
&- R_{cq,a}/(R_{o,c,zn}+R_{cq,a}) - R_{cq}/(R_{o,c,zn}+R_{cq}) - R_{cu}/(R_{o,c,zn}+R_{cu}) - R_{a}/(R_{o,c,zn}+R_{a}) + S/(S+R_{o,c,zn})) \\
&+ \beta 2(R_{c,zn}R_o/(R_{c,zn}+R_o) + R_{o,zn}R_c/(R_{o,zn}+R_c) + R_{o,c}R_{zn}/(R_{o,c}+R_{zn})) \\
\frac{dR_{o,a,cu}}{dt} &= r\left(1-\frac{N}{N_{max}}\right)(1-\alpha_{o,a,cu})R_{o,a,cu} + \lambda\rho_R v - R_{o,a,cu}(\delta+\delta_{cq}+\delta_{c}+\delta_{zn}) \\
&+ \beta R_{o,a,cu}(-R_{c,cq,zn}/(R_{o,a,cu}+R_{c,cq,zn}) - R_{c,zn}/(R_{o,a,cu}+R_{c,zn}) - R_{cq,zn}/(R_{o,a,cu}+R_{cq,zn}) \\
&- R_{c,cq}/(R_{o,a,cu}+R_{c,cq}) - R_{cq}/(R_{o,a,cu}+R_{cq}) - R_{zn}/(R_{o,a,cu}+R_{zn}) - R_{c}/(R_{o,a,cu}+R_{c}) + S/(S+R_{o,a,cu})) \\
&+ \beta 2(R_{a,cu}R_o/(R_{a,cu}+R_o) + R_{o,a}R_{cu}/(R_{o,a}+R_{cu}) + R_{o,cu}R_a/(R_{o,cu}+R_a)) \\
\frac{dR_{o,a,zn}}{dt} &= r\left(1-\frac{N}{N_{max}}\right)(1-\alpha_{o,a,zn})R_{o,a,zn} + \lambda\rho_R v - R_{o,a,zn}(\delta+\delta_{cq}+\delta_{c}+\delta_{cu}) \\
&+ \beta R_{o,a,zn}(-R_{c,cq,cu}/(R_{o,a,zn}+R_{c,cq,cu}) - R_{c,cu}/(R_{o,a,zn}+R_{c,cu}) - R_{cq,cu}/(R_{o,a,zn}+R_{cq,cu}) \\
&- R_{c,cq}/(R_{o,a,zn}+R_{c,cq}) - R_{cq}/(R_{o,a,zn}+R_{cq}) - R_{cu}/(R_{o,a,zn}+R_{cu}) - R_{c}/(R_{o,a,zn}+R_{c}) + S/(S+R_{o,a,zn})) \\
&+ \beta 2(R_{a,zn}R_o/(R_{a,zn}+R_o) + R_{o,zn}R_a/(R_{o,zn}+R_a) + R_{o,a}R_{zn}/(R_{o,a}+R_{zn})) \\
\frac{dR_{o,cu,zn}}{dt} &= r\left(1-\frac{N}{N_{max}}\right)(1-\alpha_{o,cu,zn})R_{o,cu,zn} + \lambda\rho_R v - R_{o,cu,zn}(\delta+\delta_{cq}+\delta_{a}+\delta_{c}) \\
&+ \beta R_{o,cu,zn}(-R_{c,cq,a}/(R_{o,cu,zn}+R_{c,cq,a}) - R_{c,a}/(R_{o,cu,zn}+R_{c,a}) - R_{c,cq}/(R_{o,cu,zn}+R_{c,cq}) \\
&- R_{cq,a}/(R_{o,cu,zn}+R_{cq,a}) - R_{cq}/(R_{o,cu,zn}+R_{cq}) - R_{a}/(R_{o,cu,zn}+R_{a}) - R_{c}/(R_{o,cu,zn}+R_{c}) + S/(S+R_{o,cu,zn})) \\
&+ \beta 2(R_{cu,zn}R_o/(R_{cu,zn}+R_o) + R_{o,zn}R_{cu}/(R_{o,zn}+R_{cu}) + R_{o,cu}R_{zn}/(R_{o,cu}+R_{zn})) \\
\frac{dR_{c,a,cu}}{dt} &= r\left(1-\frac{N}{N_{max}}\right)(1-\alpha_{c,a,cu})R_{c,a,cu}\delta_o + \lambda\rho_R v - R_{c,a,cu}(\delta+\delta_{cq}+\delta_{zn}) \\
&+ \beta R_{c,a,cu}(-R_{o,cq,zn}/(R_{o,cq,zn}+R_{c,a,cu}) - R_{o,zn}/(R_{c,a,cu}+R_{o,zn}) - R_{cq,zn}/(R_{c,a,cu}+R_{cq,zn}) \\
&- R_{o,cq}/(R_{c,a,cu}+R_{o,cq}) - R_{zn}/(R_{c,a,cu}+R_{zn}) - R_{o}/(R_{c,a,cu}+R_{o}) + S/(S+R_{c,a,cu})) \\
&+ \beta 2(R_{c,cu}R_a/(R_{c,cu}+R_a) + R_{a,cu}R_c/(R_{a,cu}+R_c) + R_{c,a}R_{cu}/(R_{c,a}+R_{cu})) \\
\frac{dR_{c,a,zn}}{dt} &= r\left(1-\frac{N}{N_{max}}\right)(1-\alpha_{c,a,zn})R_{c,a,zn}\delta_o + \lambda\rho_R v - R_{c,a,zn}(\delta+\delta_{cq}+\delta_{cu}) \\
&+ \beta R_{c,a,zn}(-R_{o,cq,cu}/(R_{o,cq,cu}+R_{c,a,zn}) - R_{o,cu}/(R_{c,a,zn}+R_{o,cu}) - R_{cq,cu}/(R_{c,a,zn}+R_{cq,cu}) \\
&- R_{o,cq}/(R_{c,a,zn}+R_{o,cq}) - R_{cq}/(R_{c,a,zn}+R_{cq}) - R_{cu}/(R_{c,a,zn}+R_{cu}) - R_{o}/(R_{c,a,zn}+R_{o}) + S/(S+R_{c,a,zn})) \\
&+ \beta 2(R_{a,zn}R_c/(R_{a,zn}+R_c) + R_{c,zn}R_a/(R_{c,zn}+R_a) + R_{c,a}R_{zn}/(R_{c,a}+R_{zn})) \\
\frac{dR_{c,cu,zn}}{dt} &= r\left(1-\frac{N}{N_{max}}\right)(1-\alpha_{c,cu,zn})R_{c,cu,zn}\delta_o + \lambda\rho_R v - R_{c,cu,zn}(\delta+\delta_{cq}+\delta_{a}) \\
&+ \beta R_{c,cu,zn}(-R_{o,cq,a}/(R_{o,cq,a}+R_{c,cu,zn}) - R_{o,a}/(R_{c,cu,zn}+R_{o,a}) - R_{o,cq}/(R_{c,cu,zn}+R_{o,cq}) \\
&- R_{cq,a}/(R_{c,cu,zn}+R_{cq,a}) - R_{cq}/(R_{c,cu,zn}+R_{cq}) - R_{a}/(R_{c,cu,zn}+R_{a}) - R_{o}/(R_{c,cu,zn}+R_{o}) + S/(S+R_{c,cu,zn})) \\
&+ \beta 2(R_{c,zn}R_{cu}/(R_{c,zn}+R_{cu}) + R_{cu,zn}R_c/(R_{cu,zn}+R_c) + R_{c,cu}R_{zn}/(R_{c,cu}+R_{zn}))
\end{aligned}$$

$$
\begin{aligned}
\frac{dR_{a,cu,zn}}{dt} &= r\left(1-\frac{N}{N_{max}}\right)(1-\alpha_{a,cu,zn})R_{a,cu,zn}\delta_o + \lambda\rho_R v - R_{a,cu,zn}(\delta+\delta_{cq}+\delta_c)\\
&+ \beta R_{a,cu,zn}(-R_{o,c,cq}/(R_{o,c,cq}+R_{a,cu,zn}) - R_{o,c}/(R_{a,cu,zn}+R_{o,c}) - R_{o,cq}/(R_{a,cu,zn}+R_{o,cq})\\
&- R_{c,cq}/(R_{a,cu,zn}+R_{c,cq}) + S/(S+R_{a,cu,zn}) - R_{cq}/(R_{a,cu,zn}+R_{cq}) - R_c/(R_{a,cu,zn}+R_c) - R_o/(R_{a,cu,zn}+R_o))\\
&+ \beta 2(R_{cu,zn}R_a/(R_{cu,zn}+R_a) + R_{a,zn}R_{cu}/(R_{a,zn}+R_{cu}) + R_{a,cu}R_{zn}/(R_{a,cu}+R_{zn}))\\
\frac{dR_{c,cq,a}}{dt} &= r\left(1-\frac{N}{N_{max}}\right)(1-\alpha_{c,cq,a})R_{c,cq,a}\delta_o + \lambda\rho_R v - R_{c,cq,a}(\delta+\delta_{cu}+\delta_{zn})\\
&+ \beta R_{c,cq,a}(-R_{o,cu,zn}/(R_{o,cu,zn}+R_{c,cq,a}) - R_{cu,zn}/(R_{c,cq,a}+R_{cu,zn}) - R_{o,zn}/(R_{c,cq,a}+R_{o,zn})\\
&- R_{o,cu}/(R_{c,cq,a}+R_{o,cu}) - R_{zn}/(R_{c,cq,a}+R_{zn}) - R_{cu}/(R_{c,cq,a}+R_{cu}) - R_o/(R_{c,cq,a}+R_o) + S/(S+R_{c,cq,a}))\\
&+ \beta 2(R_{c,a}R_{cq}/(R_{c,a}+R_{cq}) + R_{cq,a}R_c/(R_{cq,a}+R_c) + R_{c,cq}R_a/(R_{c,cq}+R_a))\\
\frac{dR_{o,cq,a}}{dt} &= r\left(1-\frac{N}{N_{max}}\right)(1-\alpha_{o,cq,a})R_{o,cq,a} + \lambda\rho_R v - R_{o,cq,a}(\delta+\delta_{cu}+\delta_{zn})\\
&+ \beta R_{o,cq,a}(-R_{c,cu,zn}/(R_{o,cq,a}+R_{c,cu,zn}) - R_{cu,zn}/(R_{o,cq,a}+R_{cu,zn}) + S/(S+R_{o,cq,a}) - R_{c,zn}/(R_{o,cq,a}+R_{c,zn})\\
&- R_{c,cu}/(R_{o,cq,a}+R_{c,cu}) - R_c/(R_{o,cq,a}+R_c) - R_{zn}/(R_{o,cq,a}+R_{zn}) - R_{cu}/(R_{o,cq,a}+R_{cu}))\\
&+ \beta 2(R_{cq,a}R_{cq}/(R_{cq,a}+R_o) + R_{o,a}R_{cq}/(R_{o,a}+R_{cq}) + R_{o,cq}R_a/(R_{o,cq}+R_a))\\
\frac{dR_{cq,a,cu}}{dt} &= r\left(1-\frac{N}{N_{max}}\right)(1-\alpha_{cq,a,cu})R_{cq,a,cu}\delta_o + \lambda\rho_R v - R_{cq,a,cu}(\delta+\delta_{zn})\\
&+ \beta R_{cq,a,cu}(-R_{o,c,zn}/(R_{o,c,zn}+R_{cq,a,cu}) - R_{c,zn}/(R_{cq,a,cu}+R_{c,zn}) + S/(S+R_{cq,a,cu}) - R_{o,c}/(R_{cq,a,cu}+R_{o,c})\\
&- R_c/(R_{cq,a,cu}+R_c) - R_{cq}/(R_{c,a,cu}+R_{cq}) - R_{zn}/(R_{cq,a,cu}+R_{zn}) - R_o/(R_{cq,a,cu}+R_o)\\
&+ \beta 2(R_{cq,cu}R_a/(R_{cq,cu}+R_a) + R_{a,cu}R_{cq}/(R_{a,cu}+R_{cq}) + R_{cq,a}R_{cu}/(R_{cq,a}+R_{cu}))\\
\frac{dR_{cq,a,zn}}{dt} &= r\left(1-\frac{N}{N_{max}}\right)(1-\alpha_{cq,a,zn})R_{cq,a,zn}\delta_o + \lambda\rho_R v - R_{cq,a,zn}(\delta+\delta_{cu})\\
&\beta R_{cq,cu,zn}(-R_{o,c,a}/(R_{o,c,a}+R_{cq,cu,zn}) - R_{o,cu}/(R_{cq,a,zn}+R_{o,cu}) - R_{c,cu}/(R_{cq,a,zn}+R_{c,cu})\\
&- R_{o,c}/(R_{cq,a,zn}+R_{o,c}) - R_o/(R_{cq,a,zn}+R_o) - R_c/(R_{cq,a,zn}+R_c) - R_{cu}/(R_{cq,a,zn}+R_{cu}) + S/(S+R_{cq,a,zn}))\\
&+ \beta 2(R_{cq,zn}R_a/(R_{cq,zn}+R_a) + R_{a,zn}R_{cq}/(R_{a,zn}+R_{cq}) + R_{cq,a}R_{zn}/(R_{cq,a}+R_{zn}))\\
\frac{dR_{o,c,cq}}{dt} &= r\left(1-\frac{N}{N_{max}}\right)(1-\alpha_{o,c,cq})R_{o,c,cq} + \lambda\rho_R v - R_{o,c,cq}(\delta+\delta_a+\delta_{cu}+\delta_{zn})\\
&+ \beta R_{o,c,cq}(-R_{a,cu,zn}/(R_{o,c,cq}+R_{a,cu,zn}) - R_{cu,zn}/(R_{o,c,cq}+R_{cu,zn}) - R_{a,zn}/(R_{o,c,cq}+R_{a,zn})\\
&- R_{a,cu}/(R_{o,c,cq}+R_{a,cu}) - R_a/(R_{o,c,cq}+R_a) - R_{cu}/(R_{o,c,cq}+R_{cu}) - R_{zn}/(R_{o,c,cq}+R_{zn}) + S/(S+R_{o,c,cq}))\\
&+ \beta 2(R_{o,cq}R_c/(R_{o,cq}+R_c) + R_{c,cq}R_o/(R_{c,cq}+R_o) + R_{o,c}R_{cq}/(R_{o,c}+R_{cq}))\\
\frac{dR_{c,cq,cu}}{dt} &= r\left(1-\frac{N}{N_{max}}\right)(1-\alpha_{c,cq,cu})R_{c,cq,cu}\delta_o + \lambda\rho_R v - R_{c,cq,cu}(\delta+\delta_a+\delta_{zn})\\
&+ \beta R_{c,cq,cu}(-R_{o,a,zn}/(R_{o,a,zn}+R_{c,cq,cu}) - R_{o,zn}/(R_{c,cq,cu}+R_{o,zn}) - R_{a,zn}/(R_{c,cq,cu}+R_{a,zn})\\
&- R_{o,a}/(R_{c,cq,cu}+R_{o,a}) - R_a/(R_{c,cq,cu}+R_a) - R_o/(R_{c,cq,cu}+R_o) + S/(S+R_{c,cq,cu}))\\
&+ \beta 2(R_{c,cu}R_{cq}/(R_{c,cu}+R_{cq}) + R_{cq,cu}R_c/(R_{cq,cu}+R_c) + R_{c,cq}R_{cu}/(R_{c,cq}+R_{cu}))\\
\frac{dR_{c,cq,zn}}{dt} &= r\left(1-\frac{N}{N_{max}}\right)(1-\alpha_{c,cq,zn})R_{c,cq,zn}\delta_o + \lambda\rho_R v - R_{c,cq,zn}(\delta+\delta_a+\delta_{cu})\\
&+ \beta R_{c,cq,zn}(-R_{o,a,cu}/(R_{o,a,cu}+R_{c,cq,zn}) - R_{o,cu}/(R_{c,cq,zn}+R_{o,cu}) - R_{a,cu}/(R_{c,cq,zn}+R_{a,cu})\\
&- R_{o,a}/(R_{c,cq,zn}+R_{o,a}) - R_{cu}/(R_{o,cq,zn}+R_{cu}) - R_{zn}/(R_{c,cq,cu}+R_{zn}) - R_{cu}/(R_{c,cq,zn}+R_{cu})\\
&- R_a/(R_{c,cq,zn}+R_a) - R_o/(R_{c,cq,zn}+R_o) + S/(S+R_{c,cq,zn}))\\
&+ \beta 2(R_{c,zn}R_{cq}/(R_{c,zn}+R_{cq}) + R_{cq,zn}R_c/(R_{cq,zn}+R_c) + R_{c,cq}R_{zn}/(R_{c,cq}+R_{zn}))\\
\frac{dR_{o,cq,cu}}{dt} &= r\left(1-\frac{N}{N_{max}}\right)(1-\alpha_{o,cq,cu})R_{o,cq,cu} + \lambda\rho_R v - R_{o,cq,cu}(\delta+\delta_a+\delta_{zn})\\
&+ \beta R_{o,cq,cu}(-R_{c,a,zn}/(R_{o,cq,cu}+R_{c,a,zn}) - R_{c,zn}/(R_{o,cq,cu}+R_{c,zn}) + S/(S+R_{o,cq,cu}) - R_{a,zn}/(R_{o,cq,cu}+R_{a,zn})\\
&- R_{c,a}/(R_{o,cq,cu}+R_{c,a}) - R_{zn}/(R_{o,cq,cu}+R_{zn}) - R_c/(R_{o,cq,cu}+R_c) - R_a/(R_{o,cq,cu}+R_a))\\
&+ \beta 2(R_{o,cu}R_{cq}/(R_{o,cu}+R_{cq}) + R_{cq,cu}R_o/(R_{cq,cu}+R_o) + R_{o,cq}R_{cu}/(R_{o,cq}+R_{cu}))
\end{aligned}
$$

$$
\begin{aligned}
\frac{dR_{o,cq,zn}}{dt} &= r\left(1-\frac{N}{N_{max}}\right)(1-\alpha_{o,cq,zn})R_{o,cq,zn} + \lambda\rho_R v - R_{o,cq,zn}(\delta+\delta_a+\delta_{cu}) \\
&+ \beta R_{o,cq,zn}(-R_{c,a,cu}/(R_{o,cq,zn}+R_{c,a,cu}) - R_{c,cu}/(R_{o,cq,zn}+R_{c,cu}) - R_{a,cu}/(R_{o,cq,zn}+R_{a,cu}) \\
&- R_{c,a}/(R_{o,cq,zn}+R_{c,a}) - R_c/(R_{o,cq,zn}+R_c) + S/(S+R_{o,cq,zn}) - R_a/(R_{o,cq,zn}+R_a)) \\
&+ \beta 2(R_{cq,zn}R_o/(R_{cq,zn}+R_o) + R_{o,zn}R_{cq}/(R_{o,zn}+R_{cq}) + R_{o,cq}R_{zn}/(R_{o,cq}+R_{zn}))
\end{aligned}
$$

$$
\begin{aligned}
\frac{dR_{cq,cu,zn}}{dt} &= r\left(1-\frac{N}{N_{max}}\right)(1-\alpha_{cq,cu,zn})R_{cq,cu,zn}\delta_o + \lambda\rho_R v - R_{cq,cu,zn}(\delta+\delta_a) \\
&+ \beta R_{cq,cu,zn}(-R_{o,c,a}/(R_{o,c,a}+R_{cq,cu,zn}) - R_{o,c}/(R_{cq,cu,zn}+R_{o,c}) - R_{o,a}/(R_{cq,cu,zn}+R_{o,a}) \\
&- R_{c,a}/(R_{cq,cu,zn}+R_{c,a}) - R_c/(R_{cq,cu,zn}+R_c) - R_o/(R_{cq,cu,zn}+R_o) - R_a/(R_{cq,cu,zn}+R_a) + S/(S+R_{cq,cu,zn})) \\
&+ \beta 2(R_{cq,zn}R_{cu}/(R_{cq,zn}+R_{cu}) + R_{cu,zn}R_{cq}/(R_{cu,zn}+R_{cq}) + R_{cq,cu}R_{zn}/(R_{cq,cu}+R_{zn}))
\end{aligned}
$$

$$
\begin{aligned}
\frac{dR_{o,c,a,cu}}{dt} &= r\left(1-\frac{N}{N_{max}}\right)(1-\alpha_{o,c,a,cu})R_{o,c,a,cu} + \lambda\rho_R v - R_{o,c,a,cu}(\delta+\delta_{cq}+\delta_{zn}) \\
&+ \beta R_{o,c,a,cu}(-R_{cq,zn}/(R_{o,c,a,cu}+R_{cq,zn}) - R_{zn}/(R_{o,c,a,cu}+R_{zn}) - R_{cq}/(R_{o,c,a,cu}+R_{cq}) + S/(S+R_{o,c,a,cu})) \\
&+ \beta 2(R_{o,cu}R_{c,a}/(R_{o,cu}+R_{c,a}) + R_{o,a}R_{c,cu}/(R_{o,a}+R_{c,cu}) + R_{o,c}R_{a,cu}/(R_{o,c}+R_{a,cu}) \\
&+ R_{c,a,cu}R_o/(R_{c,a,cu}+R_o) + R_{o,a,cu}R_c/(R_{o,a,cu}+R_c) + R_{o,c,cu}R_a/(R_{o,c,cu}+R_a) + R_{o,c,a}R_{cu}/(R_{o,c,a}+R_{cu}))
\end{aligned}
$$

$$
\begin{aligned}
\frac{dR_{o,c,a,zn}}{dt} &= r\left(1-\frac{N}{N_{max}}\right)(1-\alpha_{o,c,a,zn})R_{o,c,a,zn} + \lambda\rho_R v - R_{o,c,a,zn}(\delta+\delta_{cq}+\delta_{cu}) \\
&+ \beta R_{o,c,a,zn}(-R_{cq,cu}/(R_{o,c,a,zn}+R_{cq,cu}) - R_{cu}/(R_{o,c,a,zn}+R_{cu}) - R_{cq}/(R_{o,c,a,zn}+R_{cq}) + S/(S+R_{o,c,a,zn})) \\
&+ \beta 2(R_{o,zn}R_{c,a}/(R_{o,zn}+R_{c,a}) + R_{o,a}R_{c,zn}/(R_{o,a}+R_{c,zn}) + R_{o,c}R_{a,zn}/(R_{o,c}+R_{a,zn}) \\
&+ R_{c,a,zn}R_o/(R_{c,a,zn}+R_o) + R_{o,a,zn}R_c/(R_{o,a,zn}+R_c) + R_{o,c,zn}R_a/(R_{o,c,zn}+R_a) + R_{o,c,a}R_{zn}/(R_{o,c,a}+R_{zn}))
\end{aligned}
$$

$$
\begin{aligned}
\frac{dR_{o,c,cu,zn}}{dt} &= r\left(1-\frac{N}{N_{max}}\right)(1-\alpha_{o,c,cu,zn})R_{o,c,cu,zn} + \lambda\rho_R v - R_{o,c,cu,zn}(\delta+\delta_{cq}+\delta_a) \\
&+ \beta R_{o,c,cu,zn}(-R_{cq,a}/(R_{o,c,cu,zn}+R_{cq,a}) - R_a/(R_{o,c,cu,zn}+R_a) - R_{cq}/(R_{o,c,cu,zn}+R_{cq}) + S/(S+R_{o,c,cu,zn}) \\
&+ \beta 2(R_{o,zn}R_{c,cu}/(R_{o,zn}+R_{c,cu}) + R_{o,cu}R_{c,zn}/(R_{o,cu}+R_{c,zn}) + R_{o,c}R_{cu,zn}/(R_{o,c}+R_{cu,zn}) \\
&+ R_{c,cu,zn}R_o/(R_{c,cu,zn}+R_o) + R_{o,cu,zn}R_c/(R_{o,cu,zn}+R_c) + R_{o,c,zn}R_{cu}/(R_{o,c,zn}+R_{cu}) + R_{o,c,cu}R_{zn}/(R_{o,c,cu}+R_{zn})
\end{aligned}
$$

$$
\begin{aligned}
\frac{dR_{o,a,cu,zn}}{dt} &= r\left(1-\frac{N}{N_{max}}\right)(1-\alpha_{o,a,cu,zn})R_{o,a,cu,zn} + \lambda\rho_R v - R_{o,a,cu,zn}(\delta+\delta_{cq}+\delta_c) \\
&+ \beta R_{o,a,cu,zn}(-R_{c,cq}/(R_{o,a,cu,zn}+R_{c,cq}) - R_c/(R_{o,a,cu,zn}+R_c) - R_{cq}/(R_{o,a,cu,zn}+R_{cq}) + S/(S+R_{o,a,cu,zn})) \\
&+ \beta 2(R_{o,a,cu}R_{zn}/(R_{o,a,cu}+R_{zn}) + R_{o,a,zn}R_{cu}/(R_{o,a,zn}+R_{cu}) + R_{o,cu,zn}R_a/(R_{o,cu,zn}+R_a) \\
&+ R_{a,cu,zn}R_o/(R_{a,cu,zn}+R_o) + R_{o,a}R_{cu,zn}/(R_{o,a}+R_{cu,zn}) + R_{o,cu}R_{a,zn}/(R_{o,cu}+R_{a,zn}) \\
&+ R_{o,zn}R_{a,cu}/(R_{o,zn}+R_{a,cu}))
\end{aligned}
$$

$$
\begin{aligned}
\frac{dR_{c,a,cu,zn}}{dt} &= r\left(1-\frac{N}{N_{max}}\right)(1-\alpha_{c,a,cu,zn})R_{c,a,cu,zn}\delta_o + \lambda\rho_R v - R_{c,a,cu,zn}(\delta+\delta_{cq}) \\
&+ \beta R_{c,a,cu,zn}(-R_{o,cq}/(R_{c,a,cu,zn}+R_{o,cq}) - R_o/(R_{c,a,cu,zn}+R_o) - R_{cq}/(R_{c,a,cu,zn}+R_{cq}) + S/(S+R_{c,a,cu,zn})) \\
&+ \beta 2(R_{c,zn}R_{a,cu}/(R_{c,zn}+R_{a,cu}) + R_{c,cu}R_{a,zn}/(R_{c,cu}+R_{a,zn}) + R_{c,a}R_{cu,zn}/(R_{c,a}+R_{cu,zn}) \\
&+ R_{c,a,cu}R_{zn}/(R_{c,a,cu}+R_{zn}) + R_{c,a,zn}R_{cu}/(R_{c,a,zn}+R_{cu}) + R_{c,cu,zn}R_a/(R_{c,cu,zn}+R_a) + R_{a,cu,zn}R_c/(R_{a,cu,zn}+R_c))
\end{aligned}
$$

$$
\begin{aligned}
\frac{dR_{o,cq,a,cu}}{dt} &= r\left(1-\frac{N}{N_{max}}\right)(1-\alpha_{o,cq,a,cu})R_{o,cq,a,cu} + \lambda\rho_R v - R_{o,cq,a,cu}(\delta+\delta_{zn}) \\
&+ \beta R_{o,cq,a,cu}(-R_{c,zn}/(R_{o,cq,a,cu}+R_{c,zn}) - R_{zn}/(R_{o,cq,a,cu}+R_{zn}) - R_c/(R_{o,cq,a,cu}+R_c) + S/(S+R_{o,cq,a,cu})) \\
&+ \beta 2(R_{o,cq,a}R_{cu}/(R_{o,cq,a}+R_{cu}) + R_{o,cq,cu}R_a/(R_{o,cq,cu}+R_a) + R_{o,a,cu}R_{cq}/(R_{o,a,cu}+R_{cq}) \\
&+ R_{cq,a,cu}R_o/(R_{cq,a,cu}+R_o) + R_{o,cq}R_{a,cu}/(R_{o,cq}+R_{a,cu}) + R_{o,a}R_{cq,cu}/(R_{o,a}+R_{cq,cu}) \\
&+ R_{o,cu}R_{cq,a}/(R_{o,cu}+R_{cq,a})
\end{aligned}
$$

$$
\begin{aligned}
\frac{dR_{o,cq,a,zn}}{dt} &= r\left(1-\frac{N}{N_{max}}\right)(1-\alpha_{o,cq,a,zn})R_{o,cq,a,zn} + \lambda\rho_R v - R_{o,cq,a,zn}(\delta+\delta_{cu}) \\
&+ \beta R_{o,cq,a,zn}(-R_{c,cu}/(R_{o,cq,a,zn}+R_{c,cu}) - R_{cu}/(R_{o,cq,a,zn}+R_{cu}) - R_c/(R_{o,cq,a,zn}+R_c) + S/(S+R_{o,cq,a,zn})) \\
&+ \beta 2(R_{o,cq,a}R_{zn}/(R_{o,cq,a}+R_{zn}) + R_{o,cq,zn}R_a/(R_{o,cq,zn}+R_a) + R_{o,a,zn}R_{cq}/(R_{o,a,zn}+R_{cq}) \\
&+ R_{cq,a,zn}R_o/(R_{cq,a,zn}+R_o) + R_{o,cq}R_{a,zn}/(R_{o,cq}+R_{a,zn}) + R_{cq,zn}R_{o,a}/(R_{cq,zn}+R_{o,a}) \\
&+ R_{cq,a}R_{o,zn}/(R_{cq,a}+R_{o,zn})) \\
\frac{dR_{o,c,cq,zn}}{dt} &= r\left(1-\frac{N}{N_{max}}\right)(1-\alpha_{o,c,cq,zn})R_{o,c,cq,zn} + \lambda\rho_R v - R_{o,c,cq,zn}(\delta+\delta_{cu}+\delta_a) \\
&+ \beta R_{o,c,cq,zn}(-R_{a,cu}/(R_{o,c,cq,zn}+R_{a,cu}) - R_{cu}/(R_{o,c,cq,zn}+R_{cu}) - R_a/(R_{o,c,cq,zn}+R_a) + S/(S+R_{o,c,cq,zn})) \\
&+ \beta 2(R_{zn}R_{o,c,cq}/(R_{zn}+R_{o,c,cq}) + R_{cq}R_{o,c,zn}/(R_{cq}+R_{o,c,zn}) + R_c R_{o,cq,zn}/(R_c+R_{o,cq,zn}) \\
&+ R_o R_{c,cq,zn}/(R_o+R_{c,cq,zn}) + R_{o,c}R_{cq,zn}/(R_{o,c}+R_{cq,zn}) + R_{o,cq}R_{c,zn}/(R_{o,cq}+R_{c,zn}) \\
&+ R_{c,cq}R_{o,zn}/(R_{c,cq}+R_{o,zn}) \\
\frac{dR_{o,c,cq,cu}}{dt} &= r\left(1-\frac{N}{N_{max}}\right)(1-\alpha_{o,c,cq,cu})R_{o,c,cq,cu} + \lambda\rho_R v - R_{o,c,cq,cu}(\delta+\delta_{zn}+\delta_a) \\
&+ \beta R_{o,c,cq,cu}(-R_{a,zn}/(R_{o,c,cq,cu}+R_{a,zn}) - R_{zn}/(R_{o,c,cq,cu}+R_{zn}) - R_a/(R_{o,c,cq,cu}+R_a) + S/(S+R_{o,c,cq,cu})) \\
&+ \beta 2(R_{o,cu}R_{c,cq}/(R_{o,cu}+R_{c,cq}) + R_{o,cq}R_{c,cu}/(R_{o,cq}+R_{c,cu}) + R_{o,c}R_{cq,cu}/(R_{o,c}+R_{cq,cu}) \\
&+ R_{o,c,cq}R_{cu}/(R_{o,c,cq}+R_{cu}) + R_{o,c,cu}R_{cq}/(R_{o,c,cu}+R_{cq}) + R_{o,cq,cu}R_c/(R_{o,cq,cu}+R_c) \\
&+ R_{c,cq,cu}R_o/(R_{c,cq,cu}+R_o)) \\
\frac{dR_{cq,a,cu,zn}}{dt} &= r\left(1-\frac{N}{N_{max}}\right)(1-\alpha_{cq,a,cu,zn})R_{cq,a,cu,zn}\delta_o + \lambda\rho_R v - R_{cq,a,cu,zn}(\delta) \\
&+ \beta R_{cq,a,cu,zn}(-R_{o,c}/(R_{cq,a,cu,zn}+R_{o,c}) - R_o/(R_{cq,a,cu,zn}+R_o) - R_c/(R_{cq,a,cu,zn}+R_c) + S/(S+R_{cq,a,cu,zn})) \\
&+ \beta 2(R_{cq,a,cu}R_{zn}/(R_{cq,a,cu}+R_{zn}) + R_{cq,a,zn}R_{cu}/(R_{cq,a,zn}+R_{cu}) + R_{cq,cu,zn}R_a/(R_{cq,cu,zn}+R_a) \\
&+ R_{a,cu,zn}R_{cq}/(R_{a,cu,zn}+R_{cq}) + R_{cq,a}R_{cu,zn}/(R_{cq,a}+R_{cu,zn}) + R_{cq,cu}R_{a,zn}/(R_{cq,cu}+R_{a,zn}) \\
&+ R_{cq,zn}R_{a,cu}/(R_{cq,zn}+R_{a,cu})) \\
\frac{dR_{o,c,cq,a}}{dt} &= r\left(1-\frac{N}{N_{max}}\right)(1-\alpha_{o,c,cq,a})R_{o,c,cq,a} + \lambda\rho_R v - R_{o,c,cq,a}(\delta+\delta_{zn}+\delta_{cu}) \\
&+ \beta R_{o,c,cq,a}(-R_{cu,zn}/(R_{o,c,cq,a}+R_{cu,zn}) - R_{zn}/(R_{o,c,cq,a}+R_{zn}) - R_{cu}/(R_{o,c,cq,a}+R_{cu}) + S/(S+R_{o,c,cq,a})) \\
&+ \beta 2(R_{o,c,cq}R_a/(R_{o,c,cq}+R_a) + R_{o,c,a}R_{cq}/(R_{o,c,a}+R_{cq}) + R_{o,cq,a}R_c/(R_{o,cq,a}+R_c) \\
&+ R_{o,cq,a}R_c/(R_{o,cq,a}+R_c) + R_{c,cq,a}R_o/(R_{c,cq,a}+R_o) + R_{cq,a}R_{o,c}/(R_{cq,a}+R_{o,c}) \\
&+ R_{c,a}R_{o,cq}/(R_{c,a}+R_{o,cq}) + R_{c,cq}R_{o,a}/(R_{c,cq}+R_{o,a})) \\
\frac{dR_{c,cq,a,cu}}{dt} &= r\left(1-\frac{N}{N_{max}}\right)(1-\alpha_{c,cq,a,cu})R_{c,cq,a,cu}\delta_o + \lambda\rho_R v - R_{c,cq,a,cu}(\delta+\delta_{zn}) \\
&+ \beta R_{c,cq,a,cu}(-R_{o,zn}/(R_{c,cq,a,cu}+R_{o,zn}) - R_{zn}/(R_{c,cq,a,cu}+R_{zn}) - R_o/(R_{c,cq,a,cu}+R_o) + S/(S+R_{c,cq,a,cu})) \\
&+ \beta 2(R_{c,cu}R_{cq,a}/(R_{c,cu}+R_{cq,a}) + R_{c,a}R_{cq,cu}/(R_{c,a}+R_{cq,cu}) + R_{c,cq}R_{a,cu}/(R_{c,cq}+R_{a,cu}) \\
&+ R_{c,cq,a}R_{cu}/(R_{c,cq,a}+R_{cu}) + R_{c,cq,cu}R_a/(R_{c,cq,cu}+R_a) + R_{c,a,cu}R_{cq}/(R_{c,a,cu}+R_{cq}) \\
&+ R_{cq,a,cu}R_c/(R_{cq,a,cu}+R_c) \\
\frac{dR_{c,cq,a,zn}}{dt} &= r\left(1-\frac{N}{N_{max}}\right)(1-\alpha_{c,cq,a,zn})R_{c,cq,a,zn}\delta_o + \lambda\rho_R v - R_{c,cq,a,zn}(\delta+\delta_{cu}) \\
&+ \beta R_{c,cq,a,zn}(-R_{o,cu}/(R_{c,cq,a,zn}+R_{o,cu}) - R_{cu}/(R_{c,cq,a,zn}+R_{cu}) - R_o/(R_{c,cq,a,zn}+R_o) + S/(S+R_{c,cq,a,zn})) \\
&+ \beta 2(R_{c,cq,a}R_{zn}/(R_{c,cq,a}+R_{zn}) + R_{c,cq,zn}R_a/(R_{c,cq,zn}+R_a) + R_{c,a,zn}R_{cq}/(R_{c,a,zn}+R_{cq}) \\
&+ R_{cq,a,zn}R_c/(R_{cq,a,zn}+R_c) + R_{c,cq}R_{a,zn}/(R_{c,cq}+R_{a,zn}) + R_{c,a}R_{cq,zn}/(R_{c,a}+R_{cq,zn}) \\
&+ R_{c,zn}R_{cq,a}/(R_{c,zn}+R_{cq,a}))
\end{aligned}
$$

$$
\begin{aligned}
\frac{dR_{c,cq,cu,zn}}{dt} &= r\left(1-\frac{N}{N_{max}}\right)(1-\alpha_{c,cq,cu,zn})R_{c,cq,cu,zn}\delta_o + \lambda\rho_R v - R_{c,cq,cu,zn}(\delta+\delta_a) \\
&+ \beta R_{c,cq,cu,zn}(-R_{o,a}/(R_{c,cq,cu,zn}+R_{o,a}) - R_o/(R_{c,cq,cu,zn}+R_o) - R_a/(R_{c,cq,cu,zn}+R_a) + S/(S+R_{c,cq,cu,zn})) \\
&+ \beta 2(R_{cu}R_{c,cq,zn}/(R_{cu}+R_{c,cq,zn}) + R_{zn}R_{c,cq,cu}/(R_{zn}+R_{c,cq,cu}) + R_{cq}R_{c,cu,zn}/(R_{cq}+R_{c,cu,zn}) \\
&+ R_c R_{cq,cu,zn}/(R_c+R_{cq,cu,zn}) + R_{c,cq}R_{cu,zn}/(R_{c,cq}+R_{cu,zn}) + R_{c,cu}R_{cq,zn}/(R_{c,cu}+R_{cq,zn}) \\
&+ R_{c,zn}R_{cq,cu}/(R_{c,zn}+R_{cq,cu})) \\
\frac{dR_{o,cq,cu,zn}}{dt} &= r\left(1-\frac{N}{N_{max}}\right)(1-\alpha_{o,cq,cu,zn})R_{o,cq,cu,zn} + \lambda\rho_R v - R_{o,cq,cu,zn}(\delta+\delta_a) \\
&+ \beta R_{o,cq,cu,zn}(-R_{c,a}/(R_{o,cq,cu,zn}+R_{c,a}) - R_c/(R_{o,cq,cu,zn}+R_c) - R_a/(R_{o,cq,cu,zn}+R_a) + S/(S+R_{o,cq,cu,zn})) \\
&+ \beta 2(R_{cq,cu}R_{o,zn}/(R_{cq,cu}+R_{o,zn}) + R_{cq,zn}R_{o,cu}/(R_{cq,zn}+R_{o,cu}) + R_{cu,zn}R_{o,cq}/(R_{cu,zn}+R_{o,cq}) \\
&+ R_{o,cq,cu}R_{zn}/(R_{o,cq,cu}+R_{zn}) + R_{o,cq,zn}R_{cu}/(R_{o,cq,zn}+R_{cu}) + R_{o,cu,zn}R_{cq}/(R_{o,cu,zn}+R_{cq}) \\
&+ R_{cq,cu,zn}R_o/(R_{cq,cu,zn}+R_o)) \\
\frac{dR_{o,c,a,cu,zn}}{dt} &= r\left(1-\frac{N}{N_{max}}\right)(1-\alpha_{o,c,a,cu,zn})R_{o,c,a,cu,zn} + \lambda\rho_R v - R_{o,c,a,cu,zn}(\delta+\delta_{cq}) \\
&+ \beta R_{o,c,a,cu,zn}(-R_{cq}/(R_{o,c,a,cu,zn}+R_{cq}) + S/(S+R_{o,c,a,cu,zn})) + \beta 2(R_{a,cu,zn}R_{o,c}/(R_{a,cu,zn}+R_{o,c}) \\
&+ R_{o,cu,zn}R_{c,a}/(R_{o,cu,zn}+R_{c,a}) + R_{c,cu,zn}R_{o,a}/(R_{c,cu,zn}+R_{o,a}) + R_{o,c,cu}R_{a,zn}/(R_{o,c,cu}+R_{a,zn}) \\
&+ R_{o,a,cu}R_{c,zn}/(R_{o,a,cu}+R_{c,zn}) + R_{c,a,cu}R_{o,zn}/(R_{c,a,cu}+R_{o,zn}) + R_{o,c,a}R_{cu,zn}/(R_{o,c,a}+R_{cu,zn}) \\
&+ R_{c,a,zn}R_{o,cu}/(R_{c,a,zn}+R_{o,cu}) + R_{o,a,zn}R_{c,cu}/(R_{o,a,zn}+R_{c,cu}) + R_{o,c,zn}R_{a,cu}/(R_{o,c,zn}+R_{a,cu}) \\
&+ R_{o,c,a,cu}R_{zn}/(R_{o,c,a,cu}+R_{zn}) + R_{c,a,cu,zn}R_o/(R_{c,a,cu,zn}+R_o) + R_{o,a,cu,zn}R_c/(R_{o,a,cu,zn}+R_c) \\
&+ R_{o,c,cu,zn}R_a/(R_{o,c,cu,zn}+R_a) + R_{o,c,a,zn}R_{cu}/(R_{o,c,a,zn}+R_{cu})) \\
\frac{dR_{o,c,cq,a,cu}}{dt} &= r\left(1-\frac{N}{N_{max}}\right)(1-\alpha_{o,c,cq,a,cu})R_{o,c,cq,a,cu} + \lambda\rho_R v - R_{o,c,cq,a,cu}(\delta+\delta_{zn}) \\
&+ \beta R_{o,c,cq,a,cu}(-R_{zn}/(R_{o,c,cq,a,cu}+R_{zn}) + S/(S+R_{o,c,cq,a,cu})) + \beta 2(R_{o,a,cu}R_{c,cq}/(R_{o,a,cu}+R_{c,cq}) \\
&+ R_{o,c,cu}R_{cq,a}/(R_{o,c,cu}+R_{cq,a}) + R_{o,cq,cu}R_{c,a}/(R_{o,cq,cu}+R_{c,a}) + R_{c,cq,cu}R_{o,a}/(R_{c,cq,cu}+R_{o,a}) \\
&+ R_{c,a,cu}R_{o,cq}/(R_{c,a,cu}+R_{o,cq}) + R_{cq,a,cu}R_{o,c}/(R_{cq,a,cu}+R_{o,c}) + R_{c,cq,a}R_{o,cu}/(R_{c,cq,a}+R_{o,cu}) \\
&+ R_{o,cq,a}R_{c,cu}/(R_{o,cq,a}+R_{c,cu}) + R_{o,c,a}R_{cq,cu}/(R_{o,c,a}+R_{cq,cu}) + R_{c,cq,a,cu}R_o/(R_{c,cq,a,cu}+R_o) \\
&+ R_{o,c,cq}R_{a,cu}/(R_{o,c,cq}+R_{a,cu}) + R_{o,cq,a,cu}R_c/(R_{o,cq,a,cu}+R_c) + R_{o,c,a,cu}R_{cq}/(R_{o,c,a,cu}+R_{cq}) \\
&+ R_{o,c,cq,a}R_{cu}/(R_{o,c,cq,a}+R_{cu}) + R_{o,c,cq,cu}R_a/(R_{o,c,cq,cu}+R_a)) \\
\frac{dR_{o,c,cq,a,zn}}{dt} &= r\left(1-\frac{N}{N_{max}}\right)(1-\alpha_{o,c,cq,a,zn})R_{o,c,cq,a,zn} + \lambda\rho_R v - R_{o,c,cq,a,zn}(\delta+\delta_{cu}) \\
&+ \beta R_{o,c,cq,a,zn}(-R_{cu}/(R_{o,c,cq,a,zn}+R_{cu}) + S/(S+R_{o,c,cq,a,zn})) + \beta 2(R_{o,a,zn}R_{c,cq}/(R_{o,a,zn}+R_{c,cq}) \\
&+ R_{o,c,zn}R_{cq,a}/(R_{o,c,zn}+R_{cq,a}) + R_{o,cq,zn}R_{c,a}/(R_{o,cq,zn}+R_{c,a}) + R_{c,cq,zn}R_{o,a}/(R_{c,cq,zn}+R_{o,a}) \\
&+ R_{c,a,zn}R_{o,cq}/(R_{c,a,zn}+R_{o,cq}) + R_{cq,a,zn}R_{o,c}/(R_{cq,a,zn}+R_{o,c}) + R_{c,cq,a}R_{o,zn}/(R_{c,cq,a}+R_{o,zn}) \\
&+ R_{o,cq,a}R_{c,zn}/(R_{o,cq,a}+R_{c,zn}) + R_{o,c,a}R_{cq,zn}/(R_{o,c,a}+R_{cq,zn}) + R_{o,c,cq}R_{a,zn}/(R_{o,c,cq}+R_{a,zn}) \\
&+ R_{c,cq,a,zn}R_o/(R_{c,cq,a,zn}+R_o) + R_{o,cq,a,zn}R_c/(R_{o,cq,a,zn}+R_c) + R_{o,c,a,zn}R_{cq}/(R_{o,c,a,zn}+R_{cq}) \\
&+ R_{o,c,cq,zn}R_a/(R_{o,c,cq,zn}+R_a) + R_{o,c,cq,a}R_{zn}/(R_{o,c,cq,a}+R_{zn})) \\
\frac{dR_{c,cq,a,cu,zn}}{dt} &= r\left(1-\frac{N}{N_{max}}\right)(1-\alpha_{c,cq,a,cu,zn})\delta_o R_{c,cq,a,cu,zn} + \lambda\rho_R v - R_{c,cq,a,cu,zn}(\delta) \\
&+ \beta R_{c,cq,a,cu,zn}(-R_o/(R_{c,cq,a,cu,zn}+R_o) + S/(S+R_{c,cq,a,cu,zn})) + \beta 2(R_{a,cu,zn}R_{c,cq}/(R_{a,cu,zn}+R_{c,cq}) \\
&+ R_{c,cu,zn}R_{cq,a}/(R_{c,cu,zn}+R_{cq,a}) + R_{cq,cu,zn}R_{c,a}/(R_{cq,cu,zn}+R_{c,a}) + R_{c,cq,cu}R_{a,zn}/(R_{c,cq,cu}+R_{a,zn}) \\
&+ R_{c,a,cu}R_{cq,zn}/(R_{c,a,cu}+R_{cq,zn}) + R_{cq,a,cu}R_{c,zn}/(R_{cq,a,cu}+R_{c,zn}) + R_{c,cq,a}R_{cu,zn}/(R_{c,cq,a}+R_{cu,zn}) \\
&+ R_{cq,a,zn}R_{c,cu}/(R_{cq,a,zn}+R_{c,cu}) + R_{c,a,zn}R_{cq,cu}/(R_{c,a,zn}+R_{cq,cu}) + R_{c,cq,zn}R_{a,cu}/(R_{c,cq,zn}+R_{a,cu}) \\
&+ R_{c,cq,a,cu}R_c/(R_{c,cq,a,cu}+R_{zn}) + R_{cq,a,cu,zn}R_c/(R_{cq,a,cu,zn}+R_c) + R_{c,a,cu,zn}R_{cq}/(R_{c,a,cu,zn}+R_{cq}) \\
&+ R_{c,cq,cu,zn}R_a/(R_{c,cq,cu,zn}+R_a) + R_{c,cq,a,zn}R_{cu}/(R_{c,cq,a,zn}+R_{cu})
\end{aligned}
$$

$$
\begin{aligned}
\frac{dR_{o,cq,a,cu,zn}}{dt} &= r\left(1-\frac{N}{N_{max}}\right)(1-\alpha_{o,cq,a,cu,zn})R_{o,cq,a,cu,zn} + \lambda\rho_R v - R_{o,cq,a,cu,zn}(\delta) \\
&+ \beta R_{o,cq,a,cu,zn}(-R_c/(R_{o,cq,a,cu,zn}+R_c) + S/(S+R_{o,cq,a,cu,zn})) + \beta 2(R_{a,cu,zn}R_{o,cq}/(R_{a,cu,zn}+R_{o,cq}) \\
&+ R_{o,cu,zn}R_{cq,a}/(R_{o,cu,zn}+R_{cq,a}) + R_{cq,cu,zn}R_{o,a}/(R_{cq,cu,zn}+R_{o,a}) + R_{o,cq,cu}R_{a,zn}/(R_{o,cq,cu}+R_{a,zn}) \\
&+ R_{o,a,cu}R_{cq,zn}/(R_{o,a,cu}+R_{cq,zn}) + R_{cq,a,cu}R_{o,zn}/(R_{cq,a,cu}+R_{o,zn}) + R_{o,cq,a}R_{cu,zn}/(R_{o,cq,a}+R_{cu,zn}) \\
&+ R_{cq,a,zn}R_{o,cu}/(R_{cq,a,zn}+R_{o,cu}) + R_{o,a,zn}R_{cq,cu}/(R_{o,a,zn}+R_{cq,cu}) + R_{o,cq,zn}R_{a,cu}/(R_{o,cq,zn}+R_{a,cu}) \\
&+ R_{o,cq,a,cu}R_{zn}/(R_{o,cq,a,cu}+R_{zn}) + R_{cq,a,cu,zn}R_{o}/(R_{cq,a,cu,zn}+R_{o}) + R_{o,a,cu,zn}R_{cq}/(R_{o,a,cu,zn}+R_{cq}) \\
&+ R_{o,cq,cu,zn}R_{a}/(R_{o,cq,cu,zn}+R_{a}) + R_{o,cq,a,zn}R_{cu}/(R_{o,cq,a,zn}+R_{cu})) \\
\frac{dR_{o,c,cq,cu,zn}}{dt} &= r\left(1-\frac{N}{N_{max}}\right)(1-\alpha_{o,c,cq,cu,zn})R_{o,c,cq,cu,zn} + \lambda\rho_R v - R_{o,c,cq,cu,zn}(\delta) \\
&+ \beta R_{o,c,cq,cu,zn}(-R_a/(R_{o,c,cq,cu,zn}+R_a) + S/(S+R_{o,c,cq,cu,zn})) + \beta 2(R_{o,cu,zn}R_{c,cq}/(R_{o,cu,zn}+R_{c,cq}) \\
&+ R_{o,c,zn}R_{cq,cu}/(R_{o,c,zn}+R_{cq,cu}) + R_{o,cq,zn}R_{c,cu}/(R_{o,cq,zn}+R_{c,cu}) + R_{c,cq,zn}R_{o,cu}/(R_{c,cq,zn}+R_{o,cu}) \\
&+ R_{c,cu,zn}R_{o,cq}/(R_{c,cu,zn}+R_{o,cq}) + R_{cq,cu,zn}R_{o,c}/(R_{cq,cu,zn}+R_{o,c}) + R_{c,cq,cu}R_{o,zn}/(R_{c,cq,cu}+R_{o,zn}) \\
&+ R_{o,cq,cu}R_{c,zn}/(R_{o,cq,cu}+R_{c,zn}) + R_{o,c,cu}R_{cq,zn}/(R_{o,c,cu}+R_{cq,zn}) + R_{o,c,cq}R_{cu,zn}/(R_{o,c,cq}+R_{cu,zn}) \\
&+ R_{c,cq,cu,zn}R_{o}/(R_{c,cq,cu,zn}+R_{o}) + R_{o,cq,cu,zn}R_{c}/(R_{o,cq,cu,zn}+R_{c}) + R_{o,c,cu,zn}R_{cq}/(R_{o,c,cu,zn}+R_{cq}) \\
&+ R_{o,c,cq,zn}R_{cu}/(R_{o,c,cq,zn}+R_{cu}) + R_{o,c,cq,cu}R_{zn}/(R_{o,c,cq,cu}+R_{zn})) \\
\frac{dR_{o,c,cq,a,cu,zn}}{dt} &= r\left(1-\frac{N}{N_{max}}\right)(1-\alpha_{o,c,cq,a,cu,zn})R_{o,c,cq,a,cu,zn} + \lambda\rho_R v - R_{o,c,cq,a,cu,zn}(\delta) \\
&+ \beta 2(R_{o,cq,a}R_{c,cu,zn}/(R_{o,cq,a}+R_{c,cu,zn}) + R_{o,cq,cu}R_{c,a,zn}/(R_{o,cq,cu}+R_{c,a,zn}) \\
&+ R_{o,cq,zn}R_{c,a,cu}/(R_{o,cq,zn}+R_{c,a,cu}) + R_{o,a,cu}R_{c,cq,zn}/(R_{o,a,cu}+R_{c,cq,zn}) + R_{o,a,zn}R_{c,cq,cu}/(R_{o,a,zn}+R_{c,cq,cu}) \\
&+ R_{o,cu,zn}R_{c,cq,a}/(R_{o,cu,zn}+R_{c,cq,a}) + R_{o,c,cq}R_{a,cu,zn}/(R_{o,c,cq}+R_{a,cu,zn}) + R_{o,c,a}R_{cq,cu,zn}/(R_{o,c,a}+R_{cq,cu,zn}) \\
&+ R_{o,c,cu}R_{cq,a,zn}/(R_{o,c,cu}+R_{cq,a,zn}) + R_{o,c,zn}R_{cq,a,cu}/(R_{o,c,zn}+R_{cq,a,cu}) + R_{cq,a,cu,zn}R_{o,c}/(R_{cq,a,cu,zn}+R_{o,c}) \\
&+ R_{c,a,cu,zn}R_{o,cq}/(R_{c,a,cu,zn}+R_{o,cq}) + R_{c,cq,cu,zn}R_{o,a}/(R_{c,cq,cu,zn}+R_{o,a}) + R_{c,cq,a,zn}R_{o,cu}/(R_{c,cq,a,zn}+R_{o,cu}) \\
&+ R_{c,cq,a,cu}R_{o,zn}/(R_{c,cq,a,cu}+R_{o,zn}) + R_{o,a,cu,zn}R_{c,cq}/(R_{o,a,cu,zn}+R_{c,cq}) + R_{o,cq,cu,zn}R_{c,a}/(R_{o,cq,cu,zn}+R_{c,a}) \\
&+ R_{o,cq,a,zn}R_{c,cu}/(R_{o,cq,a,zn}+R_{c,cu}) + R_{o,cq,a,cu}R_{c,zn}/(R_{o,cq,a,cu}+R_{c,zn}) + R_{o,c,cu,zn}R_{cq,a}/(R_{o,c,cu,zn}+R_{cq,a}) \\
&+ R_{o,c,a,zn}R_{cq,cu}/(R_{o,c,a,zn}+R_{cq,cu}) + R_{o,c,a,cu}R_{cq,zn}/(R_{o,c,a,cu}+R_{cq,zn}) + R_{o,c,cq,zn}R_{a,cu}/(R_{o,c,cq,zn}+R_{a,cu}) \\
&+ R_{o,c,cq,cu}R_{a,zn}/(R_{o,c,cq,cu}+R_{a,zn}) + R_{o,c,cq,a}R_{cu,zn}/(R_{o,c,cq,a}+R_{cu,zn}) + R_{o,cq,a,cu,zn}R_{c}/(R_{o,cq,a,cu,zn}+R_{c}) \\
&+ R_{o,c,a,cu,zn}R_{cq}/(R_{o,c,a,cu,zn}+R_{cq}) + R_{o,c,cq,cu,zn}R_{a}/(R_{o,c,cq,cu,zn}+R_{a}) + R_{o,c,cq,a,zn}R_{cu}/(R_{o,c,cq,a,zn}+R_{cu}) \\
&+ R_{o,c,cq,a,cu}R_{zn}/(R_{o,c,cq,a,cu}+R_{zn})) + \beta R_{o,c,cq,a,cu,zn}S/(S+R_{o,c,cq,a,cu,zn}) \\
\frac{dA_{oxy}}{dt} &= -\gamma_{oxy}A_{Oxy} \\
\frac{dA_{cex}}{dt} &= -\gamma_{cex}A_{Cex} \\
\frac{dA_{amox}}{dt} &= -\gamma_{amox}A_{Amox} \\
\frac{dA_{cefq}}{dt} &= -\gamma_{cef}Cef \\
\frac{dV}{dt} &= \lambda
\end{aligned}
$$

$$
\begin{aligned}
N_{max} &= N_{max_conc} * V \\
Cu &= copper * V \\
Zn &= zinc * V \\
\delta_o &= 1 - 2\frac{(A_{Oxy}/V)^2}{1 + (A_{Oxy}/V)^2} \\
\delta_a &= \frac{A_{Amox}/V)^2}{(8^2 + (A_{Amox}/V)^2} \\
\delta_c &= \frac{(A_{cex}/V)^2}{(8^2 + (A_{cex}/V)^2)} \\
\delta_{cq} &= \frac{(A_{cefq}/V)^2}{(0.06^2 + (A_{cefq}/V)^2} \\
\delta_{cu} &= \frac{(Cu/V)^3.306794}{(103.6884^3.306794 + (Cu/V)^3.306794)} \\
\delta_{zn} &= \frac{(Zn/V)^0.888219}{(1205.832^0.888219 + (Zn/V)^0.888219)}
\end{aligned}
$$

### Supplementary Text 3: Supplementary Methods for additional data sets used for model calibration

#### Analysis of antibiotics by liquid chromatography-tandem mass spectrometry

Samples (2 mL) were prepared for liquid chromatography-tandem mass spectrometry (LC-MS/MS) using solid phase extraction (SAX, HLB cartridges). Extracts were separated by gradient elution reversed-phase chromatography (Shimadzu, Milton Keynes, UK, Gemini C18, 100 mm x 2.0 mm and 3 µm particle size) at a flow rate of 250 $\mu L.min^{-1}$. The mobile phases consisted of water with 1% ammonium acetate and 0.2% formic acid at pH 3 (A) and acetonitrile (B). LC-MS/MS analysis was in positive electrospray ionisation mode on a SCIEX Q-TRAP 4000 quadrupole linear ion-trap mass spectrometer (Applied Biosystems, Foster City, CA, USA). Identification of antibiotics in samples was performed based on retention time, the known ratio between the two MRM transitions employed for each antibiotic and reference standards. Collected ion spectra data were analysed using Analyst software (AB SCIEX). Quantification was performed using 11-point calibration lines using deuterated internal standards. Data were accepted for further statistical analysis if the relative standard deviation (RSD) of the specific internal quality control samples was <15%.

#### Filtered slurry experiment to estimate growth rates

150mL of slurry was filtered using a decontaminated slurry filter, centrifuged at 13300 rpm for 10 mins, filter sterilized twice using 0.45 μm syringe filters, stored at 4°C before use.

The experiment was as follows:

1. control (sterile filtrate, no inoculum)
2. filtrate + e.coli ATCC25922
3. filtrate + e.coli slurry isolate 1212-883
4. filtrate + e.coli ATCC25922 + 1% UHT milk
5. filtrate + e.coli slurry isolate 1212-883 + 1 % UHT milk
6. filtrate + e.coli ATCC25922 + 1% glucose
7. filtrate + e.coli slurry isolate 1212-883 + 1 % glucose.

The flasks were placed at 37°C for 24 hours and a time series of Miles and Misra plates were taken at T=0, T = 1 (hour), T = 4 and T=24. 0.01mL were triplicate spotted onto MacConkey plates. Data is not shown; estimated growth rates are in Supplementary Text 1 and Table S7.

**Supplementary Table S1**: dates of sampling from main tank for microbiology and water quality analyses

| **Sample Number** | **Month** | **Date** |
|---|---|---|
| 1 | May | 16/05/2017 |
| 2 | May | 22/05/2017 |
| 3 | June | 29/06/2017 |
| 4 | July | 11/07/2017 |
| 5 | July | 18/07/2017 |
| 6 | July | 25/07/2017 |
| 7 | August | 01/08/2017 |
| 8 | August | 16/08/2017 |
| 9 | August | 22/08/2017 |
| 10 | September | 05/09/2017 |
| 11 | September | 22/09/2017 |
| 12 | September | 27/09/2017 |
| 13 | October | 10/10/2017 |
| 14 | October | 17/10/2017 |
| 15 | October | 31/10/2017 |
| 16 | November | 14/11/2017 |
| 17 | November | 21/11/2017 |

**Supplementary Table S2:** Antibiotic discs used for AST assays.

| **Antibiotic Name** | **Antibiotic Class** | **Abbreviation** | **Concentration (mg)** |
|---|---|---|---|
| Ampicillin | Penicillins | AM | 10 |
| Amoxicillin Clavulanic Acid | B-Lactamase Inhibitor combination | AMC | 20 & 10 |
| Cefoxitin | Second Gen Ceph | FOX | 30 |
| Ceftazidime | Third Gen Ceph | CAZ | 30 |
| Cefotaxime | Third Gen Ceph | CTX | 30 |
| Cefpodoxime Protexil | Third Gen Ceph | CPD | 10 |
| Imipenem | Carbapenem | IPM | 10 |
| Nalidixic Acid | Quinolones | NA | 30 |
| Ciprofloxacin | Fluoroquinolones | CIP | 5 |
| Chloramphenicol | Phenicols | C | 30 |
| Aztreonam | Monobactam | ATM | 30 |
| Streptomycin | Aminoglycosides | S10 | 10 |
| Tetracycline | Tetracyclines | TE | 30 |
| Azithromycin | Macrolide | AZM | 10 |
| Nitrofurantoin | Nitrofurans | F | 300 |
| Trimethoprim-sulfamethoxazole | Folate Pathway Inhibitors | SXT | 1.25 & 23.75 |

**Supplementary Table S3:** Annual on-farm antibiotic use 2015-2017

| Product Name | Doses 2015 | Doses 2016 | Doses 2017 | Active Ingredients | Antibiotic class | Quantity |
|---|---|---|---|---|---|---|
| Alamycin | 0 | 39 | 80 | Oxytetracycline Hydrochloride | Tetracycline | 100mg/ml |
| Alamycin LA | 5 | 109 | 0 | Oxytetracycline Dihydrate | Tetracycline | 200mg/ml |
| Betamox | 3 | 24 | 74 | Amoxicillin | Beta-lactam | 150mg/ml |
| Bimotrim | 0 | 19 | 0 | Sulfadoxine | Sulfonamide | 200mg/ml |
| | | | | Trimethoprim | Dihydrofolate reductase inhibitor | 40mg/ml |
| Cephaguard DC | 24 | 0 | 0 | Cefquinome (as sulphate) | Beta-lactam (4th generation cephalosporin) | 3g/ syringe |
| Ceporex | 5 | 373 | 65 | Cefalexin sodium equivalent to Cefalexin | Beta-lactam (1st generation cephalosporin) | 180mg/ml |
| Engemycin DD | 112 | 0 | 0 | Oxytetracycline (as hydrochloride) | Tetracycline | 100mg/ml |
| Hexasol LA | 0 | 2 | 0 | Oxytetracycline (as dihydrate) | Tetracycline | 300mg/ml |
| | | | | Flunixin (as flunixin meglumin) | NA (anti-inflammatory) | 20mg/ml |
| Naxcel | 63 | 1 | 0 | Ceftiofur (as crystalline free acid) | Beta-lactam (3rd generation cephalosporin) | 200mg/ml |
| Orbenin Dry Cow | 0 | 1 | 0 | Cloxacillin benzathine | Beta-lactam | 500mg/dose |
| Orbenin Extra Dry Cow | 45 | 80 | 33 | Cloxacillin benzathine | Beta-lactam | 600mg/syringe |
| Tetra-Delta | 64 | 909 | 261 | Novobiocin Sodium equal to Novobiocin | Aminocoumarin | 100mg/dose |
| | | | | Neomycin Sulphate equal to Neomycin | Aminoglycoside | 105mg/dose |
| | | | | Procaine Penicillin | Beta-lactam | 100mg/dose |
| | | | | Dihydrostreptomycin Sulphate | Aminoglycoside | 100mg/dose |
| | | | | Prednisolone | NA (anti-inflammatory) | 10mg/dose |
| Ubro Yellow Milking Cow | 225 | 113 | 0 | Penethamate Hydriodide (diethylaminoethyl ester of benzylpenicillin) | Beta-lactam (prodrug) | 150mg/dose |
| | | | | Dihydrostreptomycin Sulphate | Aminoglycoside | 185mg/dose |
| | | | | Framycetin Sulphate | Aminoglycoside | 50mg/dose |
| | | | | Prednisolone | NA (anti-inflammatory) | 5mg/dose |
| Ultrapen LA | 2 | 22 | 10 | Procaine Benzylpenicillin | Beta-lactam | 300mg/ml |

**Supplementary Table S4:** Bacterial Phyla detected in the slurry tank (%age frequencies)

| **Phylum** | **01/08/2017** | **05/09/2017** | **07/06/2017** | **10/10/2017** | **11/07/2017** |
|---|---|---|---|---|---|
| unclassified | 53.2 | 52.4 | 56.7 | 54.8 | 55.6 |
| Bacteroidetes (Bacteroidota) | 14.6 | 13.4 | 13.3 | 13.7 | 13.8 |
| Firmicutes (Bacillota) | 14.1 | 14.6 | 12.7 | 12.9 | 14.0 |
| Proteobacteria (Pseudomonadota) | 4.21 | 7.18 | 3.63 | 5.80 | 3.32 |
| Spirochaetes | 2.70 | 2.20 | 3.39 | 2.95 | 2.86 |
| cannot be assigned to a (non-viral) phylum | 2.26 | 2.24 | 2.05 | 2.08 | 2.04 |
| Euryarchaeota | 1.92 | 1.99 | 1.57 | 1.79 | 2.03 |
| Tenericutes | 1.53 | 0.94 | 1.92 | 2.20 | 0.79 |
| Actinobacteria (Actinomycetota) | 0.74 | 0.78 | 0.64 | 0.67 | 0.79 |
| Synergistetes | 0.72 | 0.53 | 0.85 | 0.10 | 0.83 |
| Candidatus Cloacimonetes | 0.50 | 0.56 | 0.27 | 0.41 | 0.63 |
| Lentisphaerae | 0.31 | 0.27 | 0.25 | 0.20 | 0.36 |
| Chloroflexi | 0.28 | 0.35 | 0.23 | 0.19 | 0.33 |
| Fibrobacteres | 0.22 | 0.26 | 0.15 | 0.18 | 0.25 |
| Candidatus Riflebacteria | 0.19 | 0.16 | 0.35 | 0.22 | 0.07 |
| Verrucomicrobia | 0.22 | 0.21 | 0.16 | 0.16 | 0.23 |
| Planctomycetes | 0.20 | 0.19 | 0.16 | 0.14 | 0.22 |
| Cyanobacteria | 0.13 | 0.13 | 0.11 | 0.11 | 0.12 |
| Viruses | 0.10 | 0.11 | 0.11 | 0.13 | 0.10 |
| Candidatus Falkowbacteria | 0.125 | 0.070 | 0.150 | 0.097 | 0.085 |
| Fusobacteria | 0.090 | 0.085 | 0.088 | 0.089 | 0.079 |
| Ignavibacteriae | 0.088 | 0.089 | 0.072 | 0.075 | 0.083 |
| Elusimicrobia | 0.088 | 0.058 | 0.048 | 0.055 | 0.073 |
| Acidobacteria | 0.068 | 0.068 | 0.056 | 0.054 | 0.073 |
| Thermotogae | 0.059 | 0.054 | 0.056 | 0.054 | 0.054 |
| Chlamydiae | 0.057 | 0.054 | 0.056 | 0.058 | 0.051 |
| Nitrospirae | 0.053 | 0.051 | 0.044 | 0.041 | 0.054 |
| Candidatus Omnitrophica | 0.051 | 0.057 | 0.041 | 0.042 | 0.051 |
| Ascomycota | 0.049 | 0.048 | 0.043 | 0.044 | 0.051 |
| Candidatus Moranbacteria | 0.033 | 0.051 | 0.059 | 0.043 | 0.026 |
| Chlorobi | 0.042 | 0.041 | 0.039 | 0.037 | 0.041 |
| Candidatus Saccharibacteria | 0.050 | 0.030 | 0.040 | 0.027 | 0.043 |
| Armatimonadetes | 0.039 | 0.037 | 0.033 | 0.027 | 0.045 |
| Candidatus Nomurabacteria | 0.062 | 0.030 | 0.011 | 0.025 | 0.037 |
| Candidatus Parcubacteria | 0.051 | 0.027 | 0.017 | 0.024 | 0.029 |
| Candidatus Melainabacteria | 0.033 | 0.029 | 0.026 | 0.028 | 0.027 |

| | | | | | |
|---|---|---|---|---|---|
| Kiritimatiellaeota | 0.033 | 0.030 | 0.021 | 0.015 | 0.043 |
| Deinococcus-Thermus | 0.023 | 0.022 | 0.020 | 0.019 | 0.023 |
| Basidiomycota | 0.022 | 0.022 | 0.019 | 0.021 | 0.022 |
| Gemmatimonadetes | 0.020 | 0.019 | 0.015 | 0.015 | 0.020 |
| Candidatus Magasanikbacteria | 0.021 | 0.015 | 0.018 | 0.016 | 0.016 |
| Candidatus Peregrinibacteria | 0.019 | 0.016 | 0.014 | 0.015 | 0.016 |
| Aquificae | 0.017 | 0.016 | 0.014 | 0.015 | 0.016 |
| Candidatus Uhrbacteria | 0.018 | 0.013 | 0.014 | 0.013 | 0.015 |
| candidate division Zixibacteria | 0.016 | 0.015 | 0.013 | 0.014 | 0.016 |
| Candidatus Marinimicrobia | 0.015 | 0.015 | 0.012 | 0.014 | 0.015 |
| Candidatus Aminicenantes | 0.016 | 0.015 | 0.012 | 0.011 | 0.017 |
| Balneolaeota | 0.015 | 0.014 | 0.012 | 0.013 | 0.013 |
| Deferribacteres | 0.013 | 0.013 | 0.012 | 0.012 | 0.013 |
| Candidatus Shapirobacteria | 0.015 | 0.011 | 0.006 | 0.010 | 0.019 |
| Candidatus Kaiserbacteria | 0.015 | 0.010 | 0.008 | 0.009 | 0.011 |
| Candidatus Roizmanbacteria | 0.012 | 0.010 | 0.009 | 0.010 | 0.011 |
| Chytridiomycota | 0.010 | 0.010 | 0.008 | 0.010 | 0.010 |
| Candidatus Atribacteria | 0.010 | 0.009 | 0.010 | 0.009 | 0.010 |
| Candidatus Bathyarchaeota | 0.011 | 0.010 | 0.008 | 0.008 | 0.011 |
| Chlorophyta | 0.010 | 0.010 | 0.009 | 0.010 | 0.010 |
| Candidatus Margulisbacteria | 0.010 | 0.012 | 0.005 | 0.009 | 0.009 |
| Candidatus Woesearchaeota | 0.011 | 0.009 | 0.007 | 0.008 | 0.009 |
| Crenarchaeota | 0.010 | 0.009 | 0.008 | 0.008 | 0.009 |
| candidate division WWE3 | 0.010 | 0.009 | 0.007 | 0.007 | 0.008 |
| Nitrospinae | 0.009 | 0.009 | 0.008 | 0.007 | 0.009 |
| Mucoromycota | 0.008 | 0.008 | 0.008 | 0.008 | 0.008 |
| Candidatus Wallbacteria | 0.008 | 0.008 | 0.010 | 0.008 | 0.007 |
| Candidatus Woesebacteria | 0.009 | 0.008 | 0.007 | 0.007 | 0.008 |
| Thermodesulfobacteria | 0.008 | 0.008 | 0.007 | 0.007 | 0.008 |
| Candidatus Taylorbacteria | 0.012 | 0.006 | 0.004 | 0.004 | 0.010 |
| Candidatus Nealsonbacteria | 0.011 | 0.007 | 0.005 | 0.006 | 0.006 |
| Candidatus Rokubacteria | 0.008 | 0.008 | 0.006 | 0.006 | 0.008 |
| Candidatus Desantisbacteria | 0.008 | 0.007 | 0.006 | 0.005 | 0.008 |
| Apicomplexa | 0.007 | 0.006 | 0.006 | 0.007 | 0.007 |
| candidate division WOR-3 | 0.007 | 0.006 | 0.006 | 0.006 | 0.006 |
| Candidatus Buchananbacteria | 0.007 | 0.005 | 0.007 | 0.006 | 0.006 |
| Calditrichaeota | 0.007 | 0.007 | 0.005 | 0.005 | 0.007 |
| Candidatus Gottesmanbacteria | 0.007 | 0.006 | 0.005 | 0.005 | 0.006 |
| Candidatus Yanofskybacteria | 0.007 | 0.005 | 0.005 | 0.005 | 0.006 |
| Rhodothermaeota | 0.006 | 0.006 | 0.005 | 0.006 | 0.006 |
| Thaumarchaeota | 0.007 | 0.006 | 0.005 | 0.005 | 0.005 |
| Candidatus Raymondbacteria | 0.0059 | 0.0060 | 0.0050 | 0.0043 | 0.0062 |

| | | | | | |
|---|---|---|---|---|---|
| Candidatus Zambryskibacteria | 0.0088 | 0.0044 | 0.0027 | 0.0038 | 0.0062 |
| Candidatus Berkelbacteria | 0.0059 | 0.0046 | 0.0048 | 0.0047 | 0.0049 |
| Candidatus Wolfebacteria | 0.0069 | 0.0047 | 0.0037 | 0.0043 | 0.0053 |
| Candidatus Levybacteria | 0.0055 | 0.0045 | 0.0043 | 0.0045 | 0.0048 |
| Candidatus Diapherotrites | 0.0091 | 0.0040 | 0.0013 | 0.0027 | 0.0056 |
| Candidatus Latescibacteria | 0.0048 | 0.0045 | 0.0039 | 0.0034 | 0.0050 |
| Dictyoglomi | 0.0045 | 0.0043 | 0.0043 | 0.0040 | 0.0043 |
| Candidatus Komeilibacteria | 0.0050 | 0.0036 | 0.0045 | 0.0040 | 0.0038 |
| Candidatus Portnoybacteria | 0.0054 | 0.0039 | 0.0035 | 0.0036 | 0.0041 |
| Candidatus Kerfeldbacteria | 0.0050 | 0.0036 | 0.0041 | 0.0038 | 0.0039 |
| Candidatus Doudnabacteria | 0.0047 | 0.0038 | 0.0039 | 0.0037 | 0.0039 |
| Candidatus Firestonebacteria | 0.0044 | 0.0042 | 0.0034 | 0.0033 | 0.0046 |
| Candidatus Schekmanbacteria | 0.0040 | 0.0037 | 0.0036 | 0.0043 | 0.0042 |
| Candidatus Campbellbacteria | 0.0063 | 0.0033 | 0.0026 | 0.0029 | 0.0045 |
| candidate division NC10 | 0.0043 | 0.0040 | 0.0034 | 0.0030 | 0.0048 |
| Candidatus Pacebacteria | 0.0044 | 0.0037 | 0.0033 | 0.0035 | 0.0042 |
| Candidatus Delongbacteria | 0.0038 | 0.0036 | 0.0036 | 0.0036 | 0.0033 |
| Bacillariophyta | 0.0039 | 0.0032 | 0.0033 | 0.0037 | 0.0036 |
| Candidatus Tectomicrobia | 0.0037 | 0.0037 | 0.0032 | 0.0030 | 0.0039 |
| Candidatus Handelsmanbacteria | 0.0037 | 0.0036 | 0.0031 | 0.0025 | 0.0045 |
| Candidatus Micrarchaeota | 0.0046 | 0.0031 | 0.0027 | 0.0029 | 0.0037 |
| Candidatus Kryptonia | 0.0037 | 0.0035 | 0.0031 | 0.0033 | 0.0033 |
| Chrysiogenetes | 0.0035 | 0.0038 | 0.0030 | 0.0035 | 0.0033 |
| Candidatus Daviesbacteria | 0.0040 | 0.0033 | 0.0027 | 0.0031 | 0.0036 |
| Candidatus Kuenenbacteria | 0.0041 | 0.0028 | 0.0035 | 0.0033 | 0.0030 |
| Candidatus Hydrogenedentes | 0.0035 | 0.0034 | 0.0031 | 0.0027 | 0.0040 |
| Candidatus Fermentibacteria | 0.0035 | 0.0033 | 0.0030 | 0.0027 | 0.0039 |
| Candidatus Staskawiczbacteria | 0.0049 | 0.0030 | 0.0025 | 0.0026 | 0.0031 |
| Candidatus Giovannonibacteria | 0.0041 | 0.0025 | 0.0025 | 0.0025 | 0.0028 |
| Candidatus Lloydbacteria | 0.0047 | 0.0026 | 0.0016 | 0.0021 | 0.0033 |
| Caldiserica | 0.0031 | 0.0028 | 0.0028 | 0.0028 | 0.0026 |
| Candidatus Goldbacteria | 0.0027 | 0.0037 | 0.0019 | 0.0027 | 0.0026 |
| Candidatus Glassbacteria | 0.0028 | 0.0027 | 0.0024 | 0.0023 | 0.0029 |
| Candidatus Gracilibacteria | 0.0033 | 0.0027 | 0.0021 | 0.0025 | 0.0023 |
| Candidatus Harrisonbacteria | 0.0033 | 0.0021 | 0.0021 | 0.0022 | 0.0023 |
| Candidatus Edwardsbacteria | 0.0026 | 0.0024 | 0.0022 | 0.0020 | 0.0028 |
| Candidatus Yonathbacteria | 0.0039 | 0.0019 | 0.0014 | 0.0016 | 0.0027 |
| Candidatus Vogelbacteria | 0.0037 | 0.0019 | 0.0013 | 0.0018 | 0.0027 |
| Candidatus Jorgensenbacteria | 0.0026 | 0.0021 | 0.0020 | 0.0021 | 0.0023 |
| Candidatus Collierbacteria | 0.0027 | 0.0021 | 0.0021 | 0.0019 | 0.0022 |
| Zoopagomycota | 0.0023 | 0.0021 | 0.0021 | 0.0021 | 0.0023 |
| Candidatus Curtissbacteria | 0.0023 | 0.0020 | 0.0019 | 0.0021 | 0.0023 |

| | | | | | |
|---|---|---|---|---|---|
| Candidatus Lokiarchaeota | 0.0022 | 0.0021 | 0.0019 | 0.0018 | 0.0021 |
| Candidatus Wildermuthbacteria | 0.0027 | 0.0017 | 0.0014 | 0.0017 | 0.0018 |
| Candidatus Terrybacteria | 0.0025 | 0.0016 | 0.0013 | 0.0016 | 0.0021 |
| Candidatus Dadabacteria | 0.0018 | 0.0018 | 0.0017 | 0.0018 | 0.0018 |
| Candidatus Azambacteria | 0.0024 | 0.0017 | 0.0016 | 0.0017 | 0.0017 |
| Candidatus Sungbacteria | 0.0023 | 0.0016 | 0.0015 | 0.0015 | 0.0017 |
| Candidatus Woykebacteria | 0.0020 | 0.0017 | 0.0015 | 0.0015 | 0.0018 |
| Candidatus Ryanbacteria | 0.0020 | 0.0017 | 0.0015 | 0.0014 | 0.0017 |
| Candidatus Acetothermia | 0.0017 | 0.0017 | 0.0016 | 0.0014 | 0.0017 |
| Candidatus Beckwithbacteria | 0.0019 | 0.0015 | 0.0013 | 0.0013 | 0.0017 |
| Candidatus Lindowbacteria | 0.0017 | 0.0016 | 0.0015 | 0.0012 | 0.0017 |
| Candidatus Microgenomates | 0.0019 | 0.0014 | 0.0013 | 0.0013 | 0.0017 |
| Candidatus Amesbacteria | 0.0018 | 0.0016 | 0.0012 | 0.0013 | 0.0018 |
| Candidatus Heimdallarchaeota | 0.0017 | 0.0015 | 0.0014 | 0.0015 | 0.0015 |
| Microsporidia | 0.0014 | 0.0013 | 0.0012 | 0.0014 | 0.0014 |
| Candidatus Thorarchaeota | 0.0015 | 0.0013 | 0.0014 | 0.0011 | 0.0014 |
| Candidatus Liptonbacteria | 0.0019 | 0.0012 | 0.0010 | 0.0011 | 0.0013 |
| Candidatus Fischerbacteria | 0.0013 | 0.0013 | 0.0012 | 0.0011 | 0.0012 |
| Candidatus Coatesbacteria | 0.0012 | 0.0011 | 0.0011 | 0.0011 | 0.0013 |
| candidate division CPR3 | 0.0013 | 0.0011 | 0.0011 | 0.0011 | 0.0011 |
| Candidatus Korarchaeota | 0.0012 | 0.0011 | 0.0011 | 0.0011 | 0.0011 |
| Chromerida | 0.0012 | 0.0011 | 0.0009 | 0.0010 | 0.0013 |
| candidate division CPR2 | 0.0012 | 0.0010 | 0.0011 | 0.0010 | 0.0011 |
| Candidatus Adlerbacteria | 0.0016 | 0.0009 | 0.0006 | 0.0009 | 0.0012 |
| Candidatus Niyogibacteria | 0.0015 | 0.0009 | 0.0008 | 0.0010 | 0.0010 |
| Candidatus Colwellbacteria | 0.0014 | 0.0009 | 0.0008 | 0.0008 | 0.0009 |
| Candidatus Spechtbacteria | 0.0013 | 0.0010 | 0.0008 | 0.0009 | 0.0009 |
| Candidatus Eisenbacteria | 0.0010 | 0.0009 | 0.0008 | 0.0008 | 0.0010 |
| Candidatus Tagabacteria | 0.0014 | 0.0008 | 0.0007 | 0.0007 | 0.0009 |
| Candidatus Wirthbacteria | 0.0010 | 0.0009 | 0.0007 | 0.0006 | 0.0011 |
| Candidatus Aenigmarchaeota | 0.0013 | 0.0009 | 0.0005 | 0.0007 | 0.0010 |
| Eustigmatophyceae | 0.0011 | 0.0008 | 0.0007 | 0.0007 | 0.0009 |
| Candidatus Andersenbacteria | 0.0009 | 0.0007 | 0.0008 | 0.0008 | 0.0008 |
| Blastocladiomycota | 0.0009 | 0.0009 | 0.0006 | 0.0007 | 0.0009 |
| Phaeophyceae | 0.0007 | 0.0007 | 0.0007 | 0.0007 | 0.0008 |
| Cryptomycota | 0.0008 | 0.0008 | 0.0005 | 0.0006 | 0.0007 |
| Candidatus Blackburnbacteria | 0.0007 | 0.0006 | 0.0005 | 0.0005 | 0.0007 |
| Candidatus Aerophobetes | 0.0007 | 0.0006 | 0.0005 | 0.0006 | 0.0006 |
| Candidatus Chisholmbacteria | 0.0007 | 0.0006 | 0.0005 | 0.0004 | 0.0007 |
| Candidatus Abawacabacteria | 0.0007 | 0.0005 | 0.0004 | 0.0005 | 0.0005 |
| candidate division CPR1 | 0.0006 | 0.0005 | 0.0004 | 0.0005 | 0.0005 |
| Candidatus Odinarchaeota | 0.0006 | 0.0005 | 0.0005 | 0.0005 | 0.0005 |

| | | | | | |
|---|---|---|---|---|---|
| candidate division KD3-62 | 0.0005 | 0.0005 | 0.0004 | 0.0004 | 0.0006 |
| Candidatus Nanohaloarchaeota | 0.0004 | 0.0003 | 0.0003 | 0.0003 | 0.0003 |
| Candidatus Fraserbacteria | 0.0003 | 0.0003 | 0.0003 | 0.0003 | 0.0003 |
| Euglenida | 0.0002 | 0.0003 | 0.0002 | 0.0002 | 0.0003 |
| Candidatus Brennerbacteria | 0.0003 | 0.0003 | 0.0001 | 0.0002 | 0.0002 |
| Candidatus Parvarchaeota | 0.0003 | 0.0002 | 0.0002 | 0.0002 | 0.0002 |
| Nanoarchaeota | 0.0002 | 0.0001 | 0.0001 | 0.0001 | 0.0001 |
| Candidatus Jacksonbacteria | 0.0001 | 0.0001 | 0.0001 | 0.0001 | 0.0001 |
| Xanthophyceae | 0.00004 | 0.00002 | 0.00002 | 0.00003 | 0.00002 |
| Candidatus Poribacteria | 0.00003 | 0.00001 | 0.00002 | 0.00001 | 0.00003 |
| Candidatus Veblenbacteria | 0.00003 | 0.00003 | 0.00001 | 0.00001 | 0.00001 |
| Pinguiophyceae | 0.000003 | 0.000001 | 0.000001 | 0.000005 | 0.000002 |
| Colponemidia | 0.000001 | 0.000005 | 0.000001 | 0.000001 | 0.000001 |
| Bolidophyceae | 0.000000 | 0.000001 | 0.000000 | 0.000000 | 0.000001 |
| Picozoa | 0.000001 | 0.000000 | 0.000000 | 0.000001 | 0.000000 |
| Haplosporidia | 0.000001 | 0.000000 | 0.000000 | 0.000000 | 0.000000 |

**Supplementary Table S5:** Model Parameter Values

| Parameter | Definition | Value | Source |
|---|---|---|---|
| $\delta$ | Environmental Death Rate | N(0.0647, 7.28 x $10^{-8}$) | Estimated by MCMC from minitank experiment |
| Cu | Copper level in tank | 22.37 mg $L^{-1}$ | Arya *et al.* 2021 |
| Zn | Zinc level in tank | 32.158 mg $L^{-1}$ | Arya *et al.* 2021 |
| $\alpha_A$ | Amoxicillin resistance fitness cost | 0.05 | Expert judgment |
| $\alpha_{Ceq}$ | Cefquinome resistance fitness cost | logN(-2.942, 0.0261) | Estimated by MCMC from minitank experiment |
| $\alpha_{Cex}$ | Cefalexin resistance fitness cost | logN(-2.942, 0.0261) | Estimated by MCMC from minitank experiment |
| $\alpha_O$ | Oxytetracycline resistance fitness cost | 0.003 | Expert judgment |
| $\alpha_{Cu}$ | Copper resistance fitness cost | logN(-2.498, 6.23 x $10^{-4}$) | Estimated by MCMC from minitank experiment |
| $\alpha_{Zn}$ | Zinc resistance fitness cost | logN(-2.498, 6.23 x $10^{-4}$) | Estimated by MCMC from minitank experiment |
| $\rho$ | Proportion of resistant populations in slurry inflow | logN(-7.227, 0.00957) | Estimated by MCMC from minitank experiment |
| Vi | Initial slurry volume in tank | 1,000,000 litres | On farm observations |
| $\lambda$ | Slurry inflow rate | 1480 L $hour^{-1}$ | Estimated from on farm data |
| r | Bacterial growth rate | 0.08 $hour^{-1}$ | Estimated by MCMC from laboratory experiment |
| $N_{max}$ | Carrying capacity of slurry | 1 x $10^{10}$ cells $L^{-1}$ | Baker *et al.* 2016 |
| $\beta$ | Horizontal gene transfer rate | 0.000001 $hour^{-1}$ | Baker *et al.* 2016 |
| v | *E. coli* in slurry inflow | 2.16 x $10^{8}$ cells $L^{-1}$ | Measured experimentally |
| $\gamma_A$ | Amoxicillin decay rate | 0.00722 $hour^{-1}$ | Längin *et al.* 2008 |
| $\gamma_{Ceq}$ | Cefquinome decay rate | 0.0384 $hour^{-1}$ | Estimated by MCMC from minitank experiment |
| $\gamma_{Cex}$ | Cefalexin decay rate | 0.0176 $hour^{-1}$ | Estimated by MCMC from minitank experiment |
| $\gamma_O$ | Oxytetracycline decay rate | 0.000289 $hour^{-1}$ | Arikan *et al.* 2006 |
| $Tk_f$ | Tank draining every | 60 days | Estimated from on farm data |
| $Tk_0$ | First day of tank draining | 50 days | - |
| $Tk_r$ | Proportion of slurry remaining in tank after draining | 0.1 | Estimated from on farm data |
| $MIC_O$ | Minimum inhibitory concentration of oxytetracycline | 2 | EUCAST for *E. coli* |
| $MIC_A$ | Minimum inhibitory concentration of amoxicillin | 4 | EUCAST for *E. coli* |
| $MIC_{Cex}$ | Minimum inhibitory concentration of cefalexin | 8 | EUCAST for *E. coli* |
| $MIC_{Ceq}$ | Minimum inhibitory concentration of cefquinome | 0.06 | EUCAST for *E. coli* |

**Supplementary Table S6:** Bayes factors for alternative growth rate models supporting use of a constant microbial growth rate.

| Model | Posterior model probabilities |
|---|---|
| **Model 1 – Constant growth and death rates r = 0.08, low nutrients** | **5263.2** |
| Model 2 – Constant growth and death rates 0.183, high nutrients | 5259.7 |
| Model 3 – Variable growth and death rates $\delta = \eta r$, high nutrients | 4986.3 |
| Model 4 – Variable growth and death rates $\delta = \eta r$, low nutrients | 4992.6 |
| Model 5 – Variable growth and death rates $\delta = r - \eta$, high nutrients | 5216.4 |
| Model 6 – Variable growth and death rates $\delta = r - \eta$, low nutrients | 5100.7 |

**References for Supplementary Text**

Arikan OA, Sikora LJ, Mulbry W, Khan SU, Rice C and Foster GD 2006. The fate and effect of oxytetracycline during the anaerobic digestion of manure from therapeutically treated calves. Process Biochemistry **41:**1637-1643.

Arya S, Williams A, Vazquez Reina S, Knapp CW, Kreft J-U, Hobman JL and Stekel DJ 2021. Towards a general model for predicting minimal metal concentrations co-selecting for antibiotic resistance plasmids. *Environmental Pollution* **275:**116602.

Baker M, Hobman JL, Dodd CER, Ramsden SJ and Stekel DJ 2016. Mathematical modelling of antimicrobial resistance in agricultural waste highlights importance of gene transfer rate. *FEMS Microbiology Ecology* **92**:fiw040.

Ivask A, Rõlova T and Kahru A 2009. A suite of recombinant luminescent bacterial strains for the quantification of bioavailable heavy metals and toxicity testing. BMC Biotechnology **9**:41.

Längin A, Alexy R, König A and Kümmerer K 2008. Deactivation and transformation products in biodegradability testing of b-lactams amoxicillin and piperacillin. Chemosphere **75:**347-354.

Malik-Sheriff RS, Glont M, Tiwari T *et al.* 2020. BioModels: 15 years of sharing computational models in life science. Nucleic Acids Research **48:**D407-D415.

**Supplementary Figure S1**: ARG and MRG collocation on slurry tank contigs

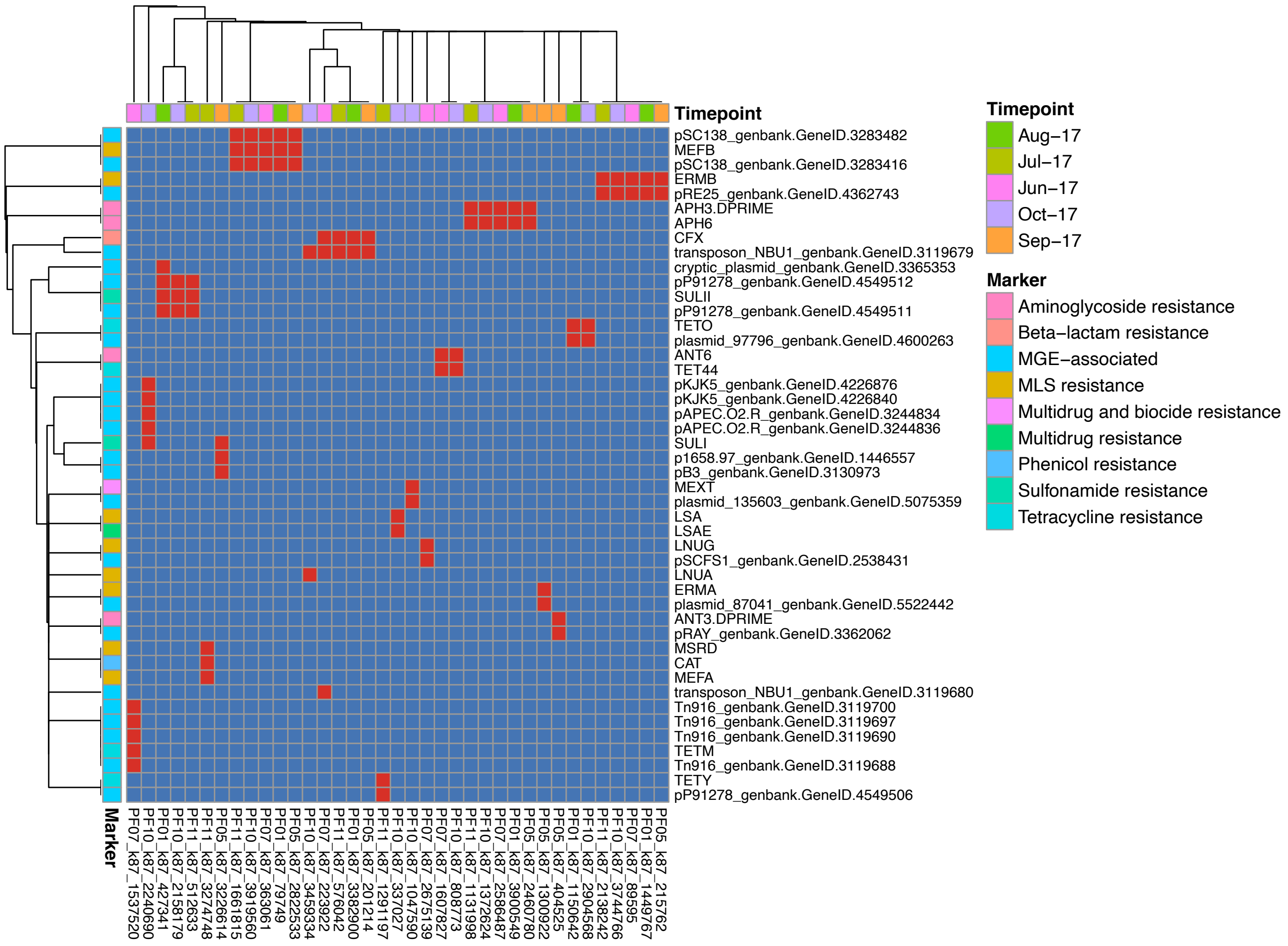

**Supplementary Figure S2**: ARG and MRG collocation on mini tank contigs

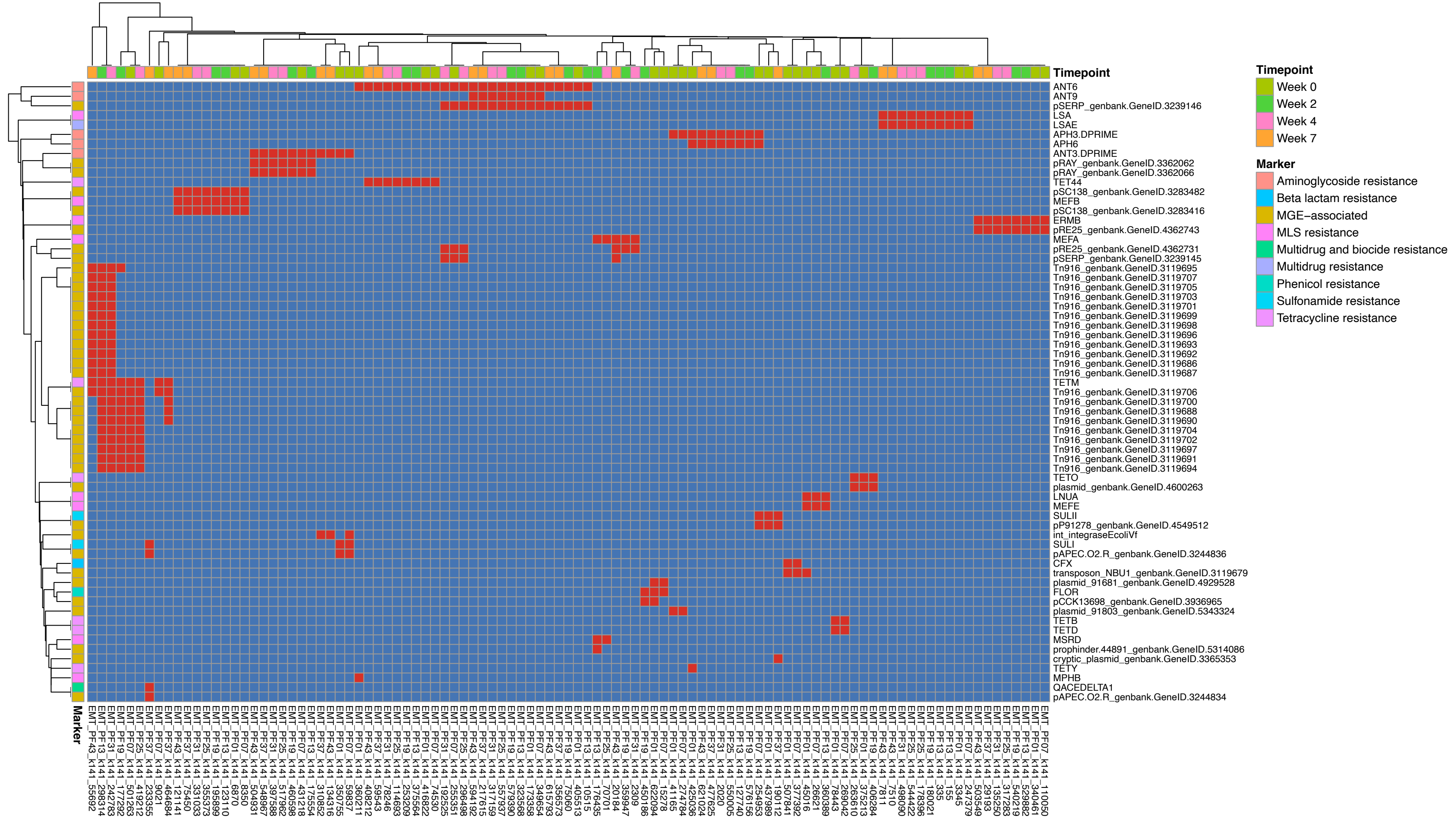

**Supplementary Figure S3:** Contig maps showing genetic context of co-located ARGs.

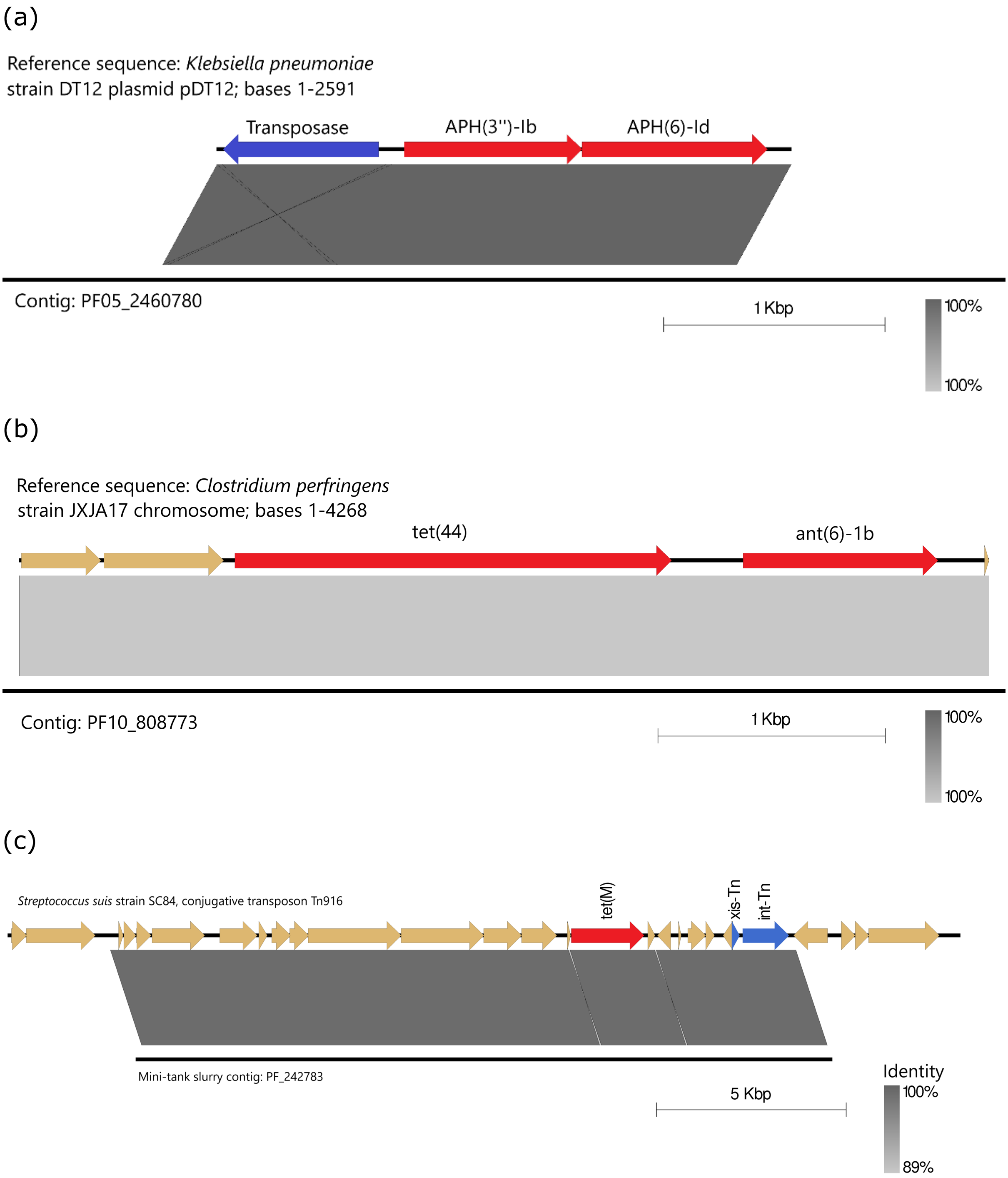

**Supplementary Figure S4:** Full Model Schematic

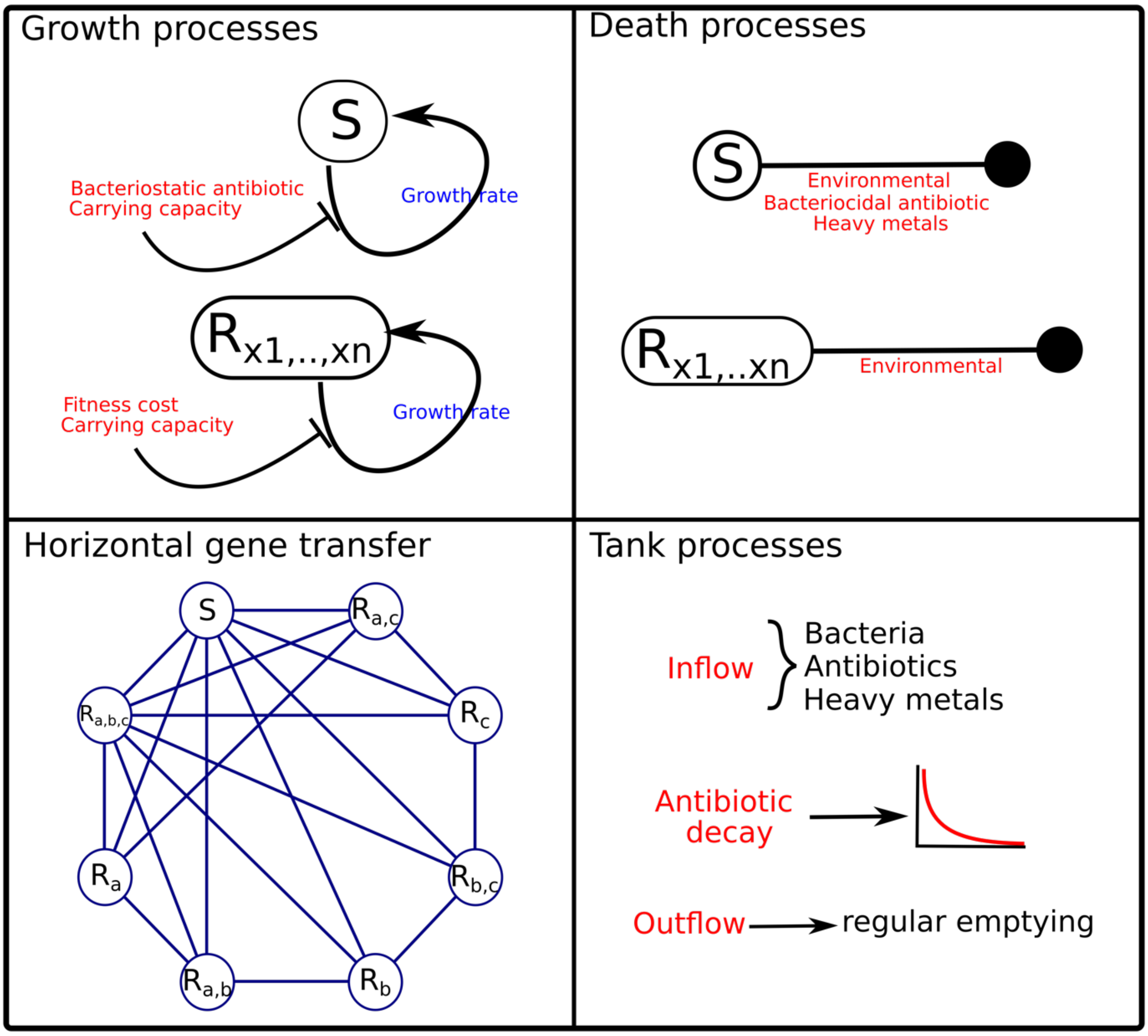

**Supplementary Figure S5:** Model fits to minitank antibiotic data

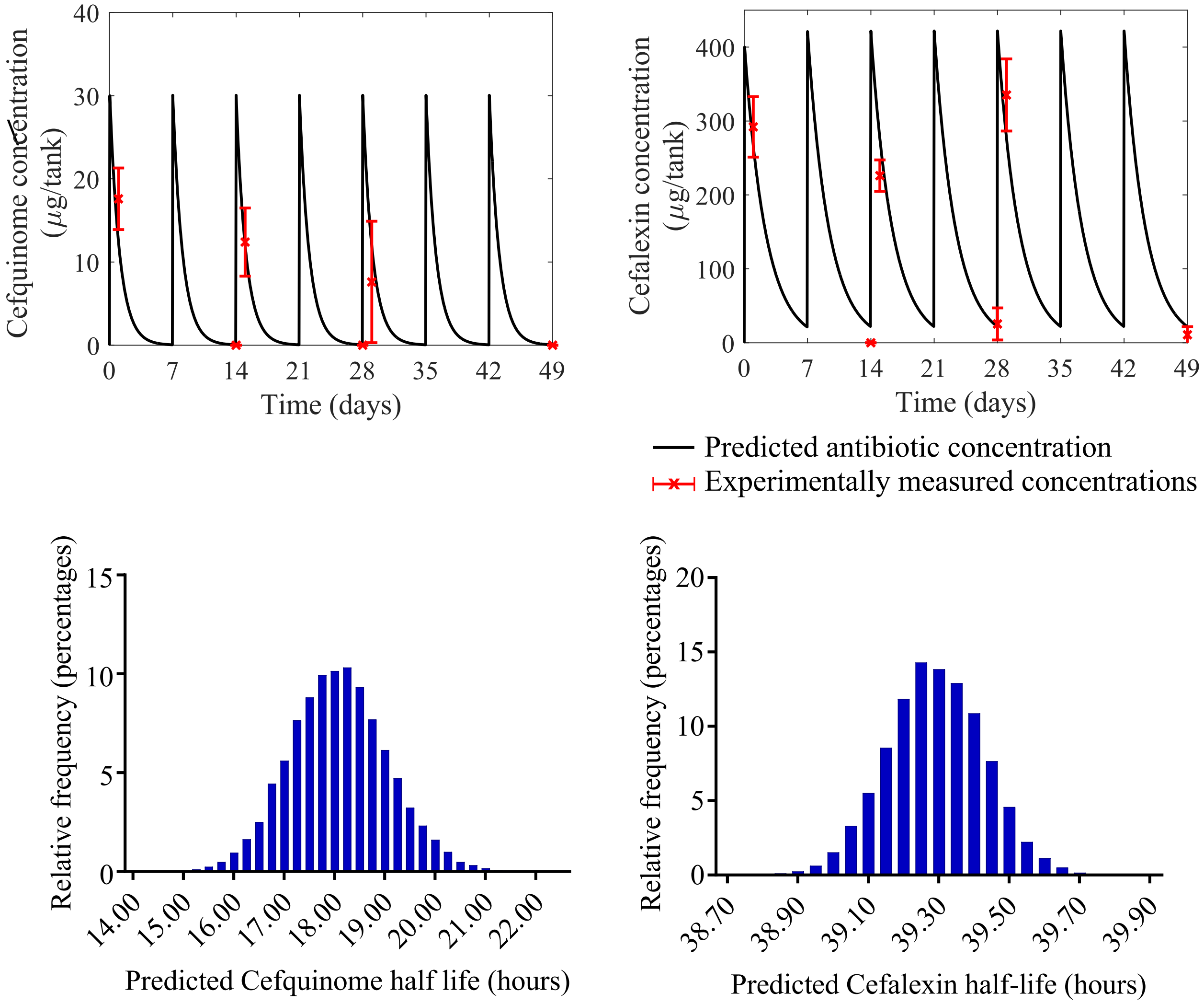

**Supplementary Figure S6:** Model fit to *E. coli* count data from mini tanks.

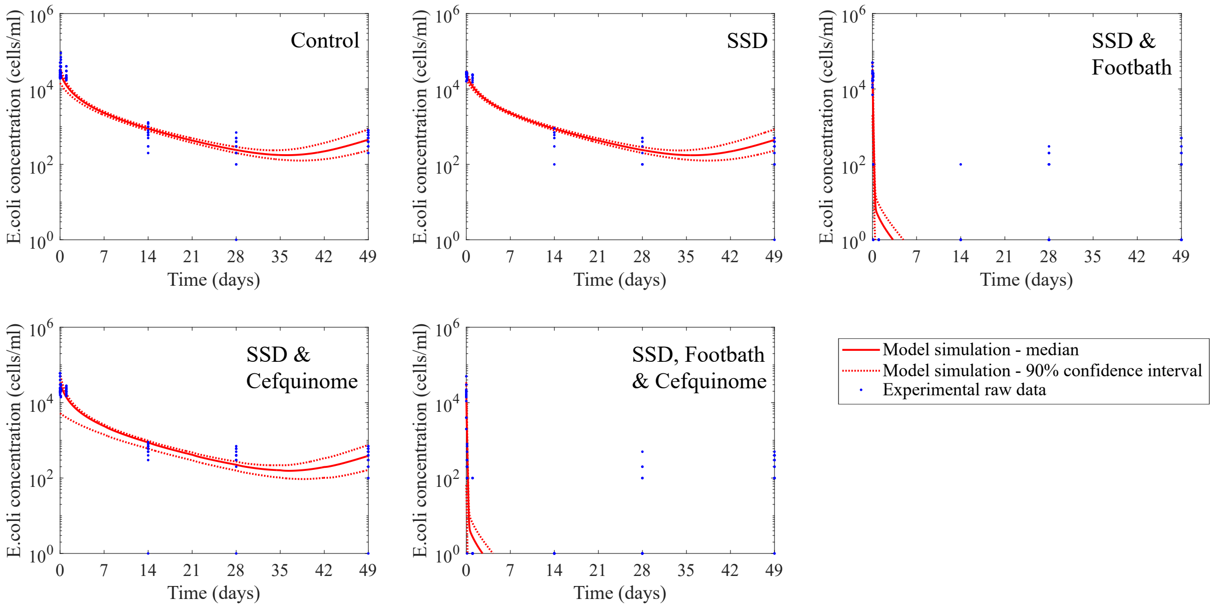

**Supplementary Figure S7:** Mini Tank Model Schematic

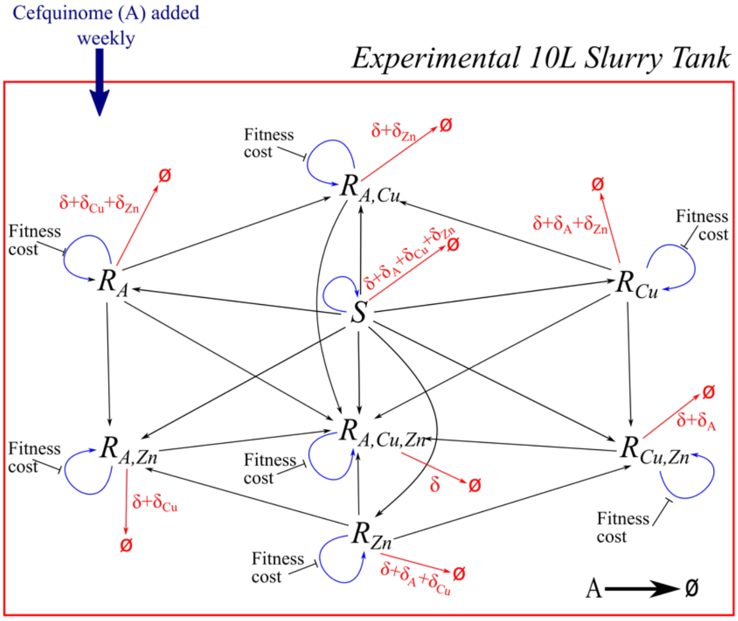


### Experimental Treatments

**Control** - Liquid slurry. Copper and Zinc at baseline levels. Cefquinome undetectable

**Control** - Liquid slurry with slurry solids added. Copper and Zinc at baseline levels. Cefquinome undetectable

**Antibiotics only**
Liquid slurry with slurry solids added. Copper and Zinc at baseline levels. Cefquinome added weekly.

**Footbath only**
Liquid slurry with slurry solids added. Additional Copper and zinc added at $t_0$. Cefquinome undetectable.

**Footbath and Cefquinome**
Liquid slurry with slurry solids added. Additional Copper and Zinc added at $t_0$. Cefquinome added weekly.

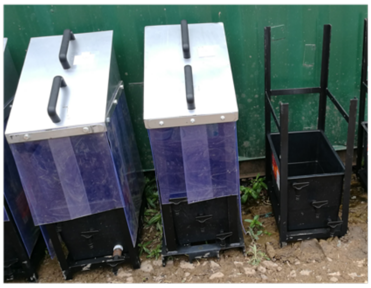

**Supplementary Figure S8:** Model simulation workflow

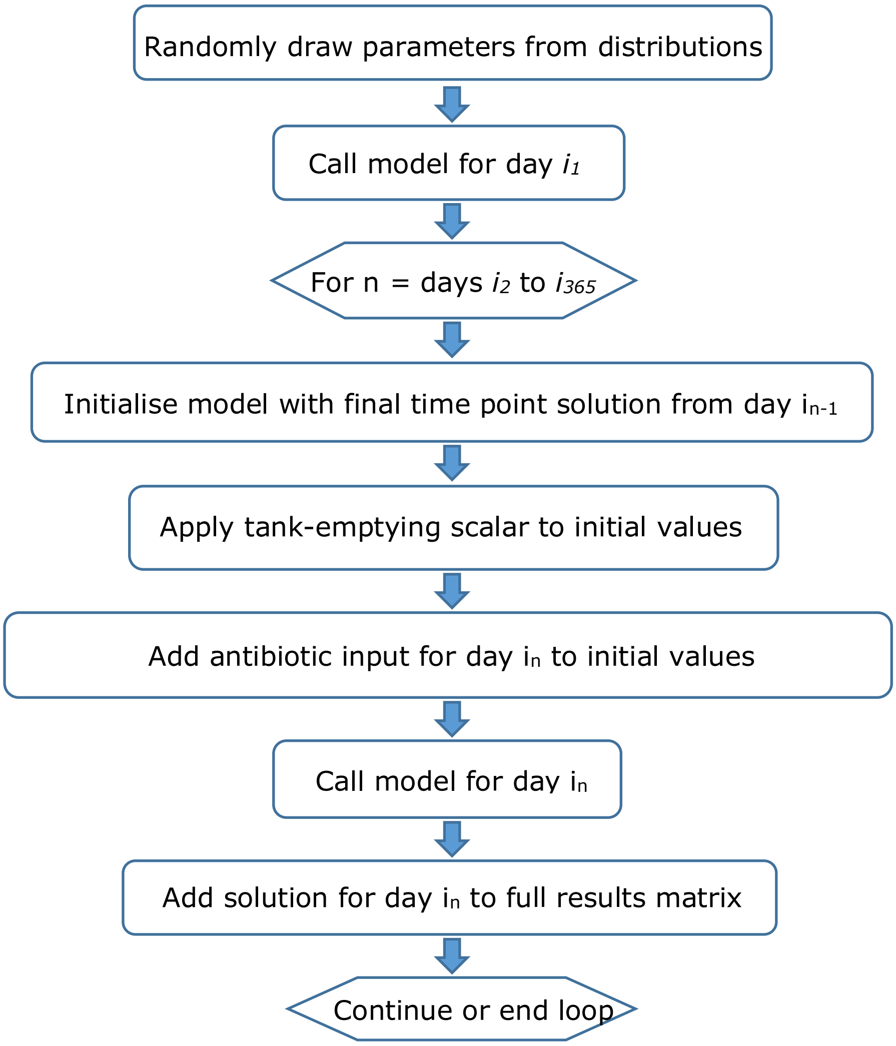

**Supplementary Figure S9:** slurry storage methods

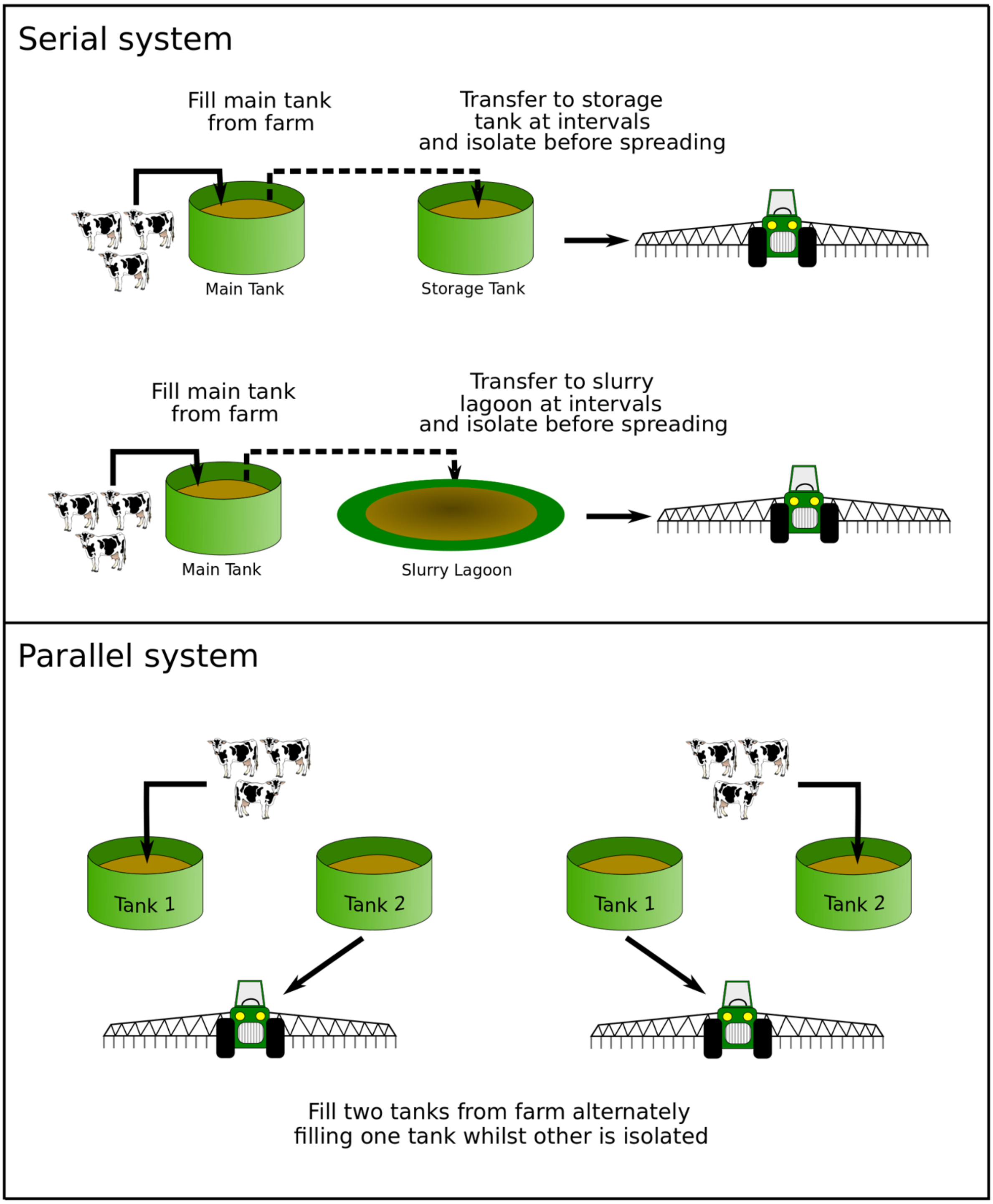

**Supplementary Figure S10:** impact on AMR of different storage methods

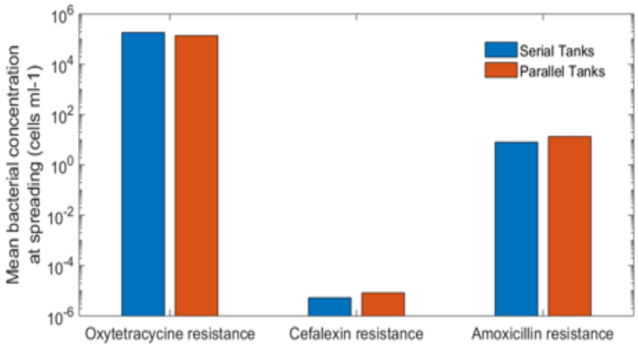
